# CflP couples magnesium availability to CovS signaling in Group B Streptococcus

**DOI:** 10.64898/2026.10.04.756536

**Authors:** Oscar J. Vazquez-Ciros, Marilia Martins Manta, Anna Riegner, Cameron W. Kenner, Yoann Le Breton, Kevin S. McIver, Nikolai Siemens, Konstantin V. Korotkov, Natalia Korotkova

## Abstract

Opportunistic pathogens must coordinate virulence-factor production with changing conditions encountered during colonization and infection. Group B *Streptococcus* (GBS), a common human opportunistic pathogen, uses the CovRS two-component system to regulate virulence, but how the CovS sensor kinase is tuned to environmental conditions remains unclear. Here, we identify CflP, a noncatalytic membrane protein of the CPBP family, as an accessory regulator of CovS. CflP associates with CovS and promotes CovS-dependent phosphorylation of the response regulator CovR, thereby repressing virulence-associated genes, including the β-hemolysin/cytolysin operon. Loss of CflP also enhances cytotoxicity toward human immune cells. CflP acts independently of Abx1, a previously identified homologous CPBP protein that promotes CovS phosphatase activity. CflP abundance is regulated post-transcriptionally by Mg²⁺ availability, linking an environmental cue to CovRS signaling. Comparative genomic analysis revealed distinct patterns of natural variation in these CovS regulators: *cflP* is predominantly maintained as an intact gene, although specific amino acid substitutions can compromise CflP abundance, whereas *abx1* is frequently disrupted by recurrent frameshifts. Together, these findings reveal an accessory regulatory layer that tunes CovS activity in response to environmental conditions and whose natural variation may contribute to strain-dependent differences in CovRS regulation. More broadly, our findings illustrate how noncatalytic membrane proteins can modulate bacterial signaling to balance virulence-factor production with adaptation to changing host environments.

**IMPORTANCE:** Group B Streptococcus (GBS) commonly colonizes the human gastrointestinal and vaginal tracts without causing disease but can cause life-threatening infections in newborns and adults. How GBS senses environmental conditions to regulate the expression of host-damaging factors remains poorly understood. We identify CflP, a membrane protein that regulates the CovRS signaling system, a central regulator of GBS virulence. CflP activates the CovS sensor kinase, promoting CovR phosphorylation and repression of genes encoding host-damaging factors. We further show that Mg²⁺ availability regulates CflP protein abundance, linking an environmental signal to CovRS activity. These findings suggest a mechanism by which GBS tunes its virulence potential in response to environmental conditions, restraining expression of host-damaging factors when nutrients are abundant while potentially permitting their induction when Mg²⁺ becomes limiting.

## INTRODUCTION

*Streptococcus agalactiae*, or Group B Streptococcus (GBS), is a Gram-positive opportunistic pathogen that colonizes and infects diverse hosts, including humans, cattle, and fish. In humans, GBS is a common member of the gastrointestinal and genitourinary microbiota, asymptomatically colonizing up to one-third of healthy adults, with the gastrointestinal tract likely serving as a reservoir for vaginal colonization in women (1–3). Despite its commensal lifestyle, GBS can transition to an invasive pathogen capable of surviving in the bloodstream, evading immune clearance, and crossing the blood-brain barrier (4–6). Approximately half of infants born to GBS-colonized mothers become colonized, and a subset develop sepsis, pneumonia, or meningitis, with mortality rates approaching 9% (7–9). Invasive GBS disease also causes substantial morbidity in nonpregnant adults (10).

The CovR/CovS two-component system is a central regulator of the transition between GBS colonization and invasive disease (11–15). In canonical two-component systems, environmental signals modulate the activity of a sensor histidine kinase (HK), which autophosphorylates in the kinase-ON state and transfers the phosphoryl group to its cognate response regulator. Many HKs also possess phosphatase activity. In the kinase-OFF state, HK removes the phosphoryl group from the cognate response regulator and resets the signaling pathway when the activating stimulus is absent (16–18). HKs typically function as homodimers, and structural and biochemical studies suggest that activation involves a conformational transition within the dimer that favors autophosphorylation (19–23).

CovS is a membrane-associated HK containing two transmembrane helices that flank an extracellular PAS-like sensing domain and a cytoplasmic catalytic region with both kinase and phosphatase activities (12, 24, 25). CovS controls the phosphorylation state of the response regulator CovR, which primarily functions as a transcriptional repressor and regulates approximately 10–15% of the GBS genome (11, 14, 15). Phosphorylation of CovR promotes its DNA-binding and repressive activity. CovR directly represses numerous virulence-associated genes, including the *cyl* operon encoding the β-hemolysin/cytolysin (β-H/C) toxin, as well as genes encoding cell-surface and secreted proteins involved in adherence and immune evasion (11, 12, 14, 26, 27). Loss of CovR results in extensive remodeling of the GBS cell surface and increased β-H/C production, contributing to enhanced cytotoxicity, inflammation, and host tissue damage (12, 15, 26, 28, 29). At the same time, CovR deficiency can increase GBS adherence to epithelial cells while impairing cellular invasion, highlighting the complex effects of CovRS regulation on host-cell interactions (12, 13, 15, 30). Thus, increased virulence-factor production can have opposing consequences for bacterial fitness: although tissue damage and inflammation may facilitate access to host tissues, dysregulated virulence-factor expression can either promote invasive disease or impair bacterial fitness depending on the host environment (12, 13, 31, 32). This context-dependent balance is consistent with a role for CovRS in regulating the transition between commensalism and invasive disease.

Although the CovR regulon has been extensively characterized using genetic and genome-wide approaches, the environmental signals that activate CovS in GBS remain poorly defined. Moreover, how CovS transitions between its kinase and phosphatase states is not understood. In addition to the core HK and response regulator, some two-component systems contain accessory proteins that directly interact with HKs and stabilize specific signaling conformations (18, 33–36). By prolonging or biasing a particular HK state, these proteins can provide an additional layer of control that couples two-component signaling to cellular physiology. Accessory regulators can themselves be controlled by ligand binding, post-translational modification, or regulated expression, allowing HK activity to be tuned according to metabolic state, environmental stress, or developmental stage (35).

In GBS, the membrane protein Abx1 has been identified as an accessory regulator of CovS (24). Abx1 interacts directly with CovS and promotes the phosphatase-competent state, resulting in reduced CovR phosphorylation and derepression of CovR-regulated genes (24). However, the signals controlling Abx1 activity and the mechanisms that promote the opposing kinase state of CovS remain unknown. Understanding these mechanisms is particularly important because inappropriate activation of β-H/C and other virulence factors can cause substantial host damage and may compromise bacterial fitness. Abx1 belongs to the CPBP (CAAX protease and bacteriocin-processing) protein family, a large and widely distributed family of bacterial membrane proteins (37). CPBPs were originally identified as bacteriocin-processing proteins because some are encoded within bacteriocin-associated loci and contribute to producer-cell resistance (38–41). However, direct biochemical evidence for proteolytic activity is lacking for many family members. Consistent with a noncatalytic function, the predicted catalytic residues of Abx1 are dispensable for its regulation of CovS (24), suggesting that CPBPs can function as protein-interaction scaffolds rather than proteases. Whether other GBS CPBPs similarly regulate CovS or other membrane-associated proteins remains unknown.

Here, we identify CflP, a previously uncharacterized CPBP-family membrane protein and the closest homolog of Abx1 in GBS, as a positive regulator of CovS. CflP directly associates with CovS and promotes CovR phosphorylation, consistent with stabilization of the CovS kinase state. We further show that CflP abundance is regulated by environmental Mg²⁺ availability, providing a mechanism that links a physiologically relevant environmental signal to CovRS activity and β-H/C production. Together, our findings reveal an accessory-protein mechanism for tuning the CovS kinase/phosphatase switch and suggest how GBS can balance virulence and commensalism in response to changing host environments.

## RESULTS

### CflP restrains β-H/C production

Expression of the β-hemolysin/cytolysin (β-H/C), encoded by the *cyl* operon, is strongly influenced by nutrient availability. Because many GBS strains produce little or no detectable pigment in nutrient-rich media, β-H/C production is commonly assessed on Granada medium, where β-H/C-producing strains form characteristic red-orange colonies (42, 43). Granada medium is a complex starch- and serum-based medium in which folate inhibitors stimulate β-H/C expression through an incompletely understood mechanism. To provide a defined system for identifying regulators of β-H/C under nutrient-limited conditions, we developed a chemically defined medium (CDM) containing low concentrations of nicotinamide, an essential vitamin that modulates β-H/C production. Because vitamin availability in the female genital tract can vary with the local environment and is influenced by Lactobacillus species (44, 45), this condition may better approximate aspects of the nutrient environment encountered by GBS during colonization.

Using this CDM, we screened a transposon mutant library of the serotype V GBS strain CJB111 for mutants with altered pigmentation. A transposon insertion in ID870_04370, which we subsequently named *cflP*, resulted in markedly increased pigmentation. To validate this phenotype, we generated in-frame *cflP* deletions in CJB111 and the serotype Ia strain A909, CJB111Δ*cflP* and A909Δ*cflP*, respectively. Because β-H/C is a poorly soluble, pigmented ornithine-rhamnolipid toxin that remains largely cell-associated and can also be released in membrane vesicles (46, 47), we quantified β-H/C in both cell pellets and cell-free supernatants of GBS strains grown in CDM. Both A909Δ*cflP* and CJB111Δ*cflP* produced significantly more cell-associated β-H/C (Fig. 1A). Deletion of *cflP* increased supernatant β-H/C 4.6-fold in CJB111 but only 1.2-fold in A909, a 3.8-fold difference in effect between backgrounds (P < 0.0001, strain × genotype interaction, two-way ANOVA), whereas the two wild-type strains did not differ in β-H/C secretion (Fig. 1A). Complementation with a high-copy-number plasmid carrying WT *cflP* restored β-H/C production to very low levels in both strains (Fig. 1A), consistent with increased CflP abundance strongly repressing β-H/C production. In this construct, *cflP* is expressed from the constitutive P_tet_/P_cat_ promoter of pDC123 vector (48). In addition, A909Δ*cflP* and CJB111Δ*cflP* produced substantially more pigment than their respective WT strains on CDM agar (Fig. 1B). CJB111Δ*cflP* was also more pigmented than the WT and complemented strains on Granada agar (Fig. S1A). These data identify CflP as a negative regulator of β-H/C production.

**Figure 1.**
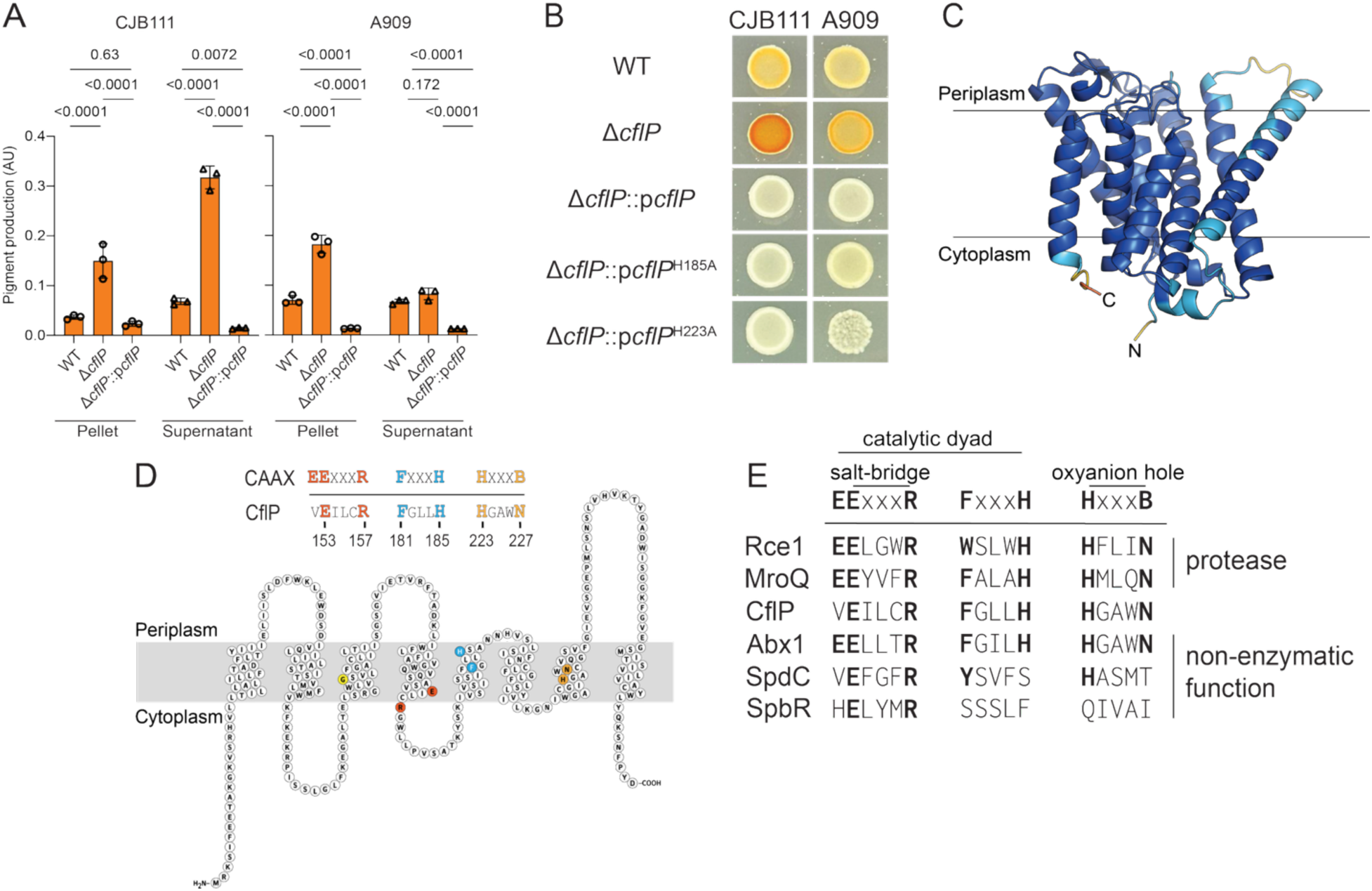
CflP is a noncatalytic protein scaffold that acts as a repressor of β-H/C production. (**A**) Quantification of pigmented ornithine rhamnolipid (β-H/C) extracted from cell pellets and cell-free supernatants of GBS strains grown in CDM-HighMg at 37 °C for 24 h. β-H/C content was measured at 522 nm and normalized to the final OD_600_. Columns and error bars represent the mean and SD, respectively (n = 3 biologically independent replicates). *P* values were determined by two-way ANOVA with Tukey’s multiple-comparison test. (**B**) Pigmentation phenotypes of A909, CJB111, and their respective isogenic *cflP* deletion mutants complemented with either WT *cflP* or *cflP* variants carrying substitutions in conserved histidine residues predicted to contribute to CPBP catalytic activity. Images were taken after growth on CDM agar at 37 °C for 24 h. Experiments were performed independently three times with similar results; a representative image is shown. (**C**) AlphaFold2-predicted structure of CflP. Colors indicate the predicted local confidence score (pLDDT), with dark blue representing high-confidence regions and yellow/orange representing lower-confidence regions. (**D**) Schematic representation of the predicted CflP membrane topology. Conserved residues within putative CPBP catalytic motifs are indicated. G106, a residue implicated in CflP folding, is highlighted in yellow. (**E**) Structure-based sequence alignment of conserved CPBP motifs (EExxxR, FxxxH, and HxxxB) of *Methanococcus maripaludis* Rce1 (PDB 4CAD), *S. aureus* MroQ (SAOUHSC_02115), GBS A909 CflP (SAK_1097) and Abx1 (SAK_1496), *S. aureus* SpdC (PDB 6U0O, SAOUHSC_02611) and SpbR (SAOUHSC_00965). Conserved residues are shown in bold.

### CflP represses β-H/C independently of conserved CPBP catalytic residues

The *cflP* gene is transcribed from its own promoter, and previous work identified it as a direct target of CovR (11, 14, 15). CflP is an integral membrane protein predicted to contain eight transmembrane α-helices, and it belongs to the CPBP (CAAX protease and bacteriocin-processing) family (Fig. 1C and D). It shares 41% amino acid sequence identity with the GBS CPBP protein Abx1 (24). The CPBP superfamily is defined by three conserved sequence motifs, EExxxR (motif 1), FxxxH (motif 2) and HxxxB (motif 3, where B is aspartate or asparagine) (49). CflP retains the histidines of CPBP motifs 2 and 3 (H185 and H223) and the motif 3 asparagine (N227). In the crystal structure of CAAX protease Rce1 (50), motif 3 histidine and asparagine form the oxyanion hole, whereas motif 2 histidine forms the catalytic dyad together with the first glutamate of motif 1 (Fig. 1E). At the position of that catalytic glutamate, however, CflP carries a valine (V152) (Fig. 1D and E). The second glutamate of motif 1 is buried and forms a salt bridge with the motif 1 arginine. This salt bridge is likely required for protein stability and is present in the experimental and predicted structures of catalytic and noncatalytic CPBP proteins (Fig. 1E and S1B).

To determine whether the predicted catalytic residues contribute to CflP-dependent regulation of β-H/C, we introduced H185A or H223A substitutions into *cflP* expressed from the p*cflP* plasmid. Expression of either mutant protein in A909Δ*cflP* or CJB111Δ*cflP* resulted in β-H/C levels comparable to those observed with WT *cflP* complementation, with β-H/C remaining largely undetectable in CDM (Fig. 1B). Thus, neither conserved histidine is required for CflP-mediated repression of β-H/C production, indicating that CflP functions independently of the predicted catalytic activity of CPBP proteins and is more consistent with a nonenzymatic regulatory or scaffolding role.

### CflP links Mg²⁺ availability to CovRS-dependent regulation of β-H/C

Because deletion of *cflP* increased β-H/C production in both A909 and CJB111, we selected A909 as the primary strain background for subsequent mechanistic studies of CflP-dependent CovRS regulation. Selected findings were subsequently examined in CJB111 to assess whether CflP function was conserved across GBS strain backgrounds.

Previous studies showed that the CovS homolog in Group A Streptococcus (GAS) responds to Mg²⁺ and sub-inhibitory concentrations of the antimicrobial peptide LL-37 (51–55). However, whether these signals regulate CovS activity in GBS remains unknown. To determine whether Mg²⁺ availability affects CovS-dependent β-H/C production, we compared A909 grown in CDM containing either high Mg²⁺ (CDM-HighMg; 28.4 mM Mg²⁺) or low Mg²⁺ (CDM-LowMg; 7 mM Mg²⁺). Because Mg²⁺ concentrations below 7 mM reduced GBS growth under these conditions, we used 7 mM Mg²⁺ as the low-Mg²⁺ condition for subsequent experiments. β-H/C production was significantly increased under low-Mg²⁺ conditions (Fig. 2A). In contrast, Mg²⁺ availability had no significant effect on β-H/C production in either A909Δ*cflP* or A909Δ*covS* (Fig. 2A), suggesting that Mg²⁺-dependent regulation of β-H/C requires both CflP and CovS.

**Figure 2.**
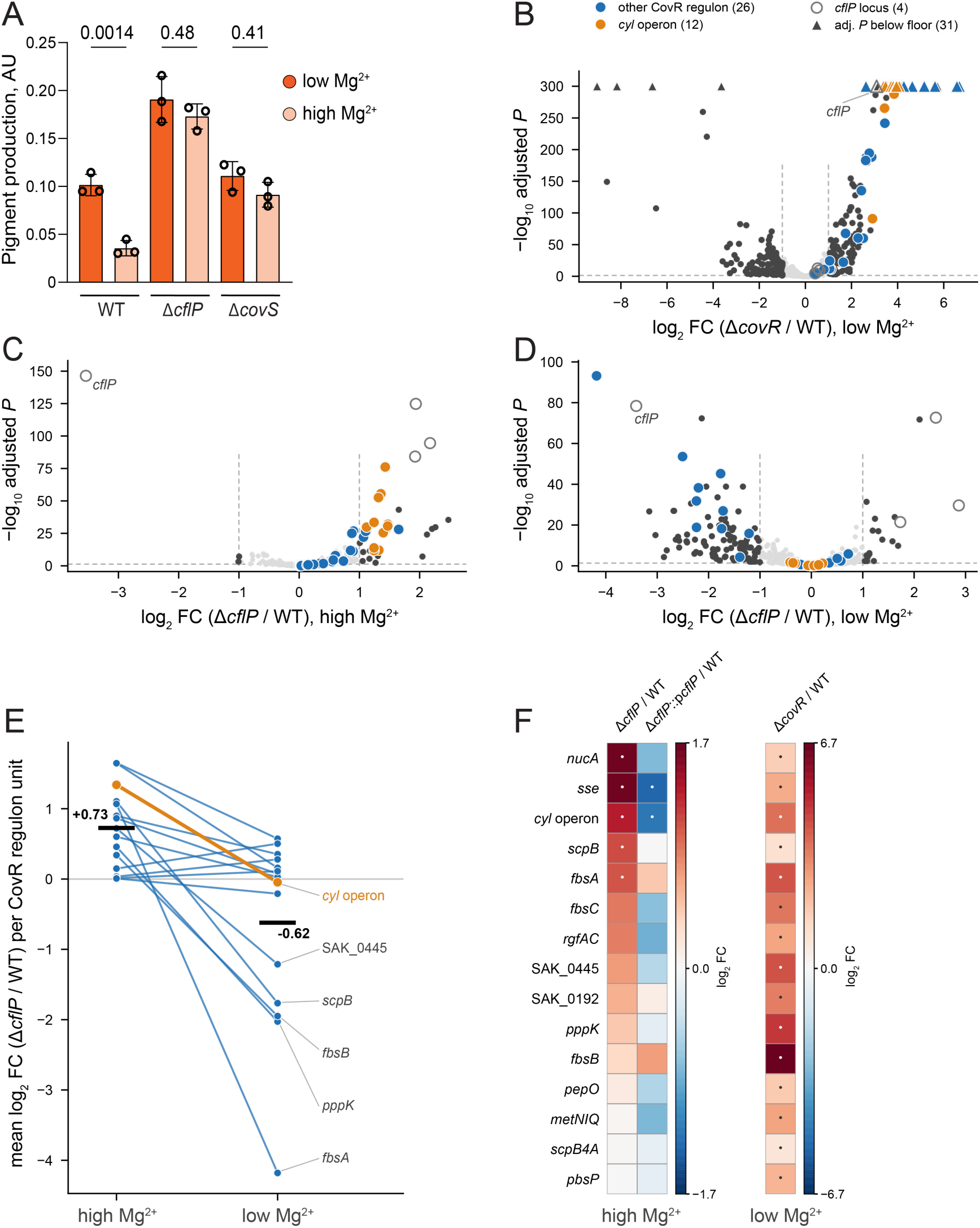
CflP-dependent transcriptional regulation is influenced by Mg²⁺ availability. (**A**) Quantification of pigmented ornithine rhamnolipid (β-H/C) extracted from cell pellets of GBS strains grown in CDM-HighMg or CDM-LowMg at 37 °C for 24 h. β-H/C content was measured at 522 nm and normalized to the final OD₆₀₀. Columns and error bars represent the mean and SD, respectively (*n* = 3 biologically independent replicates). *P* values were determined by one-way ANOVA with Tukey’s multiple-comparison test. (**B**) Volcano plot showing differentially expressed genes in A909Δ*covR* relative to A909 grown in CDM-LowMg. For 31 genes adjusted *P* falls below the double-precision floor; all are drawn as triangles at the top of the axis. 21 of the 31 are CovR regulon genes including *cflP* (nine of them *cyl* operon) and 10 lie outside the regulon. Genes of the *cyl* operon are orange and other CovR regulon genes blue; the four genes of the *cflP* locus are open circles. Differentially expressed genes were defined as those with |log₂ fold change| > 1 and adjusted *P* < 0.05 (dashed lines). (**C** and **D**) The same plots for A909Δ*cflP* relative to A909 grown in CDM-HighMg (**C**) or CDM-LowMg (**D**). (**E**) Mean log₂ fold change in A909Δ*cflP* relative to A909 for each of the 15 transcriptional units of the CovR regulon, at the two magnesium concentrations. Black bars are the means across units. The paired difference across units is significant (Wilcoxon *P* = 1.5e-03). (**F**) Mean log₂ fold change per regulon unit in Δ*cflP* and in Δ*cflP*::p*cflP*, both relative to A909 in CDM-HighMg, with A909Δ*covR* relative to A909 in CDM-LowMg as a reference on its own color scale. The units are ordered by their Δ*cflP* response. Dots mark units containing at least one gene meeting both thresholds, in all three columns. Every unit meets both thresholds in the Δ*covR* column, which is what establishes the set as CovR-controlled. The complete RNA-seq data underlying **B**–**F** are provided in Dataset S1.

Previous studies identified *cflP* as a CovR-regulated gene under rich media conditions (11, 14, 15, 24). We first compared the transcriptomes of mid-log-phase A909 WT and A909Δ*covR* grown in CDM-LowMg at 37 °C. Loss of *covR* altered the expression of 253 genes (|log₂ fold change| > 1, adjusted *P* < 0.05). All 38 genes of the CovR direct regulon were significantly derepressed, 36 of them beyond the fold-change threshold (Fig. 2B, Dataset S1). The observed changes agree with the published A909 Δ*covR* microarray (*r* = 0.93 across 37 genes with probes, Fig. S2A) (15). Consistent with published studies, *cflP* was itself derepressed in Δ*covR*.

To further define the role of CflP in CovRS-dependent gene regulation, we compared the transcriptomes of mid-log-phase A909 WT, A909Δ*cflP*, and A909Δ*cflP*::p*cflP* grown in CDM-HighMg at 37 °C. Deletion of *cflP* resulted in differential expression of 36 genes relative to WT (|log₂ fold change| > 1; adjusted *P* < 0.05), 33 of which were expressed at higher levels in the mutant (Fig. 2C). Among the differentially expressed genes were the *cyl* operon, which encodes the β-H/C biosynthesis machinery, and several established members of the CovRS regulon (11, 14, 15, 24), including *nucA*, encoding secreted nuclease (56); *scpB*, encoding the C5a peptidase ScpB (57); and *sse*, encoding the secreted esterase Sse (58). Expression of several additional virulence-associated genes was also increased in A909Δ*cflP*, including *fbsA*, encoding a fibrinogen-binding protein (59, 60); (Fig. 2F).

Complementation of A909Δ*cflP* with *cflP* altered the expression of 169 genes relative to the mutant (Fig. S2B and S2C). Complementation reversed the effect of the deletion on the CovR regulon. Of the 15 transcriptional units of the regulon, 14 changed in the direction opposite to their response to *cflP* deletion, including *cyl* operon (Fig. 2F). These results support a role for CflP in regulating β-H/C production and a subset of CovRS-controlled genes.

We next asked whether Mg²⁺ availability influences CflP-dependent transcription. In CDM-LowMg, deletion of *cflP* altered expression of 131 genes, 112 of which were expressed at lower levels in the mutant (Fig. 2D). The response differed markedly from that observed in CDM-HighMg: of the 16 genes differentially expressed under both conditions, four were affected in *cis* by the *cflP* deletion, whereas the remaining 12 showed opposite responses to loss of CflP, with increased expression under high-Mg²⁺ and decreased expression under low-Mg²⁺. Consistent with this divergence, genome-wide fold changes under high- and low-Mg²⁺ conditions were weakly negatively correlated (*r* = −0.26; Fig. S2D). Notably, *cflP* deletion under low-Mg²⁺ conditions did not derepress the *cyl* operon or other CovR-regulated genes that were induced in the mutant under high-Mg²⁺ (Fig. 2E). This difference was not attributable to reduced sensitivity of the low-Mg experiment: Δ*covR*, analyzed in the same experiment with the same WT samples, strongly derepressed all 15 CovR-regulated transcriptional units, whereas Δ*cflP* did not (Fig. S2E). Thus, the transcriptional consequences of *cflP* deletion depend strongly on Mg²⁺ availability, with CflP exerting a more pronounced effect on the CovRS regulon under high-Mg²⁺ conditions. Together with the Mg²⁺-dependent β-H/C phenotype, these findings support a role for CflP in coupling Mg²⁺ availability to CovRS signaling.

### CflP acts through CovRS to regulate β-H/C and cell envelope remodeling

To determine whether CflP regulates β-H/C through the CovRS pathway and whether this activity depends on Abx1, we examined β-H/C production and hemolysis in A909 single and double mutants lacking *cflP*, *covR*, *covS*, and/or *abx1*. β-H/C production was assessed on CDM-LowMg, THY, and BHI agar and in CDM-LowMg broth (Fig. 3A and C). Δ*covR* was strongly hyperpigmented under all conditions tested, whereas Δ*covS* showed an intermediate pigmentation phenotype (Fig. 3A). Importantly, deletion of *cflP* did not further increase β-H/C production in either Δ*covR* or Δ*covS*, and overexpression of *cflP* did not suppress β-H/C production in Δ*covR*Δ*cflP* or Δ*covS*Δ*cflP* (Fig. 3A and C). Similarly, CflP overexpression suppressed hemolysis in Δ*cflP* and Δ*abx1*Δ*cflP* but had no effect in Δ*covS*Δ*cflP* or Δ*covR*Δ*cflP* (Fig. 3A and C). These results place CflP upstream of CovR-dependent repression of the *cyl* operon and demonstrate that both CovS and CovR are required for CflP-mediated regulation, whereas Abx1 is dispensable.

**Figure 3.**
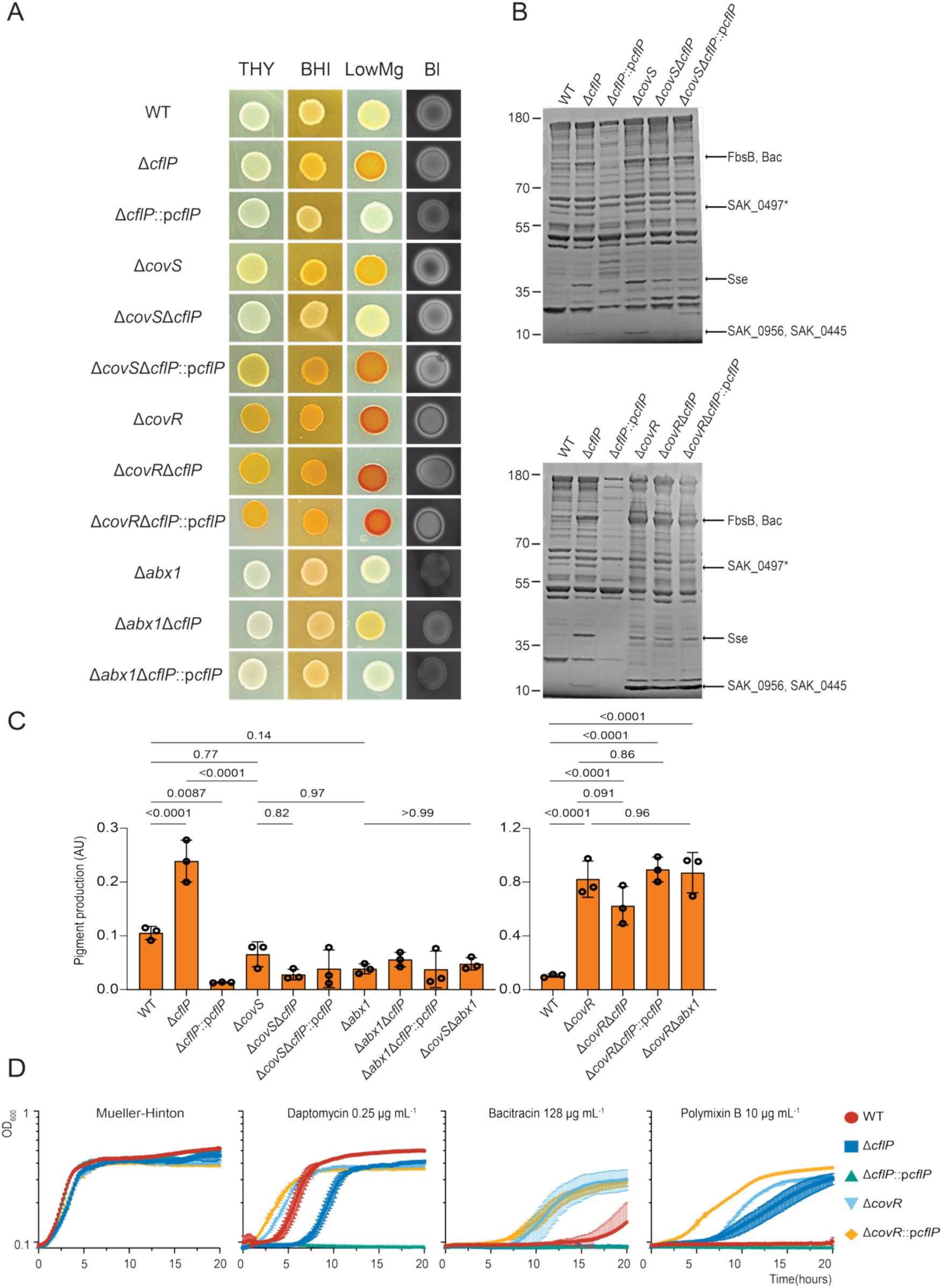
CflP regulates β-H/C production, secreted protein abundance, and antibiotic susceptibility through CovRS. (**A**) Pigmentation and hemolysis phenotypes of A909, isogenic mutants lacking *cflP, covR, covS,* or *abx1*, and complemented strains. Images were taken after growth on THY, BHI, or blood (Bl) agar at 37 °C for 24 h, or on CDM-LowMg agar under anaerobic conditions in a GasPak EZ chamber at 40 °C for 24 h. Experiments were performed independently three times with similar results; a representative image is shown. (**B**) SDS-PAGE analysis of proteins in cell-free culture supernatants of GBS strains grown in THY broth at 37 °C for 24 h. Proteins were separated on a 4–12% SurePAGE Bis-Tris gel in MOPS buffer and visualized with SYPRO Ruby. Bands indicated by arrows were excised from the Δ*cflP* sample and identified by LC-MS/MS as described in Materials and Methods. SAK_0497 is frameshifted in A909; the identified peptides map to the first reading frame. Experiments were performed independently three times with similar results; representative gels are shown. (**C**) Quantification of pigmented ornithine rhamnolipid (β-H/C) extracted from cell pellets of GBS strains grown in CDM-LowMg at 37 °C for 24 h. β-H/C content was measured at 522 nm and normalized to the final OD_600_. Columns and error bars represent the mean and SD, respectively (*n* = 3 biologically independent replicates). *P* values were determined by one-way ANOVA with Tukey’s multiple-comparison test. (**D**) Growth curves of A909, isogenic mutants lacking *cflP* or *covR*, and complemented strains in Mueller-Hinton broth without antibiotics or supplemented with daptomycin (0.25 μg mL⁻¹), bacitracin (128 μg mL⁻¹), or polymyxin B (10 μg mL⁻¹). Curves represent the mean ± SD (*n* = 3 biologically independent replicates). Dose-response growth curves for each antibiotic are shown in Fig. S4.

Unexpectedly, deletion of *abx1* alone did not alter β-H/C production in A909 under the conditions tested, despite previous reports implicating Abx1 in CovS regulation (Fig. 3A and C) (37). Similarly, *abx1* deletion did not alter the pigmentation phenotypes of Δ*covS* or Δ*covR* (Fig. 3A and C). However, Δ*abx1* was nonhemolytic on blood agar (Fig. 3A), and plasmid-mediated expression of *abx1* increased pigmentation in both A909Δ*abx1* and CJB111 (Fig. S3A and B). These findings suggest that Abx1 activity depends strongly on strain and growth conditions and may contribute to CovS regulation when Abx1 is sufficiently abundant or active.

We next asked whether CflP influences the CovRS-dependent remodeling of the secreted proteome. Secreted proteins from A909 WT and mutants lacking CflP, CovR, or CovS were analyzed by SDS-PAGE following growth in THY broth. The secretome of Δ*cflP* closely resembled that of Δ*covS* (Fig. 3B, top panel). LC-MS/MS identified six proteins more abundant in Δ*cflP*: Sse, FbsB, Bac, and the hypothetical secreted proteins SAK_0445 and SAK_0956. The increased abundance of Sse, FbsB, and SAK_0445 was consistent with elevated transcription of their corresponding genes observed by RNA-seq (Fig. 2F). SAK_0497, encoding an S8 family serine peptidase, is annotated as a pseudogene in A909. Its reading frame is interrupted by a frameshift within a homopolymeric tract. The identified peptides correspond to the first reading frame, which is transcribed and elevated in A909Δ*cflP* and reversed upon complementation (Dataset S1). Complementation of Δ*cflP* with *cflP* reduced the abundance of these proteins, whereas deletion or overexpression of *cflP* in the Δ*covS* background had little effect on the secreted protein profile (Fig. 3B, top panel). Similarly, the secretomes of Δ*covR*, Δ*covR*Δ*cflP*, and Δ*covR*Δ*cflP*::p*cflP* were comparable (Fig. 3B, bottom panel), further demonstrating that CovR and CovS are required for CflP-dependent regulation. Immunoblotting further confirmed that CflP negatively regulates Bac abundance in A909 but not in Δ*covR* or Δ*covS* (Fig. S3C).

Because CovRS-dependent regulation extensively remodels the GBS cell envelope and affects resistance to host antimicrobial defenses, we next examined susceptibility to antibiotics that perturb cell-envelope integrity. Compared with WT, Δ*covR* was more resistant to polymyxin B and bacitracin, whereas Δ*cflP* showed increased resistance to polymyxin B but increased sensitivity to daptomycin and bacitracin (Fig. 3D and Fig. S4). Complementation of Δ*cflP* restored susceptibility to all three antimicrobials (Fig. 3D and Fig. S4). In contrast, CflP overexpression did not alter the antimicrobial susceptibility of Δ*covR*, consistent with CflP acting through CovR (Fig. 3D and Fig. S4). Together, these findings demonstrate that CflP influences multiple aspects of GBS cell-envelope physiology—including secretion of virulence-associated proteins and resistance to membrane-active and cell-wall-targeting antimicrobials—and that these effects require an intact CovRS signaling pathway.

### CflP promotes CovR phosphorylation in a CovS-dependent manner

To determine how CflP regulates CovRS signaling and whether its activity depends on Abx1, we examined the phosphorylation state of CovR in A909 WT and mutants lacking *cflP*, *covS*, *abx1*, or *covR*. Cell lysates from mid-log-phase cultures grown in CDM-LowMg or THY were analyzed by Phos-Tag electrophoresis followed by immunoblotting with CovR-specific antibodies. Conventional SDS-PAGE immunoblotting showed comparable CovR abundance and electrophoretic mobility among all strains except Δ*covR* (Fig. S5B, and D) and Ponceau S staining confirmed equivalent protein loading and transfer (Fig. S5A, B, C and D). Phos-Tag analysis revealed two CovR species in both CDM- and THY-grown WT cells (Fig. 4B and C). The slower-migrating species corresponded to CovR∼P because these bands were not present in a sample treated with alkaline phosphatase or heated at 90 °C for 10 min (Fig. 4B and C, lanes 1 and 3). Thus, CovR was predominantly unphosphorylated under the conditions tested.

**Figure 4.**
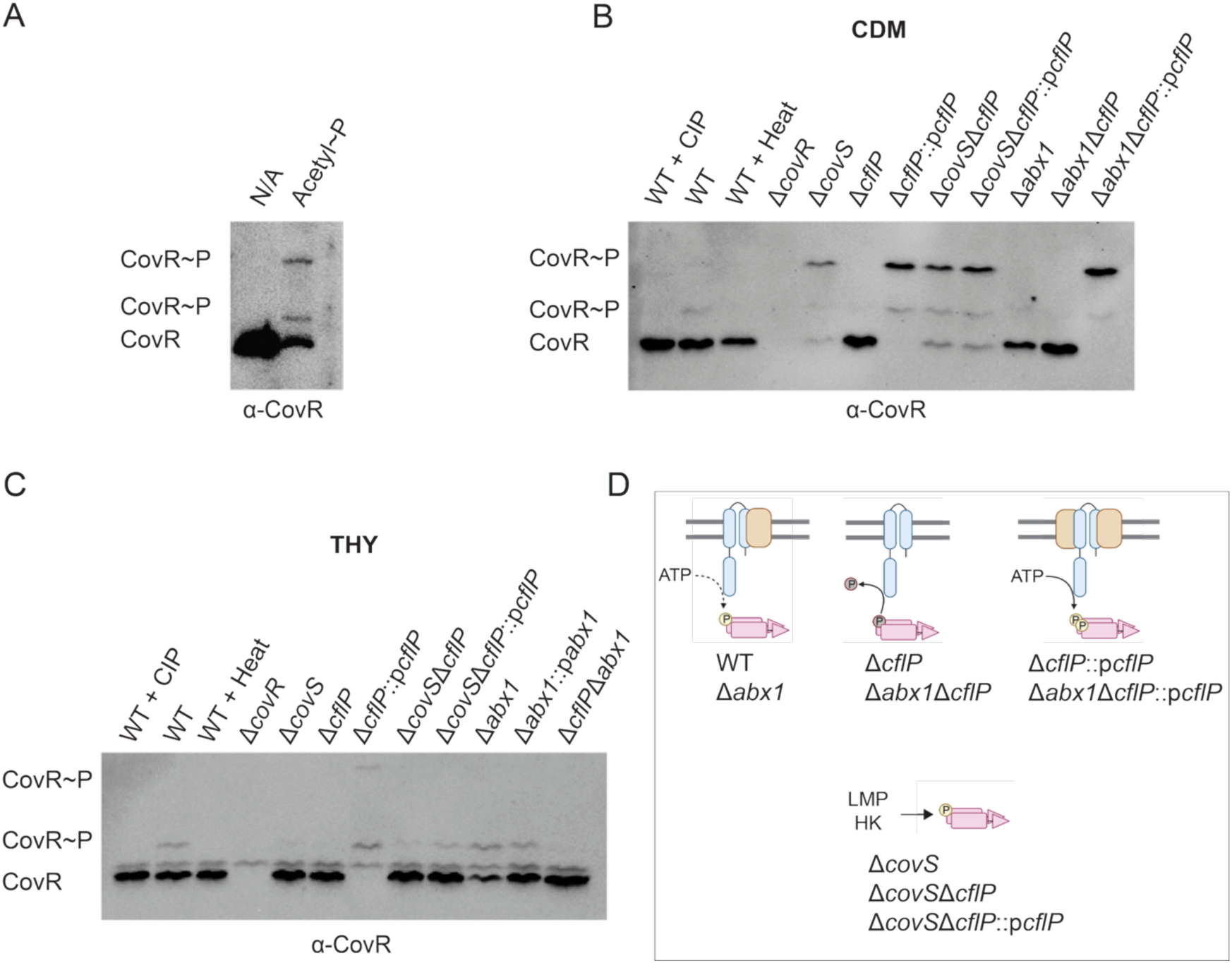
CflP promotes CovR phosphorylation through the sensor histidine kinase CovS. (**A**) Representative immunoblot of Phos-tag SDS-PAGE showing unphosphorylated CovR and phosphorylated CovR (CovR∼P). The migration positions of CovR∼P were established by incubating purified His-tagged CovR with 20 mM acetyl phosphate (acetyl∼P), which generated two distinct CovR∼P bands. (**B** and **C**) Representative Phos-tag immunoblots of CovR in cell extracts from GBS strains grown to mid-log phase in CDM-LowMg (**B**) or THY (**C**) at 37 °C. Sample preparation, Phos-tag SDS-PAGE, and immunoblotting with anti-CovR antibodies were performed as described in Materials and Methods. Experiments were performed independently twice for **A** and three times for **B** and **C**; representative images are shown. Uncropped immunoblots and Ponceau S-stained membranes are shown in Fig. S5. (**D**) Proposed models for CovR phosphorylation by CovS in the presence or absence of CflP. LMP, low-molecular-weight phosphodonor; HK, histidine kinase.

Deletion of *cflP* eliminated detectable CovR∼P, whereas complementation with plasmid-expressed *cflP* restored and strongly increased CovR phosphorylation (Fig. 4B and C). These results identify CflP as a positive regulator of CovR phosphorylation. Interestingly, two slowly migrating CovR species were observed upon CflP overexpression in CDM. Although dual phosphorylation of the GAS CovR homolog has previously been attributed to phosphorylation at an additional threonine residue by an eukaryotic-like Ser/Thr kinase (27, 61), recombinant CovR phosphorylated with acetyl phosphate also produced two distinct species *in vitro* (Fig. 4A), indicating that the two bands do not necessarily reflect phosphorylation at distinct sites by different enzymes.

The effect of CflP on CovR phosphorylation required CovS. In THY-grown Δ*covS*, CovR phosphorylation was reduced relative to WT, whereas in CDM-grown Δ*covS*, CovR phosphorylation increased (Fig. 4B and C). The elevated phosphorylation in the absence of CovS may reflect phosphorylation by acetyl phosphate or another non-cognate kinase, particularly given that THY has a lower glucose concentration than CDM and that glucose metabolism increases acetyl phosphate levels (62). Importantly, neither deletion nor overexpression of *cflP* altered CovR phosphorylation in the Δ*covS* background (Fig. 4B and C). Thus, CovS is required for CflP-dependent activation of CovR phosphorylation, consistent with CflP promoting the kinase activity of CovS.

We next examined whether Abx1 is required for CflP-dependent CovR phosphorylation. In CDM-grown cells, Δ*abx1* exhibited CovR∼P levels comparable to WT, consistent with its WT-like β-H/C phenotype under these conditions (Fig. 4B). CovR phosphorylation was undetectable in Δ*abx1*Δ*cflP*, whereas CflP overexpression restored high levels of CovR∼P in this background (Fig. 4B). Thus, CflP promotes CovR phosphorylation independently of Abx1. In contrast, THY-grown Δ*abx1* exhibited increased CovR phosphorylation, reflected by a reduction in the unphosphorylated CovR species, whereas plasmid-mediated expression of *abx1* restored the WT phosphorylation profile (Fig. 4C). These findings support the previously described role of Abx1 as a negative regulator of CovS signaling and suggest that its activity is condition-dependent. Notably, deletion of *cflP* in the Δ*abx1* background reduced CovR phosphorylation in THY-grown cells (Fig. 4C), further demonstrating that CflP promotes CovR phosphorylation independently of Abx1. Together, these results support a model in which CflP acts through CovS to favor the kinase-active state, whereas Abx1 can oppose CovS activation under conditions in which it is sufficiently abundant or active.

### Mg²⁺ regulates CflP abundance at the post-transcriptional level

To determine whether nutrient availability affects the abundance of CflP and Abx1, we generated A909 strains expressing 3×FLAG-tagged CflP or Abx1 from their native chromosomal loci (*cflP*-3×FLAG and *abx1*-3×FLAG). Both fusion strains produced WT levels of pigmentation on Granada agar and hemolysis on blood agar, indicating that the FLAG tags did not detectably impair protein function (Fig. S6A). We assessed CflP, Abx1, and CovS abundance during mid-log-phase growth in CDM and nutrient-rich media (THY, TSB, and BHI) by immunoblotting with anti-FLAG or anti-CovS antibodies. Abx1 was below the detection limit under all conditions tested and could be detected only after enrichment by anti-FLAG pull-down from 1 L of THY-grown *abx1*-3×FLAG cells (Fig. S6B), indicating that Abx1 is a low-abundance protein under these conditions.

CflP and CovS were both less abundant in CDM-HighMg than in nutrient-rich media (Fig. 5A, Fig S7). Notably, CflP was present at substantially lower levels relative to CovS in CDM-HighMg, whereas its abundance increased in BHI (Fig. 5A). Comparison of cells grown in CDM-HighMg and CDM-LowMg revealed that CovS abundance was unaffected by Mg²⁺ availability, whereas CflP abundance decreased under low-Mg²⁺ conditions (Fig. 5B). Thus, Mg²⁺ specifically influences CflP protein abundance without altering CovS levels.

**Figure 5.**
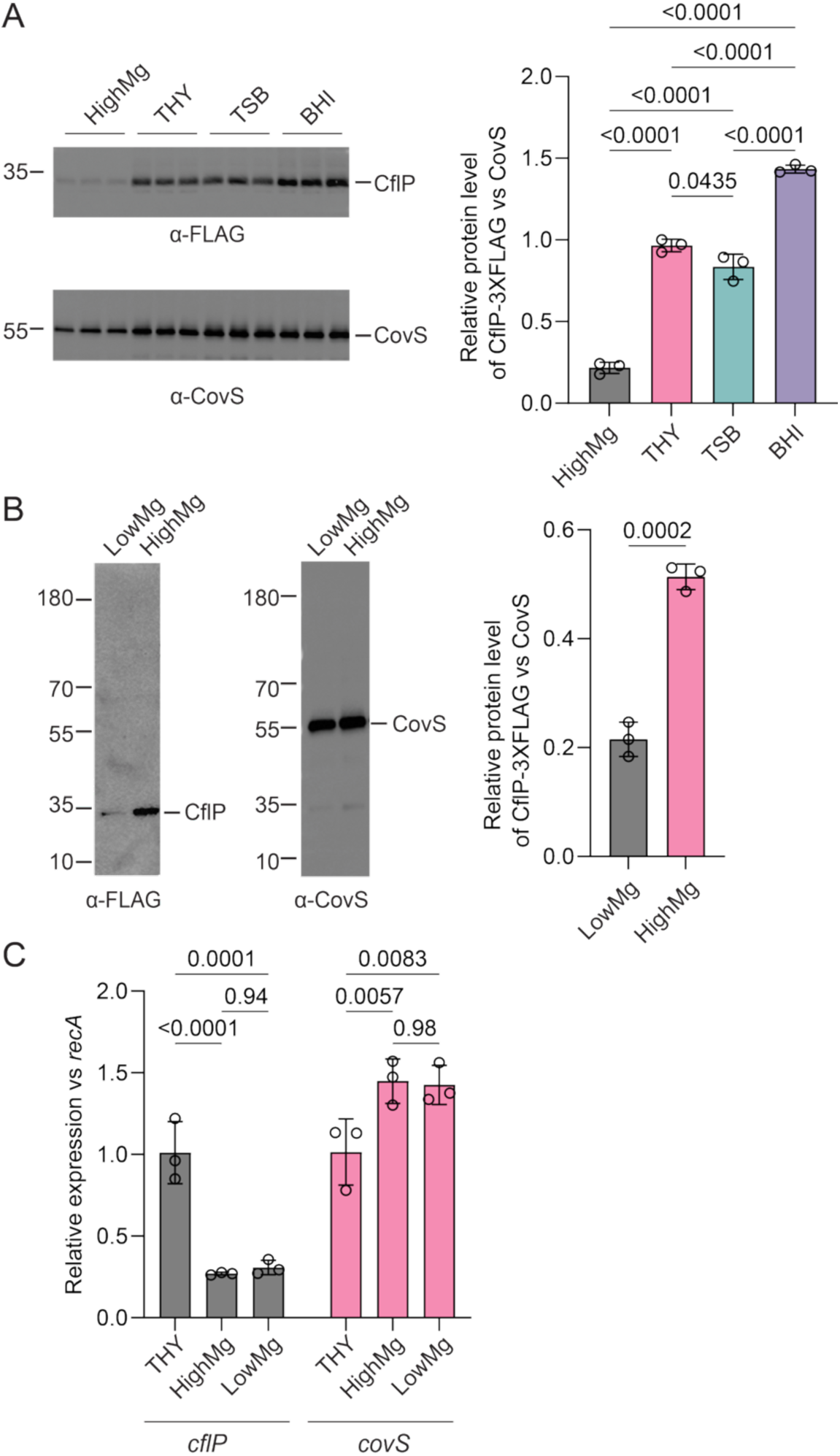
Nutrient availability and Mg²⁺ regulate CflP abundance post-transcriptionally. (**A**) Immunoblot analysis of CflP-3×FLAG and CovS in membrane fractions of mid-log-phase A909 *cflP*-3*×*FLAG cells grown in CDM-HighMg, THY, TSB, or BHI at 37 °C. Membrane proteins were isolated as described in Materials and Methods and separated on a 15% Bis-Tris gel. Samples were incubated in the presence of 100 mM DTT at 70 °C for 10 min before SDS-PAGE. Representative immunoblots from three independent biological experiments are shown; uncropped blots are provided in Fig. S7B. The specificity of the anti-FLAG and anti-CovS antibodies was confirmed using WT and Δ*covS* strains (Fig. S14). The right panel shows CflP-3×FLAG abundance normalized to CovS. Band intensities were quantified using ImageJ. Columns and error bars represent the mean ± SD (*n* = 3 biological replicates). *P* values were determined by one-way ANOVA with Tukey’s multiple-comparison test. (**B**) Immunoblot analysis of CflP-3×FLAG and CovS in membrane fractions of mid-log-phase A909 *cflP*-3*×*FLAG cells grown in CDM-HighMg or CDM-LowMg at 37 °C. Membrane proteins were separated on a 4–12% SurePAGE Bis-Tris gel (GenScript) in MOPS buffer. A representative immunoblot from three independent biological experiments is shown. The right panel shows CflP-3×FLAG abundance normalized to CovS. Band intensities were quantified using ImageJ. Columns and error bars represent the mean ± SD (*n* = 3 biological replicates). *P* values were determined by an unpaired *t* test. (**C**) qRT-PCR analysis of *cflP* and *covS* transcription in A909 cells grown to mid-log phase in THY, CDM-HighMg, or CDM-LowMg at 37 °C. Transcript levels were normalized to *recA*. Columns and error bars represent the mean ± SD (*n* = 3 biological replicates). *P* values were determined by two-way ANOVA with Tukey’s multiple-comparison test.

To determine whether Mg²⁺ regulates CflP at the transcriptional level, we measured *cflP* and *covS* transcript levels by qRT-PCR in A909 grown in CDM and THY (Fig. 5C). Growth in CDM caused only a modest increase in *covS* transcript abundance, consistent with the relatively stable CovS protein levels observed under different Mg²⁺ conditions. In contrast, *cflP* transcript levels were markedly lower in CDM than in THY but were not significantly affected by Mg²⁺ concentration within CDM. These findings indicate that although nutrient availability influences *cflP* transcription, the Mg²⁺-dependent regulation of CflP occurs primarily at the post-transcriptional level.

### CflP is a binding partner of CovS

We hypothesized that CflP activates CovS through direct physical association. To test this, we performed affinity pull-down experiments using A909 expressing chromosomally encoded CflP-3×FLAG. Because CflP is an integral membrane protein and its interaction with CovS may be transient, cells were treated with the reversible crosslinker dithiobis(succinimidyl propionate) (DSP) before membrane protein extraction and affinity purification with anti-FLAG magnetic resin. Immunoblotting of the elution fractions with anti-FLAG antibodies detected the expected ∼35-kDa CflP-3×FLAG species as well as higher-molecular-weight material above 55 kDa (Fig. 6A and B). Probing the same samples with anti-CovS antibodies detected CovS species at approximately 55, 80, and 180 kDa (Fig. 6A, B and Fig. S8B and C). The higher-molecular-weight species were consistent with crosslinked CflP–CovS complexes, whereas the ∼55-kDa species corresponded to CovS monomer. Neither CflP nor CovS was detected in corresponding elution fractions from untagged A909 WT cells, demonstrating pull-down specificity (Fig. S8B and C).

**Figure 6.**
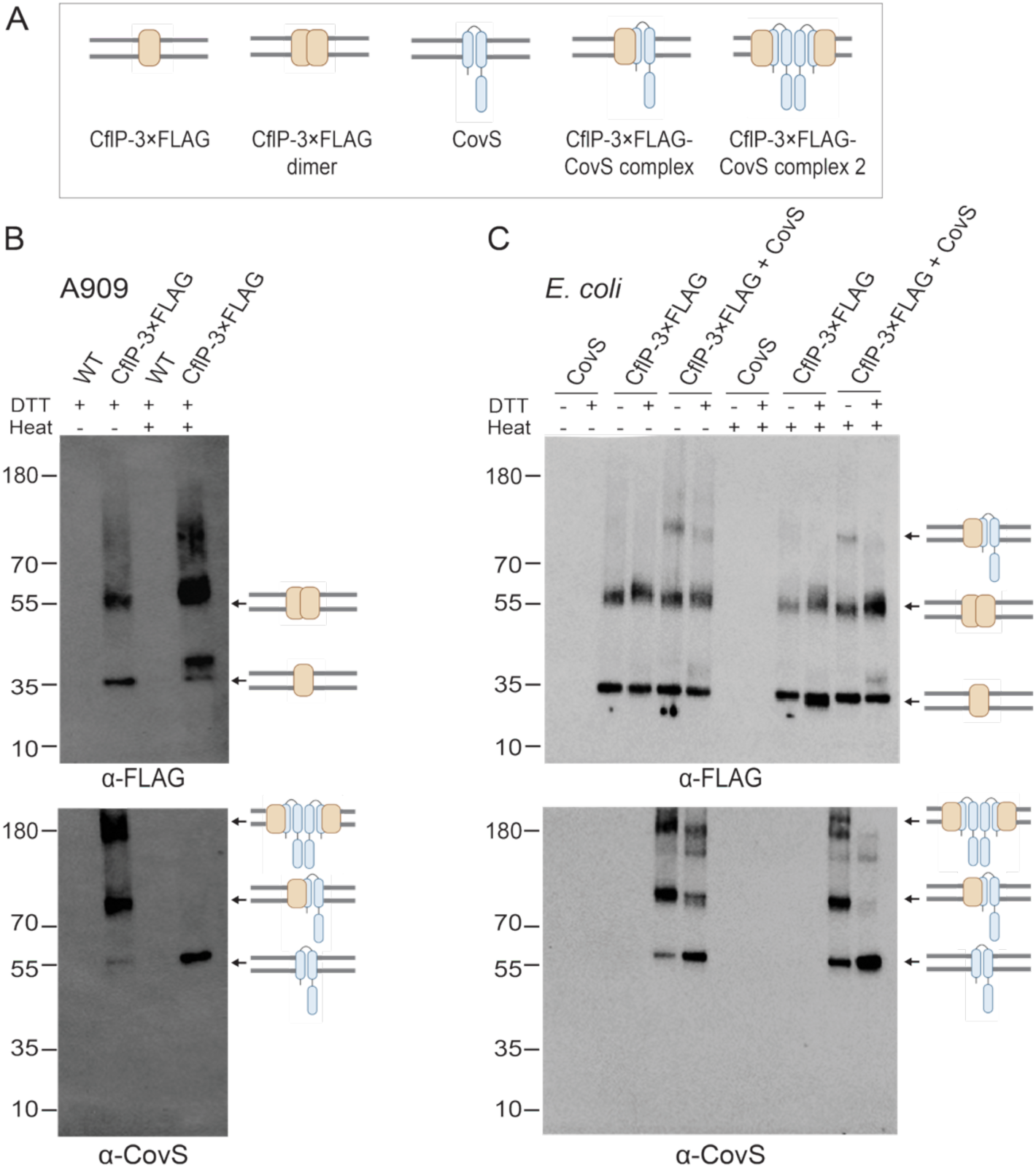
CflP is a CovS binding partner. (**A**) Proposed oligomeric states of CflP and CovS, including CflP monomers and homodimers, CovS monomers, and CflP-CovS complexes. (**B**) Immunoblot analysis of CflP-3×FLAG and CovS following FLAG affinity purification of membrane fractions from DSP-crosslinked A909 and *cflP*-3*×*FLAG cells. (**C**) Immunoblot analysis of CflP-3×FLAG and CovS following FLAG affinity purification of membrane fractions from DSP-crosslinked *E. coli* expressing CflP-3×FLAG and CovS. In **B** and **C**, samples were incubated in the presence or absence of 100 mM DTT and either maintained at room temperature or heated at 70 °C for 10 min before SDS-PAGE. Membrane proteins were separated on 4–12% SurePAGE™ Bis-Tris gels (GenScript) in MOPS buffer. CflP-3×FLAG was detected with anti-FLAG antibodies; membranes were subsequently stripped as described in Materials and Methods and probed with anti-CovS antibodies. Experiments were performed independently three times for **B** and **C**; representative images are shown. Arrows indicate bands corresponding to CflP, CovS, and putative CflP-CovS complexes consistent with the species shown in **A**.

To determine whether the higher-molecular-weight species depended on DSP crosslinking, we treated the purified complexes with DTT and heat. Because DSP contains a reducible disulfide bond, DTT disrupts DSP-mediated crosslinks. Following treatment at 70 °C for 20 min in the presence of DTT, the higher-molecular-weight CovS species disappeared, accompanied by an increase in the CovS monomer (Fig. 6A, B and Fig. S8B), supporting their identity as crosslinked CovS-containing complexes. The same treatment increased the abundance and heterogeneity of high-molecular-weight CflP species. Consistent with temperature-induced aggregation reported for some integral membrane proteins (63, 64), incubation of CflP-3×FLAG lysates at 90 °C produced similar high-molecular-weight CflP species in the absence of crosslinking (Fig. S8A).

We next asked whether CflP associates with CovS in another GBS strain. CJB111Δ*cflP* was complemented with a plasmid expressing CflP-3×FLAG, and DSP-crosslinked membrane proteins were subjected to FLAG affinity purification. CovS efficiently copurified with CflP-3×FLAG from CJB111Δ*cflP*::p*cflP*-3×FLAG but was not detected in the corresponding CJB111Δ*cflP* control (Fig. S8D). DTT treatment and heating disrupted the higher-molecular-weight CovS species and increased the abundance of CovS monomer, further confirming that the CflP–CovS complexes were DSP-dependent (Fig. S8D).

To determine whether CovS association is a general property of CPBP proteins or is specific to CflP, we examined another GBS CPBP, Abi (SAK_0054), which shares 27% amino acid identity with CflP and Abx1. Expression of Abi-3×FLAG did not alter β-H/C production in either A909 or CJB111, suggesting that Abi does not regulate β-H/C under the conditions tested (Fig. S9A). Moreover, CovS was not detected in FLAG pull-downs from DSP-crosslinked A909 or CJB111 expressing Abi-3×FLAG (Fig. S9B). Thus, CovS association is not a general property of GBS CPBPs but is specific to CflP.

Finally, we tested whether CflP–CovS complex formation requires additional GBS-specific factors. We expressed CovS, CflP-3×FLAG, or both proteins in *Escherichia coli*. Following DSP crosslinking and FLAG affinity purification, CovS was recovered specifically from cells co-expressing CovS and CflP-3×FLAG (Fig. 6A, C). These findings demonstrate that CflP and CovS are sufficient to form a stable complex in a heterologous host and strongly support a direct physical interaction between the two proteins.

### CflP and Abx1 exhibit distinct patterns of potentially disruptive sequence variation

The CflP ortholog Gbs1037 from GBS strain NEM316 was previously reported to have no effect on β-H/C production or hemolysis when overexpressed (24). CflP^NEM316^ differs from CflP^A909^ by a G106E substitution within the third predicted transmembrane helix (Fig. 7A). In the CflP model, G106 lies in a sterically constrained transmembrane environment, suggesting that substitution with a charged glutamate could impair protein folding or stability.

**Figure 7.**
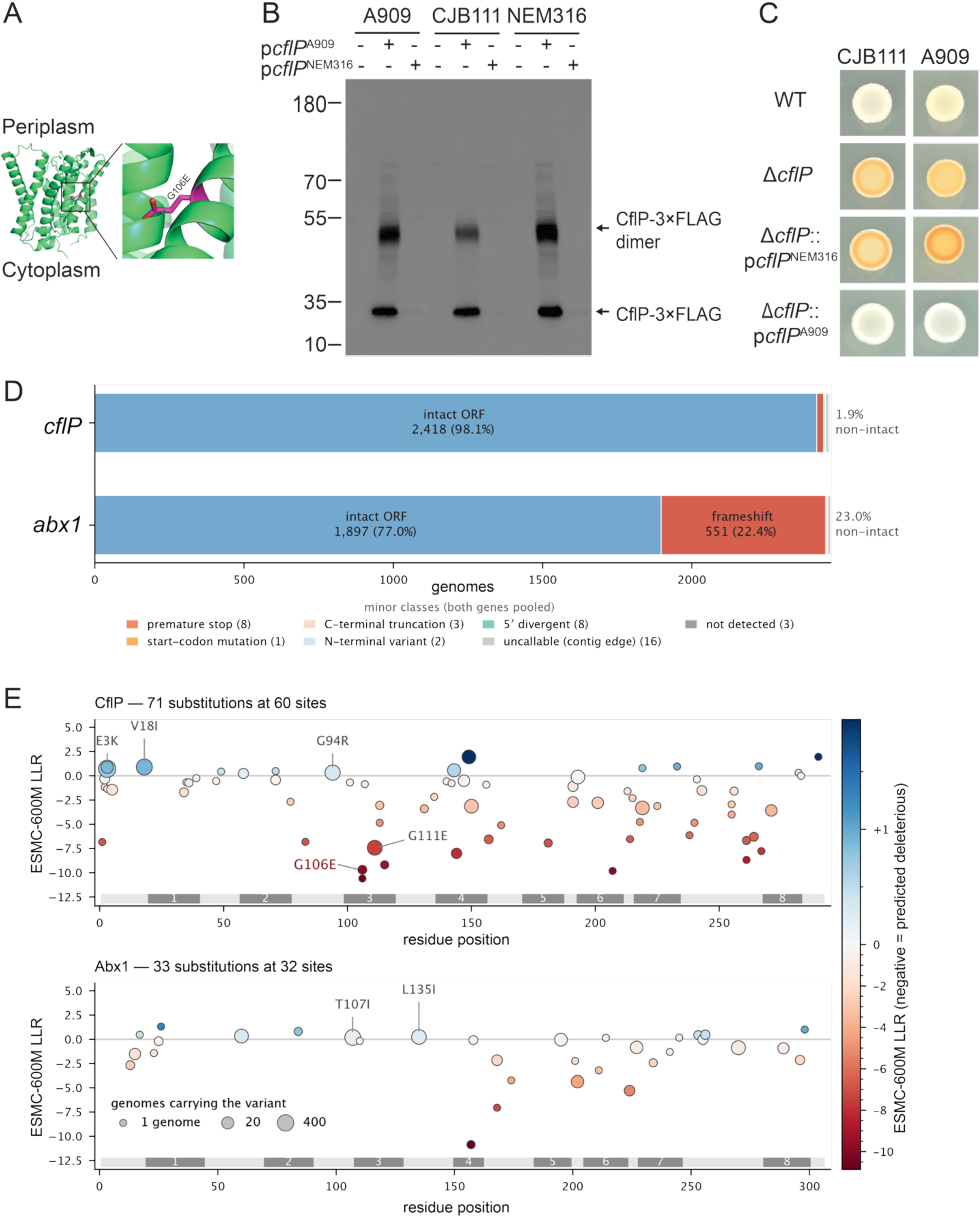
The G106E substitution impairs CflP accumulation in NEM316. (**A**) Mapping CflP^NEM316^ G106E substitution within the third transmembrane helix of CflP model. The periplasmic and cytoplasmic sides are indicated. (**B**) Immunoblot analysis of membrane-associated CflP-3×FLAG from A909, CJB111, and NEM316 strains expressing plasmid-encoded CflP from A909 (p*cflP*^A909^) or NEM316 (p*cflP*^NEM316^). Membrane proteins were isolated as described in Materials and Methods and separated on 4–12% SurePAGE™ Bis-Tris gels (GenScript) in MOPS buffer. Experiments were performed independently three times; a representative immunoblot is shown. (**C**) Pigmentation phenotypes of A909, CJB111, their respective *cflP* deletion mutants, and complemented mutants expressing p*cflP*^A909^ or p*cflP*^NEM316^. Strains were grown anaerobically on CDM agar at 40 °C for 24 h. Experiments were performed independently three times and yielded consistent results; a representative image is shown. (**D**) Classification of the *cflP* and *abx1* open reading frames in 2,465 *S. agalactiae* RefSeq assemblies. Bars represent the number of assemblies in each class; the legend lists the minor classes for both genes. (**E**) Amino-acid substitutions observed among protein alleles, plotted against residue position, with the predicted effect of each substitution on both the vertical axis and the color scale (negative = predicted deleterious; zero = neutral; positive = beneficial) and circle area proportional to the number of assemblies carrying it. Y axis denotes the log-likelihood ratio that the ESMC-600M model assigns to the substituted residue against the reference residue at the same position, given the rest of the protein as context. The grey ribbon shows the transmembrane helices. Substitutions carried by ≥100 assemblies are labeled, as is G106E.

To test this possibility, we expressed *cflP*^A909^-3×FLAG or *cflP*^NEM316^-3×FLAG in A909Δ*cflP*, CJB111Δ*cflP*, and NEM316. CflP^A909^-3×FLAG was readily detected in all three strain backgrounds, whereas CflP^NEM316^-3×FLAG was undetectable (Fig. 7B). Consistent with these differences in protein abundance, expression of *cflP*^A909^-3×FLAG, but not *cflP*^NEM316^-3×FLAG, suppressed β-H/C production in A909Δ*cflP* and CJB111Δ*cflP* (Fig. 7C). Thus, the G106E substitution markedly reduces CflP abundance and provides a mechanistic explanation for the previously reported lack of CflP activity in NEM316.

We next examined sequence variation in *cflP* and *abx1* across 2,465 *S. agalactiae* genome assemblies. The *cflP* open reading frame was intact in 98.1% of assemblies. The 2,418 intact sequences encoded 72 distinct protein alleles, 42 of which were singletons, with the A909 allele accounting for 63% of sequences (Fig. 7D; Table S1). Zero-shot mutational scoring using a state-of-the-art protein language model ESMC (65) identified G106E and G106R as the substitutions predicted to have the strongest effects on CflP sequence compatibility (Fig. 7E, Table S2). Thus, although *cflP* is generally maintained as an intact coding sequence, naturally occurring amino-acid substitutions can compromise CflP abundance and function, as demonstrated experimentally for G106E.

In contrast, *abx1* exhibited frequent coding-sequence disruption. The open reading frame was intact in 77% of assemblies; whereas 22.4% contained a frameshift; the remaining loci were at the contig boundary or contained other lesions (Fig. 7D). Missense diversity among intact sequences was comparatively limited, comprising 37 protein alleles, with the A909 allele accounting for 68.6% (Fig. 7E, Table S3). Strikingly, 543 of 551 frameshifts localized to two 5′-proximal poly-T tracts: a T_8_ tract at coding position 93, which was contracted or expanded by one nucleotide in 209 and 60 assemblies, respectively, and a T_7_ tract at position 173, which was contracted by one nucleotide in 274 assemblies (Fig. S10A, Table S4). The recurrent gain or loss of single nucleotides within these homopolymeric tracts is consistent with slipped-strand mispairing and identifies them as major sources of *abx1* coding-sequence variation.

*abx1* disruption was nonrandomly distributed among GBS lineages and host sources. Frameshift-containing alleles were enriched in CC1 and CC22, human-associated GBS lineages associated with both colonization and invasive disease in adults (66, 67), where *abx1* was disrupted in approximately 60% of assemblies compared with 22.6% across the complete panel. The two recurrent frameshift sites also differed in their distributions: variation in the position-93 T_8_ tract occurred predominantly among bovine and porcine isolates, whereas the position-173 T_7_ lesion occurred predominantly among human isolates. Host-associated differences were also evident within individual clonal complexes. For example, human CC17 and CC22 isolates predominantly retained intact *abx1*, whereas bovine isolates within these complexes frequently carried the position-93 lesion (Fig. S10B). CJB111, an ST1 strain used in this study, carries the CC1-associated position-173 frameshift, whereas A909 carries an intact *abx1* allele. In contrast to *abx1*, *cflP* disruption was uncommon across most clonal complexes (Fig. S10C).

Together, these analyses reveal distinct patterns of natural variation in the two CovS-associated CPBP proteins. *cflP* is predominantly maintained as an intact coding sequence, although individual amino-acid substitutions can compromise CflP abundance and activity. In contrast, *abx1* is frequently disrupted by recurrent frameshifts concentrated within two homopolymeric tracts. These differences suggest that the accessory regulatory inputs into CovS can vary among GBS strains and lineages.

### Loss of *cflP* increases cytotoxicity toward human immune cells

Because β-H/C promotes GBS survival during intracellular infection by damaging phagocytic cells (29), we examined whether CflP influences GBS interactions with human immune cells. Human neutrophils, monocytes, monocyte-derived macrophages, and monocyte-derived dendritic cells (moDCs) were infected with A909 or CJB111 WT and Δ*cflP* strains at an MOI of 10. Early intracellular survival of CJB111 was lower than that of A909 in neutrophils, monocytes, and macrophages but not in moDCs. However, deletion of *cflP* did not significantly alter intracellular survival in any of the cell types tested (Fig. S11A).

We next examined whether CflP affects GBS-induced cytotoxicity. To assess cytotoxicity associated with intracellular bacteria, immune cells were allowed to phagocytose GBS followed by antibiotic treatment to eliminate extracellular bacteria. Under these conditions, CJB111Δ*cflP* induced significantly greater LDH release from neutrophils and moDCs than the parental strain, whereas neither A909 nor A909Δ*cflP* produced detectable cytotoxicity (Fig. 8A). Thus, loss of CflP increases the cytotoxic potential of CJB111 without measurably affecting early intracellular survival.

**Figure 8.**
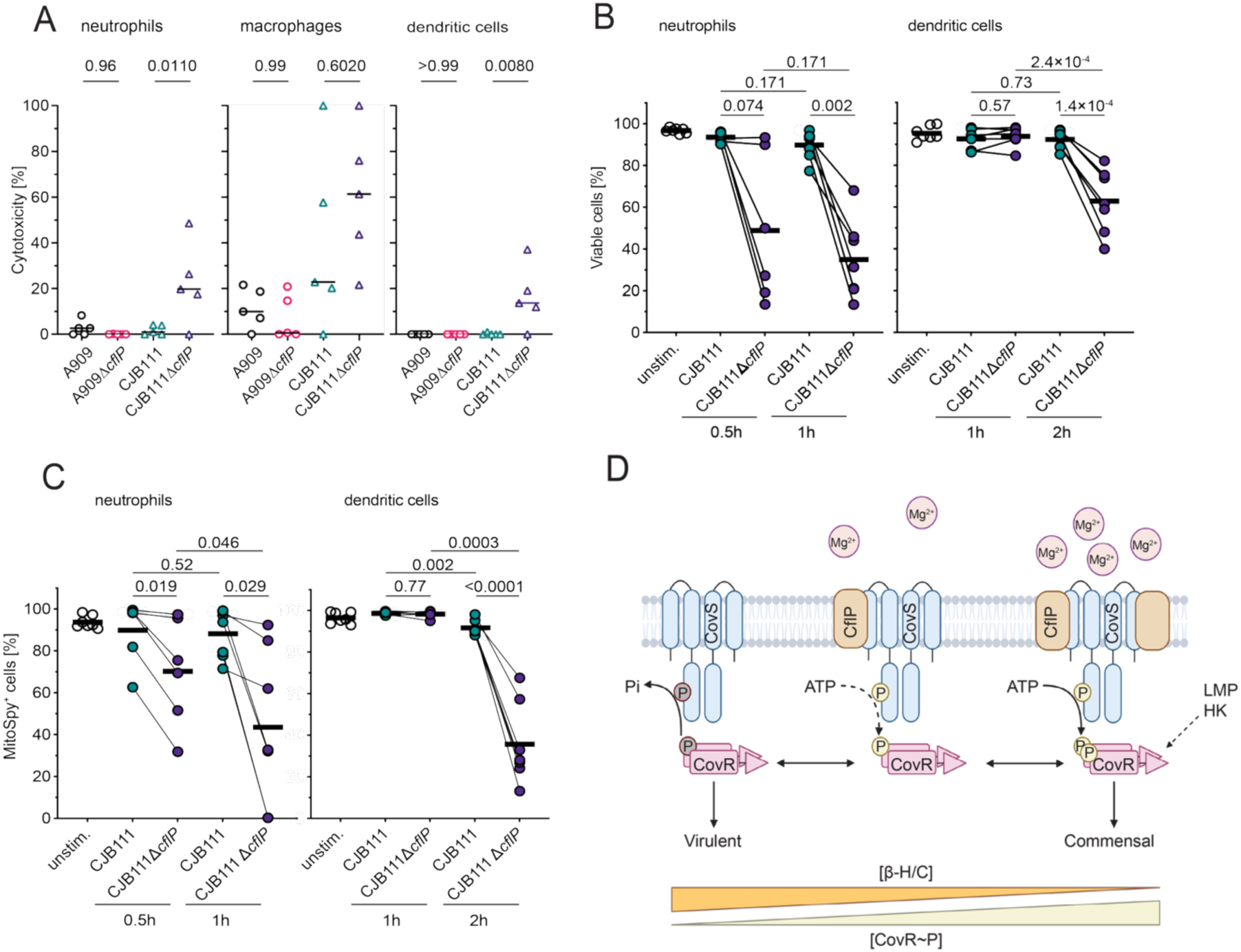
CflP restrains immune-cell damage. (**A**) Cytotoxicity of GBS toward human primary neutrophils, monocytes, and dendritic cells. Immune cells were infected with CJB111 and the isogenic *cflP* deletion mutant, CJB111Δ*cflP*, at a multiplicity of infection (MOI) of 10. Cytotoxicity was assessed by measuring lactate dehydrogenase (LDH) release into the culture supernatant. Horizontal lines represent the mean of biologically independent replicates (*n* = 5). *P*-values were calculated by one-way ANOVA with Tukey’s multiple-comparison correction. (**B**) Cell viability of human primary neutrophils and dendritic cells infected with CJB111 and CJB111Δ*cflP*. (**C**) Mitochondrial membrane potential of human primary neutrophils and dendritic cells infected with CJB111 and CJB111Δ*cflP*. In **B** and **C**, unstimulated cells served as negative controls. Immune cell damage was assessed as described in Materials and Methods. In **B–C**, each line is one biological replicate (n ≥ 6), horizontal bar is the mean; *P* values were calculated by paired *t* test on logit-transformed percentages with Holm–Šidák multiple-comparison correction. (**D**) Proposed model for CflP-dependent modulation of CovRS signaling in GBS. Under low-Mg²⁺ conditions, reduced CflP abundance is proposed to shift CovS toward its phosphatase-dominant state, decreasing CovR∼P and derepressing the β-H/C synthesis operon and other CovR-repressed genes. Conversely, high Mg²⁺ availability increases CflP abundance, promoting its association with CovS and favoring the kinase-active state, thereby promoting CovS autophosphorylation and subsequent CovR phosphorylation. Increased CovR∼P represses β-H/C and other CovR-regulated genes. A noncognate histidine kinase (HK) or low-molecular-weight phosphodonor (LMP) may also contribute to CovR phosphorylation. The relationship between CflP-dependent CovR phosphorylation and β-H/C production is indicated at the bottom of the model.

To determine whether this phenotype was directly associated with CflP-dependent regulation of β-H/C, we exposed neutrophils and moDCs to CJB111 or CJB111Δ*cflP* without antibiotic treatment, thereby allowing both cell-associated and extracellular bacteria to contribute to cytotoxicity. All infections quickly activated neutrophils (Fig. S11B, S12). In neutrophils and moDCs, CJB111Δ*cflP* caused significantly greater loss of viability than WT and induced a pronounced decrease in mitochondrial membrane potential, as measured by MitoSpy staining (Fig. 8B, C; S13). Notably, CJB111Δ*cflP* produced substantially more secreted β-H/C than CJB111, A909, or A909Δ*cflP* (Fig. 1A), correlating increased toxin production with enhanced immune-cell damage. Together, these findings indicate that CflP limits GBS cytotoxicity toward human immune cells, likely by restraining β-H/C production, and further demonstrate that CflP-dependent tuning of CovRS signaling can substantially alter the outcome of GBS–host cell interactions.

## Discussion

The CovR/CovS two-component system is a central regulator of virulence in pathogenic streptococci, including GBS and *S. pyogenes*. CovS is a bifunctional sensor histidine kinase whose kinase and phosphatase activities determine the phosphorylation state of CovR and, consequently, the expression of numerous virulence-associated genes. Here, we identify CflP, a previously uncharacterized CPBP-family membrane protein, as a CovS-associated regulator that promotes the kinase-ON state. Our findings reveal an additional layer of CovRS regulation through which GBS can fine-tune virulence in response to environmental conditions.

CflP shares the highest sequence similarity with Abx1, a CPBP protein previously shown to inhibit CovS kinase activity (24). Similar to Abx1, CflP does not require its predicted catalytic residues for regulatory activity, supporting the emerging view that CPBPs can function as noncatalytic protein scaffolds. CflP associates with CovS in two GBS strains, A909 and CJB111, and CflP and CovS form a complex when coexpressed in *E. coli*, indicating that their interaction does not require an additional GBS-specific factor. Moreover, CflP-dependent CovR phosphorylation requires CovS, strongly supporting a model in which CflP regulates CovR through CovS. We propose that CflP binding stabilizes a membrane-proximal conformation of CovS that favors the kinase-ON state and promotes CovS autophosphorylation (Fig. 8D). This model is consistent with the emerging view that many bifunctional HKs exist as ensembles of interconverting conformations and that interactions involving their transmembrane and membrane-proximal regions can shift the equilibrium between kinase- and phosphatase-competent states (68–72). Consistent with this activity, CflP overexpression caused marked CovR hyperphosphorylation and profound changes in GBS phenotypes. The ability of an accessory protein to drive CovS toward a defined signaling state resembles the activity of Abx1, which, when overexpressed, was sufficient to promote the CovS phosphatase state (24).

Our findings further indicate that CflP establishes a basal level of CovR phosphorylation that constrains expression of virulence-associated functions. Loss of CflP markedly reduced CovR phosphorylation and increased β-H/C production, abundance of virulence-associated surface proteins, and resistance to polymyxin B, a surrogate for resistance to host cationic antimicrobial peptides. Because response-regulator activity depends on both promoter architecture and phosphorylation level (14, 73), relatively small changes in CovR phosphorylation can selectively alter subsets of the CovR regulon. Consistent with this model, transcriptomic analysis showed that CflP affects a subset of genes within the established CovR regulon. Considered as transcriptional units, however, all 15 units responded in the same direction (Fig. 2E), which suggests that the effect is regulon-wide rather than gene-specific. Loss of *cflP* also increased cytotoxicity toward human immune cells in CJB111, linking CflP-dependent CovRS regulation to host-cell damage.

The ability of CflP to restrain β-H/C may be particularly important because toxin production imposes a trade-off between nutrient acquisition and host damage. β-H/C-mediated tissue injury can release nutrients that benefit GBS, but excessive toxin production causes host-cell damage and inflammation that can paradoxically impair bacterial invasion of intracellular compartments and reduce survival (13, 28, 29, 74). CflP-dependent tuning of CovS may therefore allow GBS to balance the benefits and costs of toxin production: restraining β-H/C under permissive conditions while allowing increased toxin production when nutrient limitation and host stress favor tissue damage and nutrient acquisition. In this model, CflP functions not simply as a repressor of virulence, but as a regulator that helps match the magnitude of virulence to the environmental conditions encountered by GBS.

The opposing activities of CflP and Abx1 raise the possibility that these accessory proteins compete to control the signaling state of CovS. CflP favors the kinase-ON state, whereas Abx1 promotes the phosphatase state, suggesting that their relative abundance, activity, or affinity for CovS could influence CovR phosphorylation. Such regulation could provide greater flexibility than a binary response to a single extracellular ligand and allow GBS to tune CovRS activity over a broader range of environmental conditions. CflP is itself subject to multilayered regulation. CovR directly represses *cflP* transcription (11, 14, 15, 24), and our data establish this repression under minimal media conditions (Fig. 2B). CflP therefore forms a negative-feedback circuit in which the downstream response regulator controls production of a protein that promotes CovS kinase activity. This circuit could contribute to limiting CflP-dependent activation and maintaining CovR phosphorylation within an appropriate range. Defining how this feedback circuit operates dynamically in response to environmental signals will be an important area for future investigation.

Our data further identify Mg²⁺ as a potential environmental input into this regulatory circuit. Although *cflP* transcription was substantially reduced in CDM relative to nutrient-rich media, Mg²⁺ concentration did not affect *cflP* transcript levels by qRT-PCR (Fig. 5C). In contrast, CflP protein abundance decreased under low-Mg²⁺ conditions, indicating that Mg²⁺ regulates CflP post-transcriptionally. The importance of this connection is underscored by the observation that Mg²⁺-dependent changes in β-H/C production were lost in Δ*cflP* and Δ*covS*. Mg²⁺ is a well-established environmental signal in bacterial pathogens, where depletion of this cation can indicate host nutritional immunity and trigger virulence programs (75, 76). In *S. pyogenes*, Mg²⁺ also influences CovS activity and the expression of invasive virulence factors (51–55). Our findings therefore raise the possibility that GBS uses Mg²⁺ availability to modulate CovS through CflP. Whether Mg²⁺ controls CflP translation, degradation, membrane insertion, or protein stability remains to be determined. Abx1 also appears to function conditionally: in A909 grown in CDM, *abx1* deletion had little effect on CovR phosphorylation or β-H/C production, consistent with the low abundance of Abx1 under these conditions. Together, these observations suggest that the relative abundance and activity of CflP and Abx1 may provide a mechanism for integrating environmental information into CovS signaling.

Comparative genomic analysis indicates that CflP is broadly conserved, but that naturally occurring substitutions can alter its function. The G106E substitution in NEM316 markedly reduced CflP abundance and eliminated its regulatory activity, demonstrating that sequence variation can compromise this accessory input into CovS. In contrast, *abx1* was frequently disrupted by frameshifts concentrated within two 5’-proximal homopolymeric tracts. The bidirectional length variation observed at one of these tracts is consistent with slipped-strand mispairing and raises the possibility that *abx1* may represent a phase-variable regulatory locus, although reversible switching remains to be demonstrated experimentally. Because CflP and Abx1 exert opposing effects on CovS, loss or alteration of either accessory protein could change the balance of inputs controlling CovR phosphorylation without altering the core CovRS system itself. The recurrent *abx1* frameshifts were associated with GBS lineages and host source, including differences among isolates within some clonal complexes. The frequent loss of Abx1 function, together with the strong conservation of CflP, suggests that natural variation may differentially modify the opposing accessory inputs that regulate CovS and may contribute to strain-dependent differences in CovRS signaling.

Finally, our identification of CflP as a CovS regulator expands the emerging functional repertoire of CPBP proteins. GBS genomes encode multiple CPBPs, but the functions of most remain unknown. Recent studies have shown that several CPBPs in *S. aureus* function as catalytically inactive scaffolds that interact with membrane-associated enzymes involved in cell-envelope biogenesis (77–80). Our findings extend this concept to bacterial signal transduction and suggest that CPBPs can regulate sensor HKs through direct protein-protein interactions. CPBPs may therefore represent a broader class of membrane-associated regulatory proteins that control the activity of diverse membrane enzymes and signaling proteins independently of their predicted protease activity.

In summary, we identify CflP as a noncatalytic membrane protein that associates with CovS and promotes its kinase-ON state, thereby increasing CovR phosphorylation and restraining expression of potent GBS virulence factors. Regulation of CflP abundance by environmental conditions, including Mg²⁺ availability, provides a potential mechanism for coupling CovRS signaling to environmental cues. Together with the opposing activity of Abx1, CflP expands the regulatory range of CovS and may enable GBS to tune virulence-factor production in response to changing conditions. The distinct patterns of natural variation in CflP and Abx1 further suggest that this regulatory architecture may differ among GBS strains. More broadly, our findings identify CPBPs as potential regulators of two-component signaling mechanisms and illustrate how noncatalytic accessory proteins can diversify the behavior of conserved bacterial signaling systems.

## MATERIALS AND METHODS

### Bacterial strains, antibiotics and growth conditions

Bacterial strains, plasmids, and primers used in this study are summarized in Tables S5 and S6. GBS strains (A909, CJB111, or NEM316) were grown statically at 37 °C in Todd Hewitt broth supplemented with 0.1% w/v yeast extract (THY), chemically defined medium (CDM) containing different concentrations of Mg^2+^ (CDM-HighMg and CDM-LowMg), Tryptic Soy Broth (TSB), or Brain Heart Infusion (BHI) medium containing appropriate antibiotics. For mutagenesis, if the strain carried a temperature-sensitive plasmid, growth was conducted at 30 °C. The composition of CDM-HighMg and CDM-LowMg is provided in Table S7. GBS strains were also grown on THY, CDM, BHI, blood, or Granada agar plates at 37 °C with 5% CO_2_ unless otherwise indicated. Blood and Granada agar plates were purchased from Hardy Diagnostics.

*E. coli* strains were grown in Lysogeny broth (LB) with aeration or on LB agar plates at 37 °C. The following antibiotic concentrations were used: chloramphenicol (Cm) at 15 µg mL^-1^ for *E. coli* and 5 µg mL^-1^ for GBS; kanamycin (Kan) at 60 µg mL^-1^ for *E. coli* and 60 µg mL^-1^ for GBS; and spectinomycin (Spec) at 10 µg mL^-1^ for GBS. 0.5 mM IPTG was used to induce protein expression in *E. coli*.

### Construction of GBS mutants and plasmids

Plasmids for mutant complementation (pDC123 or pJLL286 backbones (81, 82)) gene deletion (pMBsacB or pLT06 backbones (83, 84)) and *E. coli* expression (pCOLA-Duet1 or pRSF-NT backbones (85)) were constructed using standard molecular biology procedures. For gene deletions, ∼1,000-bp regions upstream and downstream of the gene of interest were PCR-amplified and cloned into pLT06 or pMBsacB. To construct complemented strains, DNA fragments containing *cflP* or *abx1* were PCR-amplified and cloned into pDC123 or pJLL286, respectively. Plasmids expressing CflP variants were generated by site-directed mutagenesis. All plasmid constructs were verified by sequencing (Eurofins Genomics).

Plasmids were introduced into GBS by electroporation, and isogenic deletion mutants were generated as described previously (83, 84). Mutant and plasmid-containing strains were confirmed by PCR and sequencing. At least three independently constructed complemented derivatives were analyzed to confirm the reproducibility of the observed phenotypes. All mutants were further validated by whole-genome sequencing to exclude secondary mutations.

### Screen of the transposon mutant library and identification of transposon insertion sites

To identify genes affecting GBS pigmentation, we screened a previously constructed saturated *Krmit* transposon library in the CJB111 background (86, 87). The library was diluted 1:10,000 in PBS, and 50 µL was plated on CDM agar and incubated anaerobically at 40 °C for 24 h in a GasPak™ EZ chamber (BD Diagnostics). Colonies exhibiting increased or decreased orange pigmentation relative to neighboring colonies were selected, inoculated into THY broth, and grown overnight at 37 °C. Cultures were collected by centrifugation and stored in BHI supplemented with 20% (v/v) glycerol for subsequent analysis.

To identify transposon insertion sites, 0.3 mL of overnight THY cultures was collected by centrifugation, resuspended in 50 µL of lysozyme (100 mg mL⁻¹), and incubated at 37 °C for 2 h. Genomic DNA was extracted using the MasterPure™ Complete DNA & RNA Purification Kit (LGC Biosearch Technologies) according to the manufacturer’s instructions. Transposon insertion sites were identified by arbitrary-primer PCR (88), followed by DNA sequencing of the resulting PCR products.

### Frozen bacterial stocks

Frozen bacterial stocks were prepared from THY-grown cultures harvested at the mid-exponential phase (OD₆₀₀ ≈ 0.5). Bacteria were collected by centrifugation, washed twice with PBS, resuspended in PBS containing 15% (v/v) glycerol, aliquoted, and stored at −80 °C until use.

### β-Hemolysin/cytolysin (β-H/C) assay

For quantitative analysis of β-H/C, GBS strains were inoculated from frozen stocks (1:50) into 5 mL of CDM and incubated at 37 °C for 16 h. Cultures were collected by centrifugation at 18,000 × *g* for 10 min, and β-H/C was extracted separately from the cell pellet and the cell-free supernatant. Cell pellets were washed once with PBS and once with 1 mL of DMSO, then resuspended in 1 mL of DMSO containing 0.2% (v/v) trifluoroacetic acid (TFA). Samples were rotated at room temperature in the dark for 1 h, then centrifuged at 18,000 × *g* for 10 min. Supernatants were collected for β-H/C quantification.

β-H/C was extracted from the cell-free supernatant by precipitating 1 mL of culture supernatant with Triton X-100 at a final concentration of 1%. The resulting pellet was dissolved in 200 µL of DMSO containing 0.2% (v/v) TFA. β-H/C was quantified by measuring absorbance at 522 nm and normalizing the signal to the final culture OD₆₀₀.

For qualitative assessment of β-H/C production on agar plates, 5 µL of bacterial suspension prepared from frozen stocks was spotted onto the indicated agar media. CDM plates were incubated anaerobically in a GasPak™ EZ chamber (BD Diagnostics) at 40 °C for 24 h. THY, BHI, and blood agar plates were incubated at 37 °C in 5% CO₂ for 24 h.

### RNA preparation and qRT-PCR

For RNA sequencing, GBS cultures were inoculated from frozen stocks and grown to mid-exponential phase (OD₆₀₀ ≈ 0.6) in CDM-HighMg or CDM-LowMg at 37 °C. A 5-mL aliquot of each culture was immediately mixed with 9 mL of RNAprotect Bacteria Reagent (Qiagen) according to the manufacturer’s recommended ratio, vortexed for 5 min, and collected by centrifugation at 18,000 × *g* for 10 min at 4 °C. Cell pellets were stored at −80 °C until RNA extraction.

Bacterial cells were resuspended in 50 µL of lysis buffer containing 10 mM Tris-HCl (pH 7.5), 1 mM EDTA, 1,250 U mL⁻¹ mutanolysin, and 1.82 mg mL⁻¹ proteinase K and 0.1-mm glass beads (Electron Microscopy Sciences) and lysed by bead beating for 20 min at room temperature using a vortex shaker. Samples were then mixed with 1 mL of QIAzol Lysis Reagent (Qiagen), preheated to 65 °C, and vortexed for 20 min. Following addition of 200 µL of chloroform, samples were centrifuged at 12,000 × *g* for 15 min at 4 °C. The upper aqueous phase was transferred to a new 2-mL tube, mixed with 1.5 volumes of 100% ethanol, and purified using RNeasy MinElute™ Cleanup spin columns (Qiagen) according to the manufacturer’s instructions. RNA concentration was measured using a NanoDrop spectrophotometer, and RNA integrity was assessed using a Qubit™ RNA IQ Assay (Invitrogen).

For qRT-PCR analysis, total RNA was extracted from A909 cultures grown to mid-exponential phase (OD₆₀₀ ≈ 0.5) in THY, CDM-HighMg, or CDM-LowMg at 37 °C, using the procedure described above. cDNA was synthesized with the iScript™ gDNA Clear cDNA Synthesis Kit (Bio-Rad). Quantitative PCR was performed using iTaq™ Universal SYBR Green Supermix (Bio-Rad). Relative expression levels of *cflP* and *covS* were calculated using the 2^-ΔΔt^ method (89), with *recA* as the reference gene, as described previously (90). Expression levels in cells grown in THY served as the reference condition for comparison with cells grown in TSB, BHI, CDM-HighMg, and CDM-LowMg.

### Gene expression analysis

#### RNA sequencing and read processing

Total RNA was prepared as described above and submitted to SeqCenter for library preparation and paired-end Illumina sequencing, with three biological replicates per strain and condition (18 libraries: A909, Δ*cflP*, and Δ*cflP*::p*cflP* in CDM-HighMg; A909, Δ*cflP*, and Δ*covR* in CDM-LowMg). The two Mg²⁺ conditions were cultured and sequenced in separate experiments. Because Mg²⁺ condition was confounded with experimental batch, direct comparisons between CDM-HighMg and CDM-LowMg were not interpreted as Mg²⁺ effects; instead, genotype effects were determined within each condition and compared across conditions. Reads were quality- and adapter-trimmed by SeqCenter, assessed with FastQC, and aligned to the GBS A909 (ATCC BAA-1138) genome using Bowtie2 v2.5.5 (91). Alignments were converted to BAM format with SAMtools v1.24 (92).

#### Read counting and differential expression

Fragments were assigned to coding sequences using featureCounts v2.1.1 (-t CDS -g ID -p --countReadPairs -s 2) and analyzed with DESeq2 v1.50.2 in R v4.5.3 (93). CDM-HighMg samples were analyzed together using a ∼strain design, allowing Δ*cflP*/WT, Δ*cflP*::p*cflP*/WT, and Δ*cflP*/Δ*cflP*::p*cflP* contrasts to share normalization and dispersion estimates. The CDM-LowMg Δ*cflP*/WT and Δ*covR*/WT comparisons were analyzed separately. No expression pre-filter was applied. Log₂ fold changes were shrunk using ashr v2.2.63, and *P* values were adjusted by the Benjamini–Hochberg method (94). Genes with adjusted *P* < 0.05 and |log₂ fold change| > 1 were considered differentially expressed.

#### Locus-tag correspondence

Reads were quantified against a Prokka annotation of the GBS ATCC BAA-1138 genome containing 2,067 protein-coding features with HFMGPFCL_ locus tags. These were mapped to the RefSeq A909 annotation (NC_007432.1) by reciprocal best BLASTP hits (BLAST+ v2.17.0) (95), requiring *E* < 1 × 10⁻⁵ and ≥50% coverage of both proteins. Unmatched proteins were mapped by TBLASTN using the same *E*-value threshold. In total, 2,060 of 2,067 quantified features were assigned to RefSeq genes; the complete correspondence is provided in Dataset S1.

#### Definition of the CovR regulon

The direct CovR regulon was defined from the published GBS BM110 ChIP-seq and transcriptomic analyses (14), which identified 21 CovR-binding loci controlling 51 genes. BM110 genes were mapped to A909 by protein homology as described above. Five loci, including *hvgA* and the *pil-1* operon, lacked A909 counterparts. The remaining loci comprised 39 A909 genes, including the 12-gene *cyl* operon, *cflP*, and 26 additional genes. *cflP* was excluded from regulon-level analyses because it was both the deleted gene and a direct member of the CovR regulon.

### Analysis of secreted proteins

GBS strains were inoculated from frozen stocks at a 1:50 dilution into 5 mL of THY and grown overnight. 1 mL of each culture was collected for analysis. Cells were removed by centrifugation at 18,000 × *g* for 10 min, and proteins in the cell-free supernatant were precipitated by adding 1 mL of a 3:1 (v/v) methanol:chloroform mixture. Samples were incubated on ice for 30 min and centrifuged at 18,000 × *g* for 20 min at 4 °C. The upper phase was removed, and 1 mL of methanol was added to the lower phase. Samples were incubated on ice for 20 min and centrifuged at 18,000 × *g* for 20 min at 4 °C. The resulting protein pellets were dried at 60 °C, resuspended in 30 µL Laemmli sample buffer, and separated by SDS-PAGE on 4–12% SurePAGE™ Bis-Tris gels (GenScript) using MOPS running buffer.

To analyze expression of Bac protein, proteins were transferred to nitrocellulose membranes by electroblotting and probed with rabbit anti-Bac antibodies (1:1,000). HRP-conjugated goat anti-rabbit IgG (Thermo Fisher Scientific, catalog no. 32460; 1:10,000) was used as the secondary antibody.

To identify differentially abundant secreted proteins, gels were stained with SYPRO Ruby according to the manufacturer’s instructions. Protein bands of interest were excised and subjected to modified in-gel trypsin digestion at the Taplin Biological Mass Spectrometry Facility (Harvard Medical School, Boston, MA). Proteins were identified by LC–MS/MS as previously described (96).

### Antimicrobial sensitivity assay

GBS cultures were grown to an OD₆₀₀ of 0.55, collected by centrifugation, resuspended in THY containing 15% (v/v) glycerol, aliquoted, and stored at −80 °C until use. GBS cultures were inoculated from these frozen stocks (1:100) in Mueller-Hinton broth containing two-fold serial dilutions of the indicated antimicrobials over a broad concentration range. For daptomycin assays, Mueller-Hinton broth was supplemented with 50 µg mL⁻¹ Ca²⁺. Aliquots (250 µL) were transferred to 96-well plates and incubated at 37 °C for 20 h. Plates were shaken before each OD₆₀₀ measurement, which was recorded at 10-min intervals using a BioTek Epoch 2 microplate spectrophotometer. Representative concentrations producing substantial to near-complete growth inhibition were selected for presentation in the figures.

### Purification of recombinant proteins from *E. coli*

Expression and purification of recombinant His-tagged CovR and CovS proteins from *E. coli*were performed as previously described (96).

### Antibody generation and purification

Rabbit polyclonal antibodies against recombinant His-tagged CovR and CovS were generated by Cocalico Biologicals. To minimize cross-reactivity, antisera were further affinity-purified using Affi-Gel 15 affinity support (Bio-Rad) according to the manufacturer’s instructions. Briefly, 20 mg of recombinant CovR or CovS was dialyzed overnight against 100 mM HEPES (pH 7.2) at 4 °C and then coupled to Affi-Gel 15 using the aqueous coupling protocol at 4 °C for 4 h. Antisera were heat-inactivated at 56 °C for 30 min to inactivate complement factors and then incubated with the antigen-coupled resin. Unbound proteins were removed by washing the resin with 5 column volumes of 1 M NaCl. Bound antibodies were eluted with 200 µL of 20 mM HCl and immediately neutralized with 100 mM Tris (pH 7.2). Antibody concentrations were determined using a NanoDrop spectrophotometer. Purified antibodies were validated by immunoblotting at a 1:10,000 dilution using lysates from the A909 WT strain and its isogenic Δ*covR* or Δ*covS* mutants (Fig. S14). Goat anti-rabbit IgG conjugated to horseradish peroxidase (HRP; Thermo Fisher Scientific, 32460) was used as the secondary antibody at a 1:10,000 dilution.

### Phos-Tag assay

Phos-Tag SDS-PAGE was performed based on previously described methods (97, 98), with modifications as described below.

#### Phos-Tag gel preparation

Phos-Tag acrylamide gels were prepared in 1-mm Novex gel cassettes using 50 µM Phos-Tag. The resolving gel was prepared by mixing 2.106 mL H₂O, 2.05 mL Acryl/Bis™ 37.5:1 (40% w/v), 1.65 mL Tris-HCl (1.5 M, pH 8.8), 60 µL Phos-Tag™ AAL-107 (5 mM aqueous solution), 66 µL MnCl₂ (10 mM), 66 µL SDS (10% w/v), 60 µL ammonium persulfate (10% w/v), and 10 µL TEMED. The stacking gel was prepared by mixing 1.26 mL H₂O, 0.22 mL Acryl/Bis™ 37.5:1 (40% w/v), 0.5 mL Tris-HCl (0.5 M, pH 6.8), 20 µL SDS (10% w/v), 20 µL ammonium persulfate (10% w/v), and 10 µL TEMED.

#### Sample preparation and Phos-Tag SDS-PAGE

GBS cultures (15 mL) grown in CDM or THY to an OD₆₀₀ of approximately 0.5 were collected by centrifugation at 5,000 × *g* for 10 min at 4 °C and washed once with 10 mL of ice-cold 20 mM Tris-HCl (pH 7.4). Cell pellets were resuspended in 0.7 mL of ice-cold lysis buffer containing 20 mM Tris-HCl (pH 7.2), 100 mM NaCl, 1% Triton X-100, 1× protease inhibitor cocktail, and 2 µg mL⁻¹ DNase and transferred to pre-chilled 2-mL screw-cap tubes containing disruptor beads. Cells were lysed using a cell disruptor for two 15-min cycles at 4 °C. Cellular debris was removed by centrifugation at 18,000 × *g* for 15 min at 4 °C.

A 100-µL aliquot of each clarified lysate was transferred to a new tube, flash-frozen in a dry ice–ethanol bath, and kept there until all samples were collected. The remaining lysate was used to determine protein concentrations using the Pierce™ BCA Protein Assay Kit (Thermo Fisher Scientific). Samples were thawed on ice, and 7.5 µg of total protein was mixed with 2× loading buffer containing 65.8 mM Tris-HCl (pH 6.8), 2.1% LDS, 26.3% (w/v) glycerol, and 0.01% (w/v) bromophenol blue. Samples were kept on ice until loading.

To confirm the phosphorylation state of CovR, two additional A909 lysates were prepared. One sample was mixed with 2× loading buffer and heated at 90 °C for 10 min, whereas the other was treated with 50 U of calf intestinal phosphatase (CIP; NEB, M0525S) at 37 °C for 30 min before adding 2× loading buffer.

Proteins were separated by Phos-Tag SDS-PAGE at 150 V for 4 h at 4 °C using running buffer containing 20 mM Tris, 191 mM glycine, and 1% SDS. Following electrophoresis, gels were incubated twice for 10 min in 10 mL of buffer Q (20 mM Tris, 191 mM glycine) containing 1 mM EDTA to remove Mn²⁺. Gels were subsequently washed twice with buffer Q without EDTA and twice with Milli-Q water to remove residual salts. Proteins were transferred to 0.45-µm nitrocellulose membranes (Bio-Rad) and immunoblotted using purified rabbit polyclonal anti-CovR antibodies (1:10,000). Ponceau S staining was used to assess protein loading and verify transfer efficiency (Fig. S5).

#### In vitro phosphorylation of CovR

Purified His-tagged CovR (20 ng) was incubated for 30 min with 20 mM acetyl phosphate in a 10-µL reaction containing 33 mM HEPES (pH 7.5), 50 mM KCl, 5 mM MgCl₂, 1 mM DTT, 0.1 mM EDTA, and 10% glycerol at room temperature. Reactions were terminated by addition of 10 µL of 2× LDS loading buffer. Reaction products were analyzed using the same Phos-Tag SDS-PAGE and immunoblotting procedure described above.

### Coimmunoprecipitation of FLAG-tagged CPBP proteins in GBS

GBS strains expressing FLAG-tagged CPBP proteins or the corresponding parental strains were used for coimmunoprecipitation experiments based on a previously published method (78) with minor modifications. Briefly, overnight cultures were diluted 1:100 into 1 L of THY and grown statically at 37 °C to an OD₆₀₀ of 0.6. Cells were collected by centrifugation, and cell pellets were resuspended in 30 mL of PBS (pH 7.4) containing 2 mM dithiobis(succinimidyl propionate) (DSP; Thermo Fisher Scientific, 22585) and incubated on ice for 2 h. The crosslinking reaction was quenched by addition of Tris-HCl (pH 7.5) to a final concentration of 20 mM, followed by incubation on ice for 15 min. Cells were collected by centrifugation, washed twice with 30 mL of PBS (pH 7.4), and stored at −80 °C until further processing.

Cell pellets were thawed on ice and resuspended in 30 mL of lysis buffer containing 1× PBS (pH 7.4), 10 µg mL⁻¹ DNase, 1× protease inhibitor cocktail, and 250 U mL⁻¹ mutanolysin. Samples were incubated at 37 °C with shaking for 30 min and then disrupted by bead beating for 10 min at 4 °C. Unlysed cells and cellular debris were removed by centrifugation at 10,000 × *g* for 10 min at 4 °C. Membrane fractions were isolated from the resulting supernatant by ultracentrifugation at 100,000 × *g* for 1 h at 4 °C.

Immunoprecipitation was performed using anti-FLAG M2 magnetic beads (Sigma-Aldrich, M8823) as previously described (78). Bound proteins were eluted by resuspending the beads in 125 µL of elution buffer containing 1× PBS (pH 7.4), 200 mM NaCl, 1% Triton X-100, and 150 µg mL⁻¹ 3×FLAG peptide (APExBIO), followed by incubation at room temperature for 15 min. Eluted proteins were collected by magnetic separation. A 10-µL aliquot of each eluate was mixed with 10 µL of 2× loading buffer containing 65.8 mM Tris-HCl (pH 6.8), 2.1% SDS, 26.3% (w/v) glycerol, and 0.01% (w/v) bromophenol blue. Where indicated, DTT was added to a final concentration of 100 mM, and/or samples were heated at 70 °C for 10 min before electrophoresis. Proteins were separated on 4–12% SurePAGE™ Bis-Tris gels using MOPS running buffer and analyzed by immunoblotting with anti-FLAG or anti-CovS antibodies.

### Coimmunoprecipitation of CflP and CovS in *E. coli*

To determine whether the interaction between CflP and CovS occurs in the absence of other GBS proteins, *E. coli* LOBSTR(DE3)::pRARE2 strains harboring plasmids encoding untagged CovS and CflP-3×FLAG (pCOLA-covS-cflP-3×FLAG), CovS alone (pCOLA-covS), or CflP-3×FLAG alone (pCOLA-3×FLAG-cflP) were used. Overnight cultures were diluted 1:100 into 600 mL of LB and grown at 37 °C with shaking to an OD₆₀₀ of approximately 0.8. Cells were collected by centrifugation, and DSP crosslinking was performed as described above. Cell pellets were resuspended in 30 mL of PBS containing 10 µg mL⁻¹ DNase and 1× protease inhibitor cocktail and lysed using a French press. Membrane fractions were isolated, and coimmunoprecipitation and immunoblot analysis were performed as described above.

### Isolation and immunoblotting of membrane proteins

Bacteria were inoculated from frozen stocks (1:1000) and grown in THY for 16 h. The inoculum (1:500) was used to grow bacteria to mid-exponential phase (OD₆₀₀ ≈ 0.6) in THY, BHI, TSB, or CDM. Cell pellets were harvested by centrifugation at 15,000 × *g* for 15 min at 4 °C and resuspended in 1 mL of PBS containing 1× protease inhibitor cocktail and 2 µg mL⁻¹ DNase. Cells were disrupted by bead beating for 30 min at 4 °C, and unlysed cells were removed by centrifugation at 5,000 × *g* for 5 min at 4 °C. Membrane fractions were collected by centrifugation of the resulting supernatants at 18,000 × *g* for 30 min at 4 °C. Membrane pellets were washed twice with PBS and resuspended in 1× PBS containing 250 mM NaCl and 1% Triton X-100. Insoluble material was removed by centrifugation at 18,000 × *g* for 15 min at 4 °C. Protein concentrations in the resulting supernatants were determined using the Pierce™ BCA Protein Assay Kit (Thermo Fisher Scientific). Equal amounts of total protein were loaded onto either 15% acrylamide gels or 4–12% SurePAGE™ Bis-Tris gels (GenScript) and separated by SDS-PAGE. Proteins were transferred to nitrocellulose membranes by electroblotting. Ponceau S staining was used to verify protein transfer and assess total protein loading. Membranes were probed with anti-CovS or anti-FLAG® antibodies (MilliporeSigma, F1807; 1:1,000). The secondary antibody was goat anti-rabbit IgG conjugated to horseradish peroxidase (HRP) (Thermo Fisher Scientific, 32460; 1:10,000).

To probe a single nitrocellulose membrane with multiple antibodies, membranes were stripped after imaging following the first round of immunoblotting using a stripping solution containing 200 mM glycine, 1% (w/v) SDS, and 1% (v/v) Tween 20, adjusted to pH 2.2. Briefly, membranes were incubated twice in 10 mL of stripping solution at 37 °C for 20 min with agitation. Membranes were then washed four times with PBS containing 0.1% (v/v) Tween 20 and reblocked with blocking solution containing 1× PBS (pH 7.4) and 7% (w/v) skim milk. Subsequent antibody incubations and immunoblotting were performed as described above.

### Structural models and analyses

Structural models of CflP (Uniprot Q8DZU5), Abx1 (Q8DYL5), MroQ (A0A0H2XEK8), and SpbR (Q2FZN5) were downloaded from the AlphaFold Protein Structure database (99, 100). The transmembrane segments and orientation relative to the plasma membrane were determined using DeepTMHMM 1.0.57 and MembraneFold 0.0.78 (101, 102). Structure superpositions were performed with TM-align (tmtools 0.3.0) (103). The structural figures were prepared using PyMOL 3.1.6.1 (104).

### Comparative genomic analysis of CflP and Abx1

#### Genome panel

All GBS assemblies available in NCBI RefSeq at the time of analysis were retrieved using the NCBI Datasets API (taxon 1311; retrieved September 1, 2026) (105), yielding 2,465 assemblies.

Identification of *cflP* and *abx1*. Because CPBP/Abi-family proteins occur in multiple copies in GBS, loci were identified using all CPBP/Abi-family proteins annotated in A909 as queries. Each assembly was searched by TBLASTN (BLAST+ v2.17.0; *E* < 1 × 10⁻⁵) (95), and loci were assigned according to the highest-scoring query. The bit-score difference from the next-best query was recorded to assess paralogue discrimination. Genomic regions encompassing each locus and 700 nt of flanking sequence were extracted and aligned to the corresponding A909 coding sequence.

#### Allele calling and classification

Loci were classified from nucleotide alignments as intact, frameshift, premature stop, start-codon mutation, C-terminal truncation, N-terminal variant, 5′ divergent, or not detected. Frameshifts were defined by insertion or deletion lengths not divisible by three and recorded relative to the A909 reference sequence. C-terminal truncations were defined as premature stops within the final 5% of the coding sequence, and N-terminal variants as a stop at codon 3 followed by an in-frame start at codon 5. Loci that could not be reliably classified because of their proximity to contig boundaries, as well as loci that were not detected, were excluded from further analysis, leaving 2,456 assemblies for *cflP* and 2,455 for *abx1*.

#### Protein alleles and substitutions

Translated intact coding sequences were collapsed into unique protein alleles and compared with the A909 reference by global pairwise alignment using Biopython v1.88 PairwiseAligner with BLOSUM62 (106) to identify amino-acid substitutions and length differences.

#### Sequence typing

Multilocus sequence types were assigned using the PubMLST *S. agalactiae* scheme (107). Alleles required full-length, 100%-identity, gap-free BLASTN matches, and sequence types were assigned by exact seven-locus profile matching. Of 2,465 assemblies, 2,295 were assigned a sequence type and 2,063 a clonal complex; assemblies lacking a clonal-complex assignment were grouped separately.

#### Variant effect prediction

Amino-acid substitution effects were estimated using ESMC-600M (esm v3.4.0; PyTorch v2.11.0) (65) by zero-shot mutational scoring. Scores were calculated as log *P*(variant) − log *P*(reference), and observed substitutions were ranked against all 19 possible substitutions at each position in the corresponding protein.

### Isolation of human primary neutrophils and monocytes and generation of dendritic cells and macrophages

Human primary neutrophils were isolated from healthy donors by density-gradient centrifugation using Polymorphprep (Axis-Shield). Neutrophils were resuspended in RPMI 1640 (Cytiva) supplemented with 10 mM L-glutamine, 25 mM HEPES (both from HyClone), and 5% (v/v) fetal calf serum (FCS).

Human monocytes were isolated from buffy coats using the EasySep™ Human CD14 Positive Selection Kit II (STEMCELL Technologies) according to the manufacturer’s instructions. Monocyte-derived dendritic cells (moDCs) were generated by culturing monocytes for 5 days in RPMI 1640 supplemented with 10% (v/v) heat-inactivated FCS (Thermo Fisher Scientific), 89 ng mL⁻¹ granulocyte-macrophage colony-stimulating factor (GM-CSF), and 22 ng mL⁻¹ interleukin-4 (IL-4) (both from ImmunoTools). The culture medium was replaced on day 3.

Monocyte-derived macrophages were generated by culturing monocytes for 7 days in RPMI 1640 (Cytiva) supplemented with 10% (v/v) heat-inactivated FCS (Thermo Fisher Scientific) and 25 ng mL⁻¹ GM-CSF. The culture medium was replaced on days 3 and 5.

### Infections of myeloid cells

All infections were performed in RPMI 1640 containing the respective supplements used for each cell type. Monocyte-derived macrophages (2 × 10⁵ cells), monocyte-derived dendritic cells (moDCs; 1 × 10⁵ cells), monocytes (1 × 10⁵ cells), or neutrophils (2 × 10⁵ cells) were infected with GBS at a multiplicity of infection (MOI) of 10 for 1 h. Following infection, extracellular bacteria were eliminated by adding RPMI 1640 containing 100 µg mL⁻¹ streptomycin and 100 IU mL⁻¹ penicillin G (HyClone). Cells were subsequently incubated for the indicated times. For neutrophils, total infection times were 2 and 4 h; for macrophages, monocytes and moDCs, total infection times were 2, 6, and 24 h. At each time point, cells were washed and lysed, and viable intracellular bacteria were quantified by determining colony-forming units (CFU) released from the lysates by plating on blood agar.

To assess host-cell viability and mitochondrial membrane potential, neutrophils and moDCs were infected as described above, except that extracellular bacteria were not killed with antibiotics. Neutrophils were analyzed after total infection times of 0.5 and 1 h, whereas moDCs were analyzed after 1 and 2 h. At each time point, cells were washed and processed for flow cytometry. For neutrophils, the first experiment was initially performed for 1 h as a test run. We observed that the effect at 1 h was already very pronounced and therefore decided to include the earlier 0.5 h time point in all subsequent experiments. Data from all experiments were analyzed (n ≥ 6). The LDH assay (Promega) was performed according to the manufacturer’s instructions.

### Flow cytometry

Cell viability was assessed using the Zombie Aqua™ Fixable Viability Kit (BioLegend), and mitochondrial membrane potential was assessed using the MitoSpy™ Orange CMTMRos Kit (BioLegend). Nonspecific Fc receptor-mediated antibody binding was blocked using Human TruStain FcX™ (BioLegend) according to the manufacturer’s instructions. Cells were incubated with the indicated monoclonal antibodies for 30 min at 4 °C in the dark. Cells were washed between staining steps and fixed using the Cyto-Fast™ Fix/Perm Buffer Set (BioLegend). The following antibodies were used (target, clone, fluorochrome; all from BioLegend): CD209, 9E9A8, APC; CD66b, G10F5, Pacific Blue; and CD15, W6D3, Alexa Fluor 700. The gating strategies are shown in Fig. S12, S13. Data were acquired using a FACSAria™ III flow cytometer and FACSDiva™ software version 8.0 (BD Biosciences, San Jose, CA, USA) and analyzed using FCS Express 7 (De Novo Software).

### Ethics

Blood samples from healthy volunteers and anonymized buffy coats obtained from the blood bank of University Medicine Greifswald were used in this study. Buffy coats were provided anonymously. Healthy volunteers were individuals familiar with the research and provided written informed consent prior to sample collection. The study was approved by the Ethics Committee of University Medicine Greifswald (Ref. Nos. BB 014/14 and BB 006/18) and conducted in accordance with the approved protocols and applicable institutional guidelines.

### Statistical analysis

Unless otherwise indicated, statistical analyses were performed using GraphPad Prism version 10. Data are presented as the mean ± standard deviation (SD) unless otherwise stated. Experiments were performed using at least three biologically independent replicates, and data from independent replicates were pooled for statistical analysis. Statistical significance was assessed using two-tailed Student’s *t*-tests, one-way or two-way ANOVA followed by Tukey’s multiple-comparisons test, or nonparametric tests followed by Dunn’s multiple-comparisons test, as appropriate. The statistical test used for each experiment is indicated in the corresponding figure legend. A *P* value ≤ 0.05 was considered statistically significant.

## Data availability

All data generated or analyzed in this study are included in the article and its supplemental material. The *S. agalactiae* A909 genome (ATCC BAA-1138), available through the ATCC Genome Portal, served as the reference for RNA-seq analysis. Whole-genome sequencing reads for the isogenic GBS mutants generated in this study have been deposited in the NCBI Sequence Read Archive (SRA) under BioProject accession number PRJNA1518224. The RNA-seq data have been deposited in the NCBI Gene Expression Omnibus (GEO) under accession number GSE344556 and are associated with BioProject accession number PRJNA1515332. The complete differential gene expression datasets for A909 WT, A909Δ*cflP*, and A909Δ*cflP*::p*cflP* grown in CDM-HighMg, and for A909 WT, A909Δ*covR* and A909Δ*cflP* grown in CDM-LowMg, are provided in Dataset S1. Accession numbers of analyzed *S. agalactiae* RefSeq assemblies with annotated CflP and Abx1 variants are provided in Table S8; the sequences are publicly available through NCBI. The mass spectrometry proteomics data have been deposited in the ProteomeXchange Consortium under dataset identifier PXD083077.

## Competing interests

The authors declare no competing interests.

## Acknowledgements

This work was supported by National Institutes of Health (NIH) grant R21AI178652 from the National Institute of Allergy and Infectious Diseases (NIAID) to N.K. The content is solely the responsibility of the authors and does not necessarily represent the official views of the NIH.

The authors thank Dr. Lakshmi Rajagopal (Seattle Children’s Hospital) for providing the Δ*covR* strain used in pilot experiments; Dr. Victor Nizet (University of California, San Diego) for providing the Δ*bac* mutant; Dr. Gunnar Lindahl (Lund University) and Dr. Alex McCarthy (Imperial College London) for providing the anti-Bac antibodies; and Dr. Kathryn Patras (Baylor College of Medicine) for providing the CJB111 and NEM316 strains.

## Author contributions

O.J.V.-C., M.M.M., N.S., K.S.M., K.V.K., and N.K. conceived and designed the experiments. O.J.V.-C., M.M.M., A.R., C.W.K, Y.L.B. and K.V.K. performed the functional and biochemical experiments. O.J.V.-C. and K.V.K. constructed plasmids and generated the GBS mutants. M.M.M. and K.V.K. performed bioinformatic analyses. O.J.V.-C., M.M.M., A.R., and K.V.K. performed statistical analyses. O.J.V.-C., M.M.M., A.R., and K.V.K. prepared the figures. O.J.V.-C., K.V.K. and N.K. wrote the manuscript with contributions from all authors. All authors reviewed and approved the final version of the manuscript.

## Notes

### Competing Interest Statement

The authors have declared no competing interest.

## References

1. Pimentel BA, Martins CA, Mendonca JC, Miranda PS, Sanches GF, Mattos-Guaraldi AL, Nagao PE. 2016. Streptococcus agalactiae infection in cancer patients: a five-year study. Eur J Clin Microbiol Infect Dis 35:927–33.

2. Russell NJ, Seale AC, O’Driscoll M, O’Sullivan C, Bianchi-Jassir F, Gonzalez-Guarin J, Lawn JE, Baker CJ, Bartlett L, Cutland C, Gravett MG, Heath PT, Le Doare K, Madhi SA, Rubens CE, Schrag S, Sobanjo-Ter Meulen A, Vekemans J, Saha SK, Ip M, Group GBSMCI. 2017. Maternal Colonization With Group B Streptococcus and Serotype Distribution Worldwide: Systematic Review and Meta-analyses. Clin Infect Dis 65:S100–S111.

3. Barshak MB. 2019. Group B Streptococcus, an A-List Pathogen in Nonpregnant Adults. JAMA Intern Med 179:488–489.

4. Coureuil M, Lecuyer H, Bourdoulous S, Nassif X. 2017. A journey into the brain: insight into how bacterial pathogens cross blood-brain barriers. Nat Rev Microbiol 15:149–159.

5. Vornhagen J, Adams Waldorf KM, Rajagopal L. 2017. Perinatal Group B Streptococcal Infections: Virulence Factors, Immunity, and Prevention Strategies. Trends Microbiol 25:919–931.

6. van Sorge NM, Doran KS. 2012. Defense at the border: the blood-brain barrier versus bacterial foreigners. Future Microbiol 7:383–94.

7. Le Doare K, Heath PT. 2013. An overview of global GBS epidemiology. Vaccine 31 Suppl 4:D7–12.

8. Seale AC, Bianchi-Jassir F, Russell NJ, Kohli-Lynch M, Tann CJ, Hall J, Madrid L, Blencowe H, Cousens S, Baker CJ, Bartlett L, Cutland C, Gravett MG, Heath PT, Ip M, Le Doare K, Madhi SA, Rubens CE, Saha SK, Schrag SJ, Sobanjo-Ter Meulen A, Vekemans J, Lawn JE. 2017. Estimates of the Burden of Group B Streptococcal Disease Worldwide for Pregnant Women, Stillbirths, and Children. Clin Infect Dis 65:S200–S219.

9. Phares CR, Lynfield R, Farley MM, Mohle-Boetani J, Harrison LH, Petit S, Craig AS, Schaffner W, Zansky SM, Gershman K, Stefonek KR, Albanese BA, Zell ER, Schuchat A, Schrag SJ, Active Bacterial Core surveillance/Emerging Infections Program N. 2008. Epidemiology of invasive group B streptococcal disease in the United States, 1999-2005. JAMA 299:2056–65.

10. Francois Watkins LK, McGee L, Schrag SJ, Beall B, Jain JH, Pondo T, Farley MM, Harrison LH, Zansky SM, Baumbach J, Lynfield R, Snippes Vagnone P, Miller LA, Schaffner W, Thomas AR, Watt JP, Petit S, Langley GE. 2019. Epidemiology of Invasive Group B Streptococcal Infections Among Nonpregnant Adults in the United States, 2008-2016. JAMA Intern Med 179:479–488.

11. Jiang SM, Ishmael N, Dunning Hotopp J, Puliti M, Tissi L, Kumar N, Cieslewicz MJ, Tettelin H, Wessels MR. 2008. Variation in the group B Streptococcus CsrRS regulon and effects on pathogenicity. J Bacteriol 190:1956–65.

12. Lamy MC, Zouine M, Fert J, Vergassola M, Couve E, Pellegrini E, Glaser P, Kunst F, Msadek T, Trieu-Cuot P, Poyart C. 2004. CovS/CovR of group B streptococcus: a two-component global regulatory system involved in virulence. Mol Microbiol 54:1250–68.

13. Patras KA, Wang NY, Fletcher EM, Cavaco CK, Jimenez A, Garg M, Fierer J, Sheen TR, Rajagopal L, Doran KS. 2013. Group B Streptococcus CovR regulation modulates host immune signalling pathways to promote vaginal colonization. Cell Microbiol 15:1154–67.

14. Mazzuoli MV, Daunesse M, Varet H, Rosinski-Chupin I, Legendre R, Sismeiro O, Gominet M, Kaminski PA, Glaser P, Chica C, Trieu-Cuot P, Firon A. 2021. The CovR regulatory network drives the evolution of Group B Streptococcus virulence. PLoS Genet 17:e1009761.

15. Lembo A, Gurney MA, Burnside K, Banerjee A, de los Reyes M, Connelly JE, Lin WJ, Jewell KA, Vo A, Renken CW, Doran KS, Rajagopal L. 2010. Regulation of CovR expression in Group B Streptococcus impacts blood-brain barrier penetration. Mol Microbiol 77:431–43.

16. Buschiazzo A, Trajtenberg F. 2019. Two-Component Sensing and Regulation: How Do Histidine Kinases Talk with Response Regulators at the Molecular Level? Annu Rev Microbiol 73:507–528.

17. Zschiedrich CP, Keidel V, Szurmant H. 2016. Molecular Mechanisms of Two-Component Signal Transduction. J Mol Biol 428:3752–75.

18. Ishii E, Eguchi Y. 2021. Diversity in Sensing and Signaling of Bacterial Sensor Histidine Kinases. Biomolecules 11.

19. Rinaldi J, Fernandez I, Shin H, Sycz G, Gunawardana S, Kumarapperuma I, Paz JM, Otero LH, Cerutti ML, Zorreguieta A, Ren Z, Klinke S, Yang X, Goldbaum FA. 2021. Dimer Asymmetry and Light Activation Mechanism in Brucella Blue-Light Sensor Histidine Kinase. mBio 12.

20. Dikiy I, Edupuganti UR, Abzalimov RR, Borbat PP, Srivastava M, Freed JH, Gardner KH. 2019. Insights into histidine kinase activation mechanisms from the monomeric blue light sensor EL346. Proc Natl Acad Sci U S A 116:4963–4972.

21. Bhate MP, Molnar KS, Goulian M, DeGrado WF. 2015. Signal transduction in histidine kinases: insights from new structures. Structure 23:981–94.

22. Jiang P, Peliska JA, Ninfa AJ. 2000. Asymmetry in the autophosphorylation of the two-component regulatory system transmitter protein nitrogen regulator II of Escherichia coli. Biochemistry 39:5057–65.

23. Neiditch MB, Federle MJ, Pompeani AJ, Kelly RC, Swem DL, Jeffrey PD, Bassler BL, Hughson FM. 2006. Ligand-induced asymmetry in histidine sensor kinase complex regulates quorum sensing. Cell 126:1095–108.

24. Firon A, Tazi A, Da Cunha V, Brinster S, Sauvage E, Dramsi S, Golenbock DT, Glaser P, Poyart C, Trieu-Cuot P. 2013. The Abi-domain protein Abx1 interacts with the CovS histidine kinase to control virulence gene expression in group B Streptococcus. PLoS Pathog 9:e1003179.

25. Claverie C, Coppolino F, Mazzuoli MV, Guyonnet C, Jacquemet E, Legendre R, Sismeiro O, De Gaetano GV, Teti G, Trieu-Cuot P, Tazi A, Beninati C, Firon A. 2024. Constitutive activation of two-component systems reveals regulatory network interactions in Streptococcus agalactiae. Nat Commun 15:9175.

26. Jiang SM, Cieslewicz MJ, Kasper DL, Wessels MR. 2005. Regulation of virulence by a two-component system in group B streptococcus. J Bacteriol 187:1105–13.

27. Lin WJ, Walthers D, Connelly JE, Burnside K, Jewell KA, Kenney LJ, Rajagopal L. 2009. Threonine phosphorylation prevents promoter DNA binding of the Group B Streptococcus response regulator CovR. Mol Microbiol 71:1477–95.

28. Doran KS, Chang JC, Benoit VM, Eckmann L, Nizet V. 2002. Group B streptococcal beta-hemolysin/cytolysin promotes invasion of human lung epithelial cells and the release of interleukin-8. J Infect Dis 185:196–203.

29. Liu GY, Doran KS, Lawrence T, Turkson N, Puliti M, Tissi L, Nizet V. 2004. Sword and shield: linked group B streptococcal beta-hemolysin/cytolysin and carotenoid pigment function to subvert host phagocyte defense. Proc Natl Acad Sci U S A 101:14491–6.

30. Park SE, Jiang S, Wessels MR. 2012. CsrRS and environmental pH regulate group B streptococcus adherence to human epithelial cells and extracellular matrix. Infect Immun 80:3975–84.

31. Sullivan MJ, Leclercq SY, Ipe DS, Carey AJ, Smith JP, Voller N, Cripps AW, Ulett GC. 2017. Effect of the Streptococcus agalactiae Virulence Regulator CovR on the Pathogenesis of Urinary Tract Infection. J Infect Dis 215:475–483.

32. Gendrin C, Merillat S, Vornhagen J, Coleman M, Armistead B, Ngo L, Aggarwal A, Quach P, Berrigan J, Rajagopal L. 2018. Diminished Capsule Exacerbates Virulence, Blood-Brain Barrier Penetration, Intracellular Persistence, and Antibiotic Evasion of Hyperhemolytic Group B Streptococci. J Infect Dis 217:1128–1138.

33. Salazar ME, Podgornaia AI, Laub MT. 2016. The small membrane protein MgrB regulates PhoQ bifunctionality to control PhoP target gene expression dynamics. Mol Microbiol 102:430–445.

34. Szurmant H, Bu L, Brooks CL, 3rd, Hoch JA. 2008. An essential sensor histidine kinase controlled by transmembrane helix interactions with its auxiliary proteins. Proc Natl Acad Sci U S A 105:5891–6.

35. Buelow DR, Raivio TL. 2010. Three (and more) component regulatory systems - auxiliary regulators of bacterial histidine kinases. Mol Microbiol 75:547–66.

36. Jiang S, Steup LC, Kippnich C, Lazaridi S, Malengo G, Lemmin T, Yuan J. 2023. The inhibitory mechanism of a small protein reveals its role in antimicrobial peptide sensing. Proc Natl Acad Sci U S A 120:e2309607120.

37. Pei J, Mitchell DA, Dixon JE, Grishin NV. 2011. Expansion of type II CAAX proteases reveals evolutionary origin of gamma-secretase subunit APH-1. J Mol Biol 410:18–26.

38. Kjos M, Snipen L, Salehian Z, Nes IF, Diep DB. 2010. The abi proteins and their involvement in bacteriocin self-immunity. J Bacteriol 192:2068–76.

39. Vogel V, Fuchs M, Jachmann M, Bitzer A, Mauerer S, Munch J, Spellerberg B. 2022. The Role of SilX in Bacteriocin Production of Streptococcus anginosus. Front Microbiol 13:904318.

40. Biswas S, Biswas I. 2014. A conserved streptococcal membrane protein, LsrS, exhibits a receptor-like function for lantibiotics. J Bacteriol 196:1578–87.

41. Lux T, Nuhn M, Hakenbeck R, Reichmann P. 2007. Diversity of bacteriocins and activity spectrum in Streptococcus pneumoniae. J Bacteriol 189:7741–51.

42. Rosa-Fraile M, Rodriguez-Granger J, Cueto-Lopez M, Sampedro A, Gaye EB, Haro JM, Andreu A. 1999. Use of Granada medium to detect group B streptococcal colonization in pregnant women. J Clin Microbiol 37:2674–7.

43. Rosa-Fraile M, Dramsi S, Spellerberg B. 2014. Group B streptococcal haemolysin and pigment, a tale of twins. FEMS Microbiol Rev 38:932–46.

44. Cheng Q, Lv S, Yin N, Wang J. 2025. Microbial regulators of physiological and reproductive health in women of reproductive age: their local, proximal and distal regulatory roles. NPJ Biofilms Microbiomes 11:207.

45. Vander Donck L, Victor M, Van Beeck W, Van Rillaer T, Dillen J, Ahannach S, Wittouck S, Allonsius CN, Lebeer S. 2025. Host-independent synergism between Lactobacillus crispatus and other vaginal lactobacilli. Cell Rep 44:116171.

46. Rosa-Fraile M, Rodriguez-Granger J, Haidour-Benamin A, Cuerva JM, Sampedro A. 2006. Granadaene: proposed structure of the group B Streptococcus polyenic pigment. Appl Environ Microbiol 72:6367–70.

47. Armistead B, Quach P, Snyder JM, Santana-Ufret V, Furuta A, Brokaw A, Rajagopal L. 2021. Hemolytic Membrane Vesicles of Group B Streptococcus Promote Infection. J Infect Dis 223:1488–1496.

48. Chaffin DO, Rubens CE. 1998. Blue/white screening of recombinant plasmids in Gram-positive bacteria by interruption of alkaline phosphatase gene (*phoZ*) expression. Gene 219:91–9.

49. Pei J, Grishin NV. 2001. Type II CAAX prenyl endopeptidases belong to a novel superfamily of putative membrane-bound metalloproteases. Trends Biochem Sci 26:275–7.

50. Manolaridis I, Kulkarni K, Dodd RB, Ogasawara S, Zhang Z, Bineva G, Reilly NO, Hanrahan SJ, Thompson AJ, Cronin N, Iwata S, Barford D. 2013. Mechanism of farnesylated CAAX protein processing by the intramembrane protease Rce1. Nature 504:301–5.

51. Gryllos I, Grifantini R, Colaprico A, Jiang S, Deforce E, Hakansson A, Telford JL, Grandi G, Wessels MR. 2007. Mg(2+) signalling defines the group A streptococcal CsrRS (CovRS) regulon. Mol Microbiol 65:671–83.

52. Gryllos I, Levin JC, Wessels MR. 2003. The CsrR/CsrS two-component system of group A Streptococcus responds to environmental Mg2+. Proceedings of the National Academy of Sciences of the United States of America 100:4227–32.

53. Gryllos I, Tran-Winkler HJ, Cheng MF, Chung H, Bolcome R, 3rd, Lu W, Lehrer RI, Wessels MR. 2008. Induction of group A Streptococcus virulence by a human antimicrobial peptide. Proc Natl Acad Sci U S A 105:16755–60.

54. Horstmann N, Sahasrabhojane P, Saldana M, Ajami NJ, Flores AR, Sumby P, Liu CG, Yao H, Su X, Thompson E, Shelburne SA. 2015. Characterization of the effect of the histidine kinase CovS on response regulator phosphorylation in group A Streptococcus. Infect Immun 83:1068–77.

55. Horstmann N, Tran CN, Brumlow C, DebRoy S, Yao H, Nogueras Gonzalez G, Makthal N, Kumaraswami M, Shelburne SA. 2018. Phosphatase activity of the control of virulence sensor kinase CovS is critical for the pathogenesis of group A streptococcus. PLoS Pathog 14:e1007354.

56. Derre-Bobillot A, Cortes-Perez NG, Yamamoto Y, Kharrat P, Couve E, Da Cunha V, Decker P, Boissier MC, Escartin F, Cesselin B, Langella P, Bermudez-Humaran LG, Gaudu P. 2013. Nuclease A (Gbs0661), an extracellular nuclease of Streptococcus agalactiae, attacks the neutrophil extracellular traps and is needed for full virulence. Mol Microbiol 89:518–31.

57. Cheng Q, Stafslien D, Purushothaman SS, Cleary P. 2002. The group B streptococcal C5a peptidase is both a specific protease and an invasin. Infect Immun 70:2408–13.

58. Zhu H, Liu M, Sumby P, Lei B. 2009. The secreted esterase of group a streptococcus is important for invasive skin infection and dissemination in mice. Infect Immun 77:5225–32.

59. Pietrocola G, Schubert A, Visai L, Torti M, Fitzgerald JR, Foster TJ, Reinscheid DJ, Speziale P. 2005. FbsA, a fibrinogen-binding protein from Streptococcus agalactiae, mediates platelet aggregation. Blood 105:1052–9.

60. Schubert A, Zakikhany K, Pietrocola G, Meinke A, Speziale P, Eikmanns BJ, Reinscheid DJ. 2004. The fibrinogen receptor FbsA promotes adherence of Streptococcus agalactiae to human epithelial cells. Infect Immun 72:6197–205.

61. Horstmann N, Saldana M, Sahasrabhojane P, Yao H, Su X, Thompson E, Koller A, Shelburne SA, 3rd. 2014. Dual-site phosphorylation of the control of virulence regulator impacts group a streptococcal global gene expression and pathogenesis. PLoS Pathog 10:e1004088.

62. McCleary WR, Stock JB. 1994. Acetyl phosphate and the activation of two-component response regulators. J Biol Chem 269:31567–72.

63. Sagne C, Isambert MF, Henry JP, Gasnier B. 1996. SDS-resistant aggregation of membrane proteins: application to the purification of the vesicular monoamine transporter. Biochem J 316 ( Pt 3):825–31.

64. Okada N, Yamamoto T, Watanabe M, Yoshimura Y, Obana E, Yamazaki N, Kawazoe K, Shinohara Y, Minakuchi K. 2011. Identification of TMEM45B as a protein clearly showing thermal aggregation in SDS-PAGE gels and dissection of its amino acid sequence responsible for this aggregation. Protein Expr Purif 77:118–23.

65. Candido S, Hayes T, Derry A, Rao R, Lin Z, Verkuil R, Wu BZ, Lee JS, Bruguera ES, Keval JA, Kopylov M, Pak JE, Wu W, Thomas N, Mataraso S, Hsu A, Trotman-Grant AC, Fatras K, dos Santos Costa A, Badkundri R, Akın H, Oktay D, Deaton J, Montabana E, Sitwala H, Yu Y, Wiggert M, Carlin DA, Goering AW, Blazejewski T, Sandora M, Hla M, Jia TZ, Kloker LH, Sofroniew NJ, Uehara M, Pannu J, Bachas S, Liu DS, Sercu T, Rives A. 2026. Language Modeling Materializes a World Model of Protein Biology. bioRxiv doi:10.64898/2026.06.03.729735:2026.06.03.729735.

66. McGee L, Chochua S, Li Z, Mathis S, Rivers J, Metcalf B, Ryan A, Alden N, Farley MM, Harrison LH, Snippes Vagnone P, Lynfield R, Smelser C, Muse A, Thomas AR, Schrag S, Beall BW. 2021. Multistate, Population-Based Distributions of Candidate Vaccine Targets, Clonal Complexes, and Resistance Features of Invasive Group B Streptococci Within the United States, 2015-2017. Clin Infect Dis 72:1004–1013.

67. Richards VP, Velsko IM, Alam MT, Zadoks RN, Manning SD, Pavinski Bitar PD, Hassler HB, Crestani C, Springer GH, Probert BM, Town CD, Stanhope MJ. 2019. Population Gene Introgression and High Genome Plasticity for the Zoonotic Pathogen Streptococcus agalactiae. Mol Biol Evol 36:2572–2590.

68. Albanesi D, Martin M, Trajtenberg F, Mansilla MC, Haouz A, Alzari PM, de Mendoza D, Buschiazzo A. 2009. Structural plasticity and catalysis regulation of a thermosensor histidine kinase. Proc Natl Acad Sci U S A 106:16185–90.

69. Saita E, Abriata LA, Tsai YT, Trajtenberg F, Lemmin T, Buschiazzo A, Dal Peraro M, de Mendoza D, Albanesi D. 2015. A coiled coil switch mediates cold sensing by the thermosensory protein DesK. Mol Microbiol 98:258–71.

70. Fernandez P, Porrini L, Albanesi D, Abriata LA, Dal Peraro M, de Mendoza D, Mansilla MC. 2019. Transmembrane Prolines Mediate Signal Sensing and Decoding in Bacillus subtilis DesK Histidine Kinase. mBio 10.

71. Inda ME, Vandenbranden M, Fernandez A, de Mendoza D, Ruysschaert JM, Cybulski LE. 2014. A lipid-mediated conformational switch modulates the thermosensing activity of DesK. Proc Natl Acad Sci U S A 111:3579–84.

72. Mensa B, Polizzi NF, Molnar KS, Natale AM, Lemmin T, DeGrado WF. 2021. Allosteric mechanism of signal transduction in the two-component system histidine kinase PhoQ. Elife 10.

73. Gao R, Stock AM. 2015. Temporal hierarchy of gene expression mediated by transcription factor binding affinity and activation dynamics. mBio 6:e00686–15.

74. Doran KS, Liu GY, Nizet V. 2003. Group B streptococcal beta-hemolysin/cytolysin activates neutrophil signaling pathways in brain endothelium and contributes to development of meningitis. J Clin Invest 112:736–44.

75. Groisman EA, Duprey A, Choi J. 2021. How the PhoP/PhoQ System Controls Virulence and Mg(2+) Homeostasis: Lessons in Signal Transduction, Pathogenesis, Physiology, and Evolution. Microbiol Mol Biol Rev 85:e0017620.

76. Groisman EA, Hollands K, Kriner MA, Lee EJ, Park SY, Pontes MH. 2013. Bacterial Mg2+ homeostasis, transport, and virulence. Annu Rev Genet 47:625–46.

77. Willing S, Schneewind O, Missiakas D. 2021. Regulated cleavage of glycan strands by the murein hydrolase SagB in S. aureus involves a direct interaction with LyrA (SpdC). J Bacteriol 203.

78. Park Y, Sisley TA, Stone MC, Corrigan RM, Vickery CR, Grundling A, Walker S. 2025. SpbR controls lipoteichoic acid length by directly inhibiting signal peptidase SpsB in Staphylococcus aureus. Proc Natl Acad Sci U S A 122:e2426464122.

79. Gardiner JHt, Komazin G, Matsuo M, Cole K, Gotz F, Meredith TC. 2020. Lipoprotein N-Acylation in Staphylococcus aureus Is Catalyzed by a Two-Component Acyl Transferase System. mBio 11.

80. Schaefer K, Owens TW, Page JE, Santiago M, Kahne D, Walker S. 2021. Structure and reconstitution of a hydrolase complex that may release peptidoglycan from the membrane after polymerization. Nat Microbiol 6:34–43.

81. Mascari CA, Djoric D, Little JL, Kristich CJ. 2022. Use of an Interspecies Chimeric Receptor for Inducible Gene Expression Reveals that Metabolic Flux through the Peptidoglycan Biosynthesis Pathway is an Important Driver of Cephalosporin Resistance in Enterococcus faecalis. J Bacteriol 204:e0060221.

82. Mascari CA, Little JL, Kristich CJ. 2023. PASTA-kinase-mediated signaling drives accumulation of the peptidoglycan synthesis protein MurAA to promote cephalosporin resistance in Enterococcus faecalis. Mol Microbiol 120:811–829.

83. Hooven TA, Bonakdar M, Chamby AB, Ratner AJ. 2019. A Counterselectable Sucrose Sensitivity Marker Permits Efficient and Flexible Mutagenesis in Streptococcus agalactiae. Appl Environ Microbiol 85.

84. Thurlow LR, Thomas VC, Hancock LE. 2009. Capsular polysaccharide production in Enterococcus faecalis and contribution of CpsF to capsule serospecificity. J Bacteriol 191:6203–10.

85. Korotkov KV, Delarosa JR, Hol WGJ. 2013. A dodecameric ring-like structure of the N0 domain of the type II secretin from enterotoxigenic Escherichia coli. J Struct Biol 183:354–362.

86. Burcham LR, Akbari MS, Alhajjar N, Keogh RA, Radin JN, Kehl-Fie TE, Belew AT, El-Sayed NM, McIver KS, Doran KS. 2022. Genomic Analyses Identify Manganese Homeostasis as a Driver of Group B Streptococcal Vaginal Colonization. mBio 13:e0098522.

87. Burcham LR, Le Breton Y, Radin JN, Spencer BL, Deng L, Hiron A, Ransom MR, Mendonca JDC, Belew AT, El-Sayed NM, McIver KS, Kehl-Fie TE, Doran KS. 2020. Identification of Zinc-Dependent Mechanisms Used by Group B Streptococcus To Overcome Calprotectin-Mediated Stress. mBio 11.

88. Das S, Noe JC, Paik S, Kitten T. 2005. An improved arbitrary primed PCR method for rapid characterization of transposon insertion sites. J Microbiol Methods 63:89–94.

89. Livak KJ, Schmittgen TD. 2001. Analysis of relative gene expression data using real-time quantitative PCR and the 2(-Delta Delta C(T)) Method. Methods 25:402–8.

90. Gopalakrishna KP, Hillebrand GH, Bhavana VH, Elder JL, D’Mello A, Tettelin H, Hooven TA. 2023. Group B Streptococcus Cas9 variants provide insight into programmable gene repression and CRISPR-Cas transcriptional effects. Commun Biol 6:620.

91. Langmead B, Salzberg SL. 2012. Fast gapped-read alignment with Bowtie 2. Nat Methods 9:357–9.

92. Danecek P, Bonfield JK, Liddle J, Marshall J, Ohan V, Pollard MO, Whitwham A, Keane T, McCarthy SA, Davies RM, Li H. 2021. Twelve years of SAMtools and BCFtools. Gigascience 10.

93. Love MI, Huber W, Anders S. 2014. Moderated estimation of fold change and dispersion for RNA-seq data with DESeq2. Genome Biol 15:550.

94. Benjamini Y, Hochberg Y. 1995. Controlling the false discovery rate: a practical and powerful approach to multiple testing. Journal of the Royal Statistical Society: Series B (Methodological) 57:289–300.

95. Camacho C, Coulouris G, Avagyan V, Ma N, Papadopoulos J, Bealer K, Madden TL. 2009. BLAST+: architecture and applications. BMC Bioinformatics 10:421.

96. Rahman MM, Zamakhaeva S, Rush JS, Chaton CT, Kenner CW, Hla YM, Tsui HT, Uversky VN, Winkler ME, Korotkov KV, Korotkova N. 2025. Glycosylation of serine/threonine-rich intrinsically disordered regions of membrane-associated proteins in streptococci. Nat Commun 16:4011.

97. Barbieri CM, Stock AM. 2008. Universally applicable methods for monitoring response regulator aspartate phosphorylation both in vitro and in vivo using Phos-tag-based reagents. Anal Biochem 376:73–82.

98. Wayne KJ, Li S, Kazmierczak KM, Tsui HC, Winkler ME. 2012. Involvement of WalK (VicK) phosphatase activity in setting WalR (VicR) response regulator phosphorylation level and limiting cross-talk in Streptococcus pneumoniae D39 cells. Mol Microbiol 86:645–60.

99. Jumper J, Evans R, Pritzel A, Green T, Figurnov M, Ronneberger O, Tunyasuvunakool K, Bates R, Zidek A, Potapenko A, Bridgland A, Meyer C, Kohl SAA, Ballard AJ, Cowie A, Romera-Paredes B, Nikolov S, Jain R, Adler J, Back T, Petersen S, Reiman D, Clancy E, Zielinski M, Steinegger M, Pacholska M, Berghammer T, Bodenstein S, Silver D, Vinyals O, Senior AW, Kavukcuoglu K, Kohli P, Hassabis D. 2021. Highly accurate protein structure prediction with AlphaFold. Nature 596:583–589.

100. Varadi M, Bertoni D, Magana P, Paramval U, Pidruchna I, Radhakrishnan M, Tsenkov M, Nair S, Mirdita M, Yeo J, Kovalevskiy O, Tunyasuvunakool K, Laydon A, Zidek A, Tomlinson H, Hariharan D, Abrahamson J, Green T, Jumper J, Birney E, Steinegger M, Hassabis D, Velankar S. 2024. AlphaFold Protein Structure Database in 2024: providing structure coverage for over 214 million protein sequences. Nucleic Acids Res 52:D368–D375.

101. Hallgren J, Tsirigos KD, Pedersen MD, Almagro Armenteros JJ, Marcatili P, Nielsen H, Krogh A, Winther O. 2022. DeepTMHMM predicts alpha and beta transmembrane proteins using deep neural networks. bioRxiv doi:10.1101/2022.04.08.487609:2022.04.08.487609.

102. Gutierrez S, Tyczynski WG, Boomsma W, Teufel F, Winther O. 2022. MembraneFold: Visualising transmembrane protein structure and topology. bioRxiv doi:10.1101/2022.12.06.518085:2022.12.06.518085.

103. Zhang Y, Skolnick J. 2005. TM-align: a protein structure alignment algorithm based on the TM-score. Nucleic Acids Res 33:2302–9.

104. Schrödinger LLC. 2026. The PyMOL Molecular Graphics System, Version 3.1.

105. O’Leary NA, Cox E, Holmes JB, Anderson WR, Falk R, Hem V, Tsuchiya MTN, Schuler GD, Zhang X, Torcivia J, Ketter A, Breen L, Cothran J, Bajwa H, Tinne J, Meric PA, Hlavina W, Schneider VA. 2024. Exploring and retrieving sequence and metadata for species across the tree of life with NCBI Datasets. Sci Data 11:732.

106. Cock PJ, Antao T, Chang JT, Chapman BA, Cox CJ, Dalke A, Friedberg I, Hamelryck T, Kauff F, Wilczynski B, de Hoon MJ. 2009. Biopython: freely available Python tools for computational molecular biology and bioinformatics. Bioinformatics 25:1422–3.

107. Jolley KA, Bray JE, Maiden MCJ. 2018. Open-access bacterial population genomics: BIGSdb software, the PubMLST.org website and their applications. Wellcome Open Res 3:124.

