## Supplementary material for "CflP couples magnesium availability to CovS signaling in Group B Streptococcus": Figures S1 to S14; Tables S1 to S7

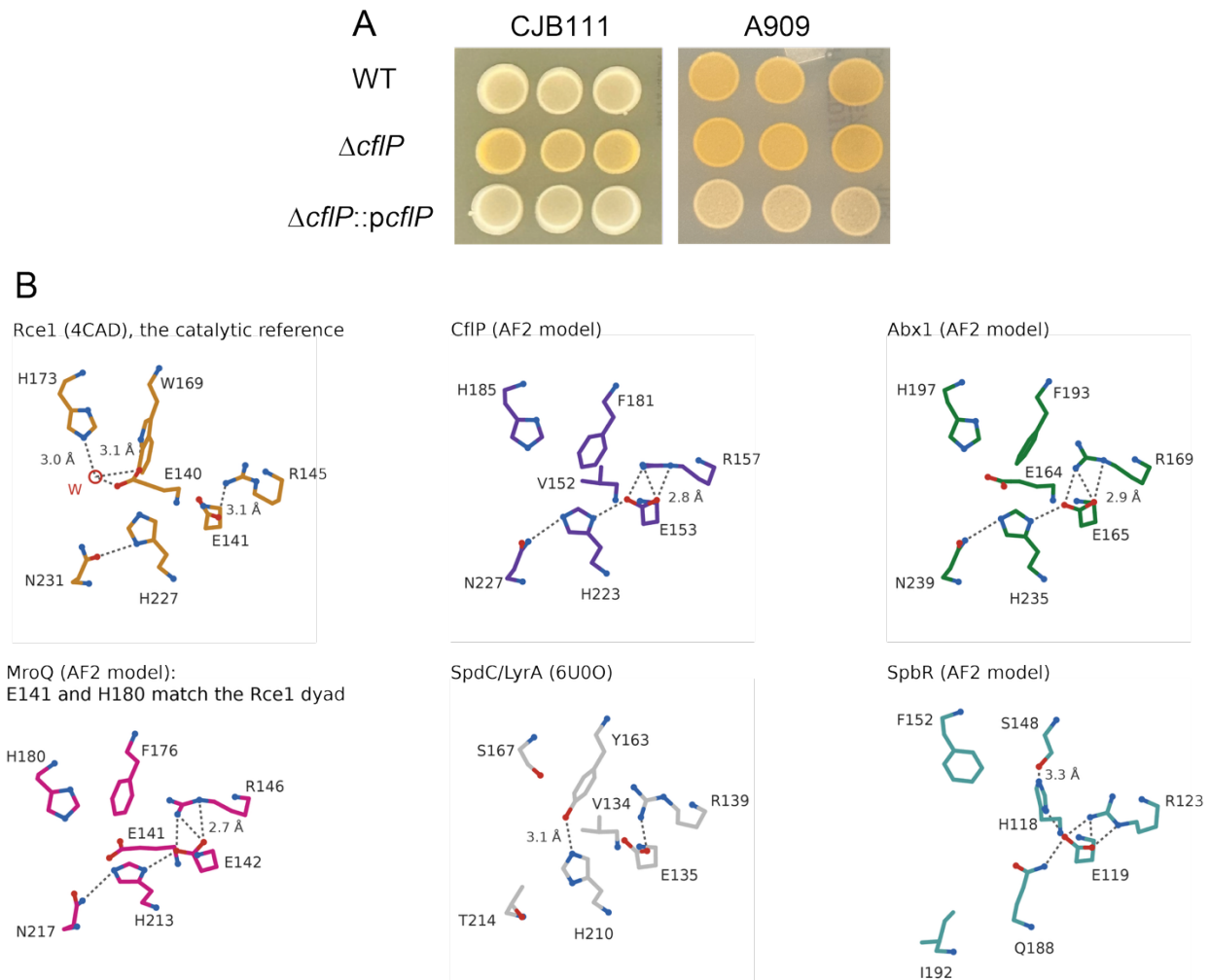

**Figure S1. CfIP negatively regulates β-H/C production and the CPBP family conserved sequence motifs.**

(A) Pigmentation phenotypes of A909 and CJB111, their respective isogenic *cfIP* deletion mutants, and complemented mutants expressing the WT *cfIP* gene. GBS strains were grown on Granada agar at 37 °C for 24 h. Experiments were performed independently twice and yielded consistent results; representative images are shown. (B) The three sequence motifs that define a proteolytically competent CPBP enzyme, EExxxR (motif 1), FxxxH (motif 2) and HxxxB (motif 3, where B is asparagine or aspartate), contribute seven side chains to a single site. The residues corresponding to the Rce1 motifs were identified using TM-align (1). Whole-chain TM-align onto CfIP, normalized by CfIP, gives 0.92 for Abx1, 0.65 for SpdC/LyrA, 0.60 for Rce1, 0.51 for MroQ and 0.46 for SpbR. Seven-point fits onto the CfIP model give Cα root-mean-square deviations of 0.26 Å for Abx1, 0.87 Å for SpdC/LyrA, 0.99 Å for Rce1, 1.27 Å for SpbR and 1.33 Å for MroQ. Each sub-panel shows those seven residues, with the backbone reduced to N and Cα and the carbonyl omitted. Nitrogen is blue and oxygen red; dashed lines mark side-chain nitrogen or oxygen pairs 2.4 to 3.6 Å apart. CfIP, Abx1, MroQ and SpbR are AlphaFold2 models, with pLDDT between 93 and 99 at every residue drawn.

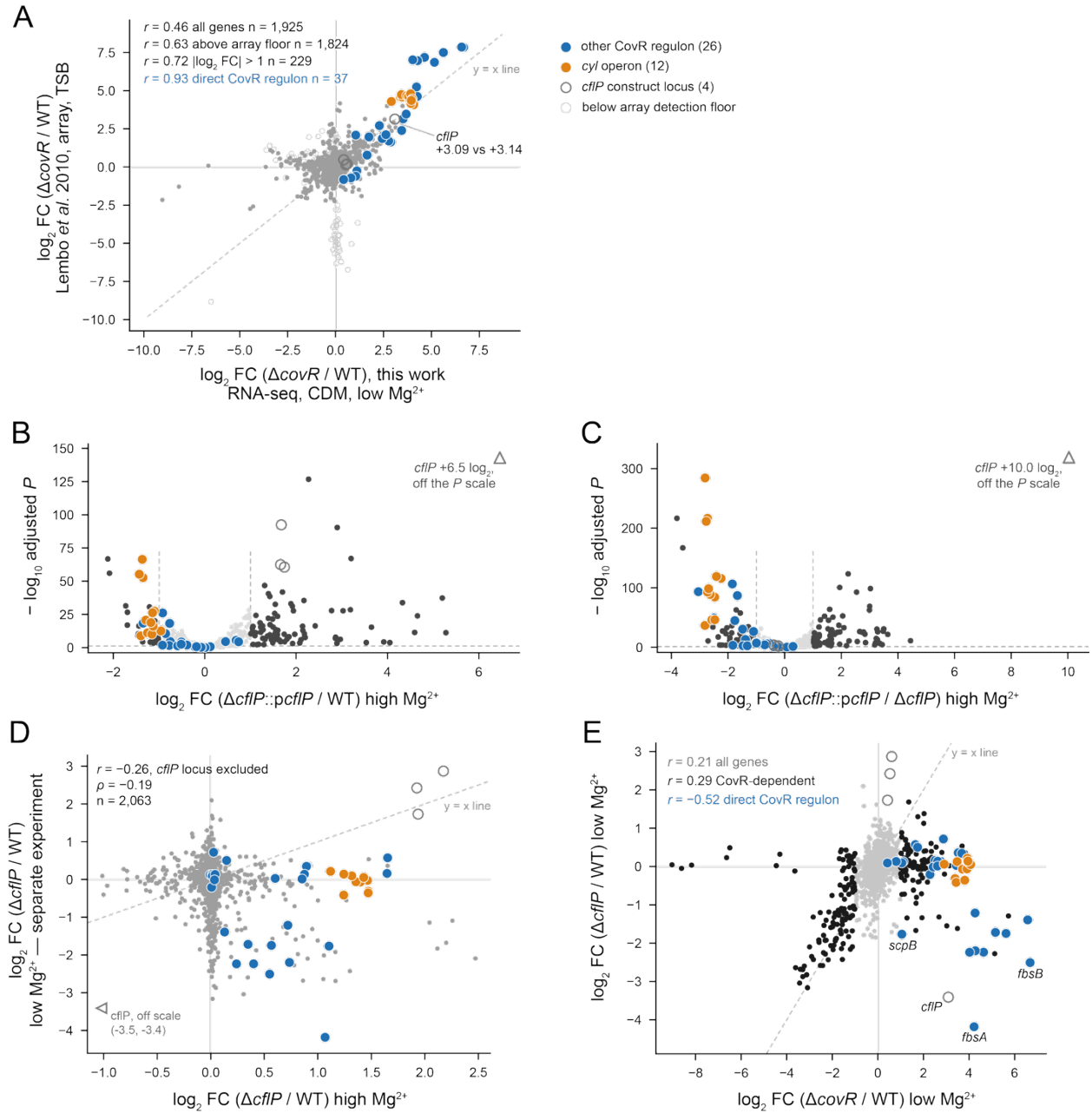

**Figure S2. CflP and CovR transcriptome comparisons.**

(A) Cross-platform validation of the $\Delta covR$ dataset. Per-gene $\log_2$ fold change in $\Delta covR$ relative to A909 WT in this work (RNA-seq, CDM-lowMg) against the published A909 $\Delta covR$ microarray of Lembo *et al.* (2), for the 1,925 genes present on both platforms; the dashed line is $y = x$. Genes were matched by A909 locus tag. Genes of the *cyl* operon are orange and other CovR regulon genes blue; the four genes of the *cflP* locus are open circles. Remaining genes are grey, drawn in a lighter shade for low-intensity probes. Pearson correlation is 0.46 across all genes, 0.63 above that intensity floor, 0.72 among the 229 genes changing more than two-fold in this work, and 0.93 for the 37 CovR regulon genes carrying a probe (SAK\_RS09660 has no probe). The agreement is near zero among genes with no signal ($r = -0.01$ for $|\log_2 FC| < 0.25$, $n = 1,222$), and rises monotonically to $r = +0.79$ for genes above four-fold. **(B)** Volcano plot of $\Delta cflP::pcflP$ relative to WT and **(C)** relative to $\Delta cflP$, both in CDM-highMg. Dashed lines mark the applied thresholds (adjusted $P < 0.05$, $|\log_2 \text{fold change}| > 1$), which 127 and 169 genes satisfy, respectively. In both panels, *cflP* itself lies off the vertical scale; its adjusted $P$ underflows the double-precision floor and is drawn as an open triangle; every other gene is on scale. Together, the panels show that the plasmid complementation reverses the direction of the deletion effect on the CovR regulon and carries it past the WT level. **(D)** Per-gene $\log_2$ fold change for $\Delta cflP$ relative to WT in CDM-highMg against the same contrast in CDM-lowMg, for all 2,063 genes with the *cflP* construct locus excluded; $r = -0.26$, Spearman $\rho = -0.19$. This comparison crosses experiments and the anti-correlation cannot be separated from batch; it is shown to establish that the CDM-highMg gene set is not recovered in the CDM-lowMg conditions, not to quantify a magnesium effect. **(E)** Loss of CflP does not reproduce loss of CovR. Per-gene $\log_2$ fold change for $\Delta covR$ against $\Delta cflP$, both relative to WT and in CDM-lowMg. All 38 CovR regulon genes rise on loss of CovR, yet 18 of the 38 move in the opposite direction on loss of CflP at this concentration ($r = 0.21$ genome-wide).

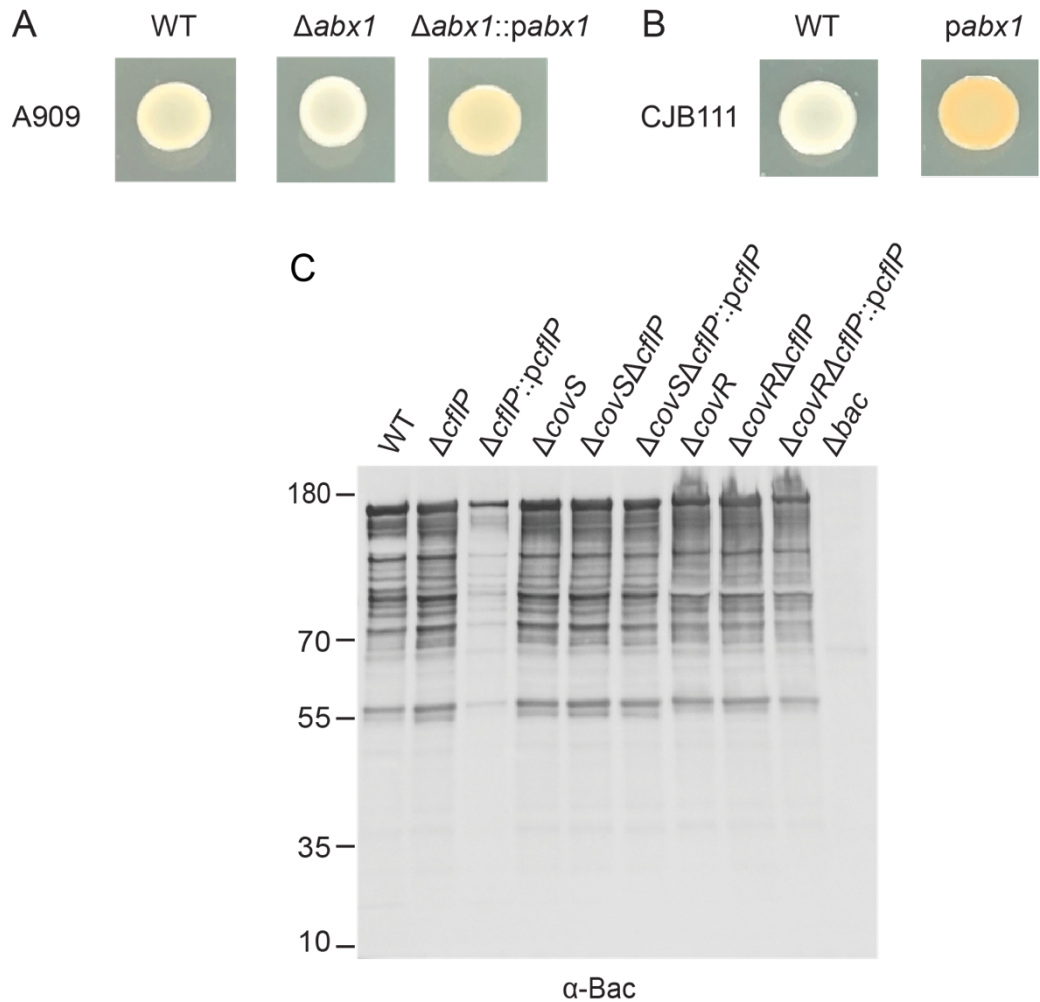

**Figure S3. CflP and Abx1 differentially regulate CovRS-dependent phenotypes in GBS.**

**(A)** Pigmentation phenotypes of A909, an isogenic mutant lacking *abx1*, and the complemented mutant expressing a WT copy of *abx1*. **(B)** Pigmentation phenotypes of CJB111 and CJB111 carrying a plasmid expressing a WT copy of *abx1*. In **A** and **B**, GBS strains were grown on CDM-LowMg agar plates under anaerobic conditions at 40 °C for 24 h. **(C)** Immunoblot analysis of Bac (β-protein) in cell-free culture supernatants from GBS strains (A909 background) grown in THY broth at 37 °C for 24 h. Bac was detected using anti-Bac antibodies. An isogenic *bac* deletion mutant ($\Delta bac$) in the A909 background served as a negative control. In **A–C**, experiments were performed independently twice and yielded consistent results; representative images are shown.

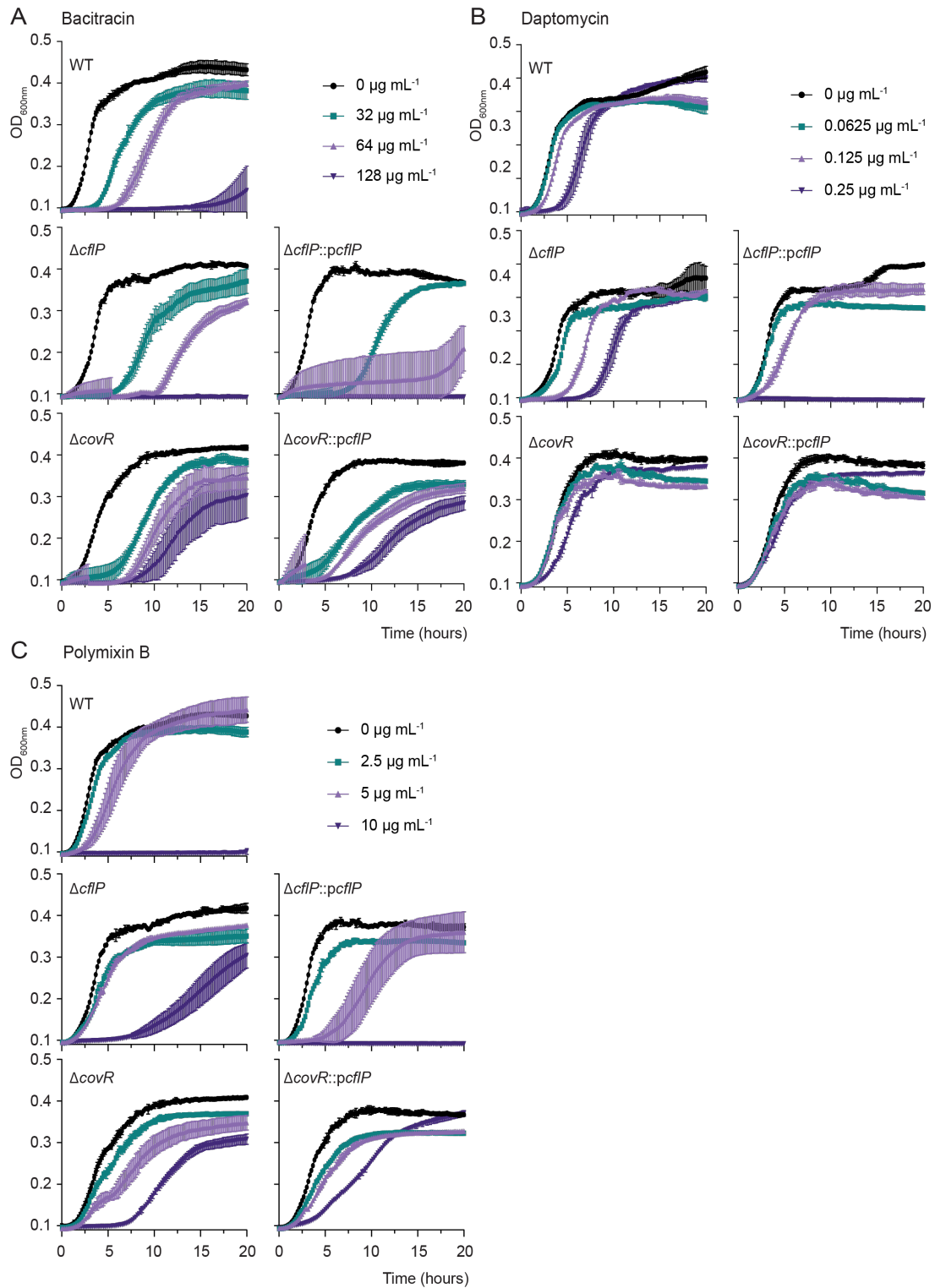

**Figure S4. CflP influences GBS A909 susceptibility to membrane-active and cell wall-targeting antimicrobials in a CovR-dependent manner.**

Growth curves of A909 WT, isogenic mutants lacking *cflP* or *covR*, and the corresponding complemented mutants in the presence of increasing concentrations of bacitracin (A), daptomycin

(**B**), or polymyxin B (**C**). Strains were grown in Mueller-Hinton broth as described in Materials and Methods. Curves represent the mean ± S.D. of three biologically independent replicates ($n = 3$).

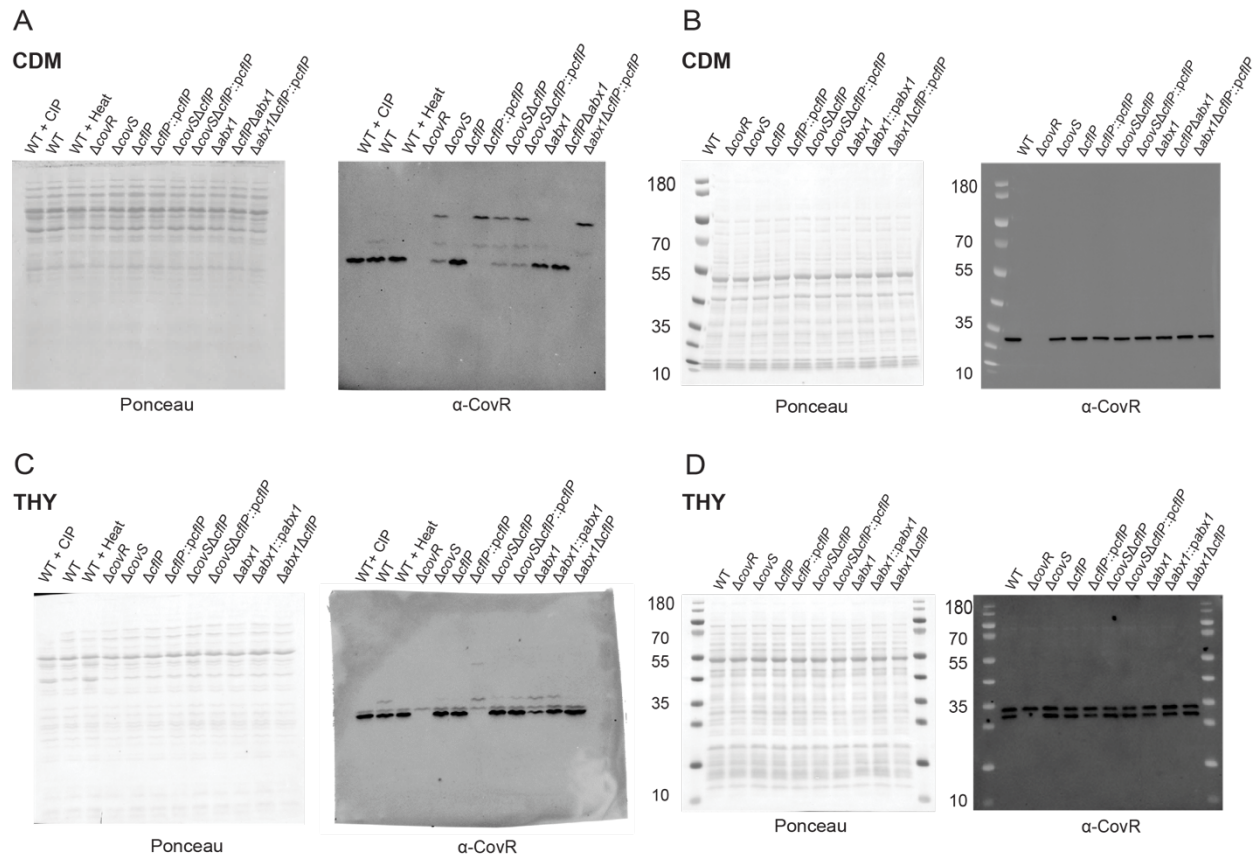

**Figure S5. CflP regulates CovR phosphorylation in GBS A909 grown in CDM and THY.**

Representative Ponceau S-stained membranes and immunoblots of Phos-tag (A and C) and conventional SDS-PAGE (B and D) gels showing CovR in cell extracts from GBS strains grown to exponential phase in CDM-LowMg (A and B) or THY (C and D) at 37 °C. Phos-tag immunoblots were used to assess CovR phosphorylation, whereas conventional SDS-PAGE immunoblots were used to assess total CovR protein levels. Growth, sample preparation, Phos-tag SDS-PAGE, and immunoblotting with anti-CovR antibodies were performed as described in Materials and Methods. Ponceau S staining was used to assess protein loading and transfer. Samples shown in A–D correspond to those analyzed in Fig. 4B and C. Proteins were separated on 4–12% SurePAGE™ Bis-Tris gels (GenScript) in MOPS buffer. Experiments were performed independently twice in A, B, and three times in D and C. A representative image from one experiment is shown.

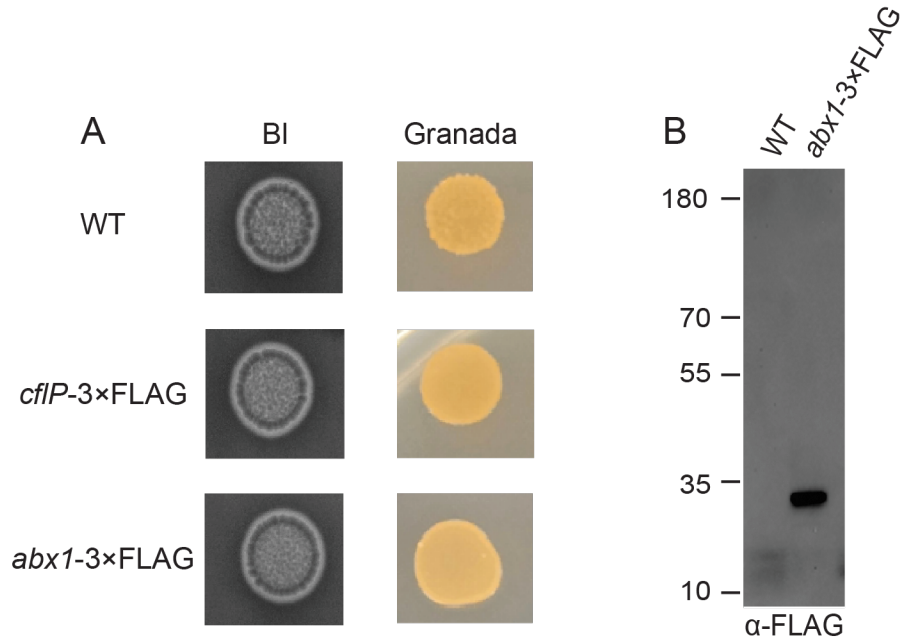

**Figure S6. Characterization of A909 strains expressing 3×FLAG-tagged CflP or Abx1 from their native chromosomal loci.**

**(A)** Pigmentation phenotypes of A909 WT and the *cflP*-3xFLAG and *abx1*-3xFLAG strains. GBS strains were grown on blood agar (left panel) or Granada agar (right panel) at 37 °C for 24 h. **(B)** Immunoblot analysis of Abx1-3×FLAG following affinity purification of membrane fractions from DSP-treated A909 WT and A909 *abx1*-3xFLAG strains. Strains were grown in THY broth. Bacterial growth, sample preparation, affinity purification, and immunoblotting with anti-FLAG antibodies were performed as described in Materials and Methods. Membrane proteins were separated on 4–12% SurePAGE™ Bis-Tris gels (GenScript) in MOPS buffer. Samples were incubated in the absence of DTT at 70 °C for 10 min before SDS-PAGE. Experiments were performed independently twice in **A** and **B**. A representative image from one experiment is shown.

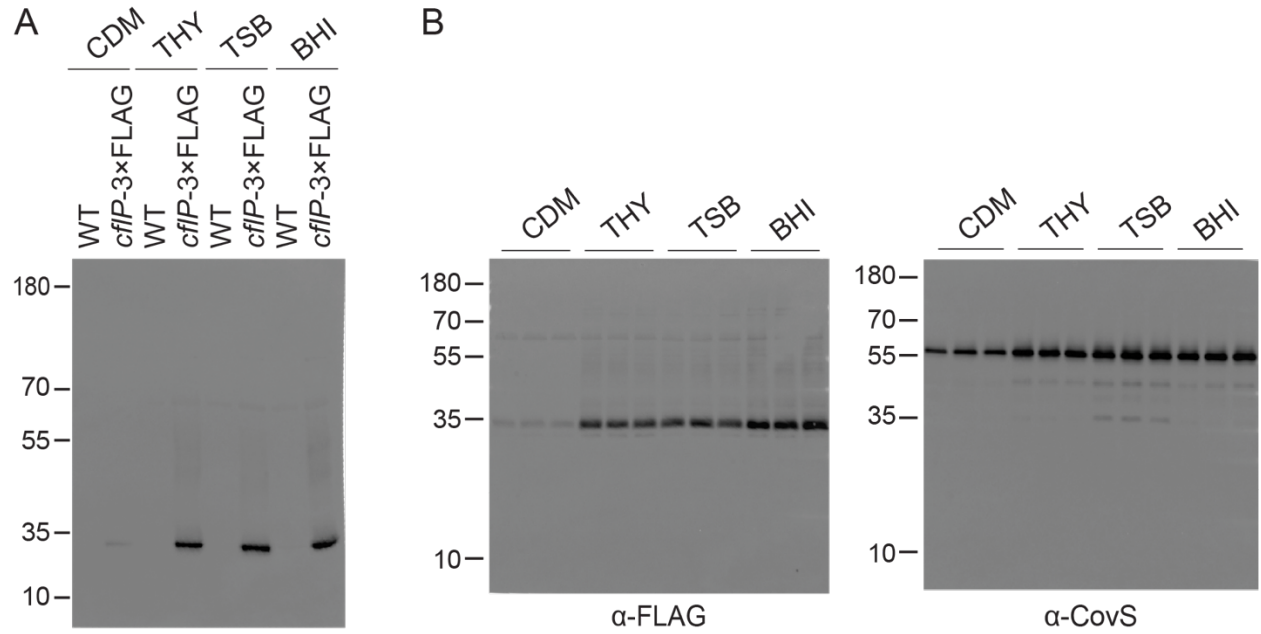

Figure S7. **CflP abundance in GBS A909 strains grown in different culture media.**

(A) Immunoblot analysis of CflP-3×FLAG in membrane fractions of mid-log-phase A909 WT (negative control) and *cflP-3×FLAG* strains grown in CDM-HighMg, THY, TSB, or BHI at 37 °C. Membrane proteins were isolated as described in Materials and Methods and separated on a 4–12% SurePAGE™ Bis-Tris gel (GenScript) in MOPS buffer. The membrane was probed with anti-FLAG antibodies. Experiments were performed independently three times. A representative image from one experiment is shown. (B) Uncropped immunoblot corresponding to Fig. 5A. Samples were separated on a 15% Bis-Tris gel. (A, B) Samples were incubated in the presence of 100 mM DTT at 70 °C for 10 min before SDS-PAGE.

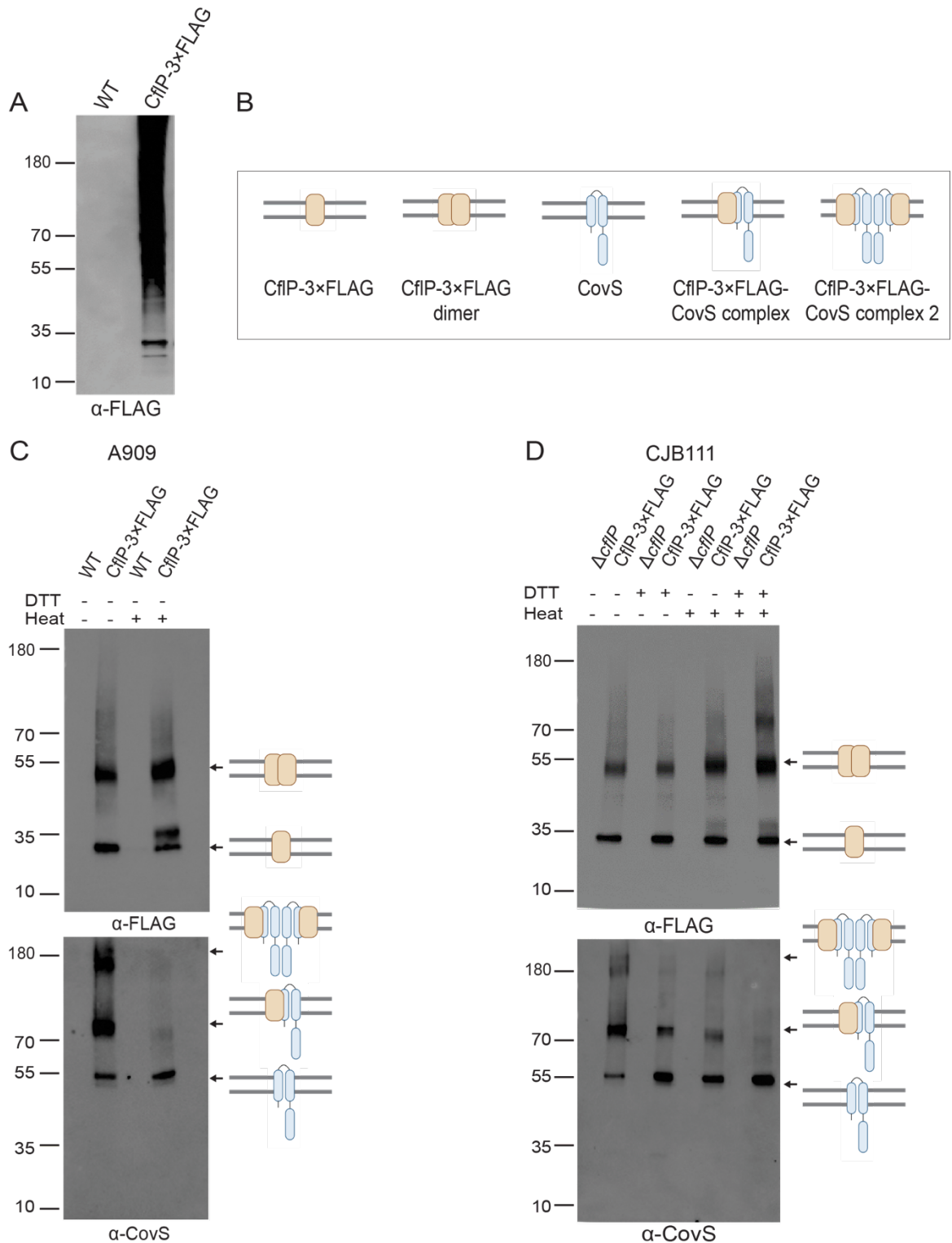

**Figure S8. Temperature-induced aggregation of CflP and analysis of the CflP–CovS interaction in A909 and CJB111.**

**(A)** Immunoblot analysis of CflP-3×FLAG after incubation at 90 °C for 10 min in the presence of 100 mM DTT. Proteins were separated on a 4–12% SurePAGE™ Bis-Tris gel (GenScript) in

MOPS buffer. **(B)** Schematic representation of potential CflP and CovS complexes, including CflP monomers and homodimers, CovS monomers, and CflP–CovS heterodimers and heterotetramers. **(C and D)** Immunoblot analysis of CflP-3×FLAG and CovS following FLAG affinity purification of membrane fractions from DSP-treated A909 WT and A909 *cflP*-3×FLAG strains **(C)** or DSP-treated CJB111Δ*cflP* strains carrying either no vector or pDC123-*cflP*-3×FLAG **(D)**. Samples were incubated with or without 100 mM DTT and either maintained at room temperature or heated at 70 °C for 10 min before SDS-PAGE. Membrane proteins were separated on 4–12% SurePAGE™ Bis-Tris gels (GenScript) in MOPS buffer. CflP-3×FLAG was detected with anti-FLAG antibodies; membranes were subsequently stripped as described in Materials and Methods and probed with anti-CovS antibodies. Experiments were performed independently three times. A representative image from one experiment is shown. Arrows indicate CflP, CovS, and putative CflP–CovS complexes in **C** and **D**.

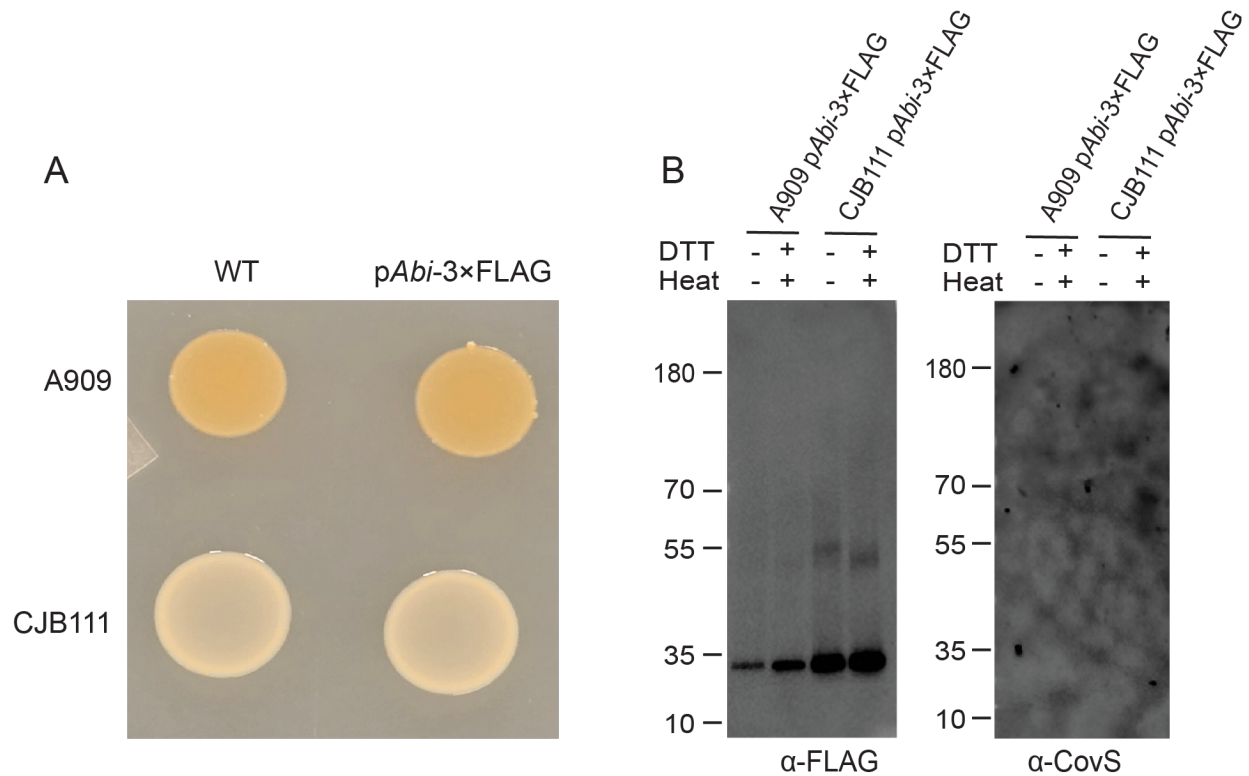

Figure S9. **CPBP *Abi* (SAK\_0054) does not regulate pigmentation or associate with CovS.** **(A)** Pigmentation phenotypes of A909 and CJB111 and their corresponding strains carrying a plasmid expressing *Abi*-3×FLAG (p*Abi*-3×FLAG). Images were taken after GBS strains were grown on Granada agar at 37 °C for 24 h. **(B)** Immunoblot analysis of *Abi*-3×FLAG and CovS following FLAG affinity purification of membrane fractions from DSP-treated A909::p*Abi*-3×FLAG and CJB111::p*Abi*-3×FLAG strains. Samples were incubated with or without 100 mM DTT and either maintained at room temperature or heated at 70 °C for 10 min before SDS-PAGE. Membrane proteins were separated on a 4–12% SurePAGE™ Bis-Tris gel (GenScript) in MOPS buffer. *Abi*-3×FLAG was detected with anti-FLAG antibodies; membranes were subsequently stripped as described in Materials and Methods and probed with anti-CovS antibodies. Experiments were performed independently twice in **A** and **B**. A representative image from one experiment is shown.

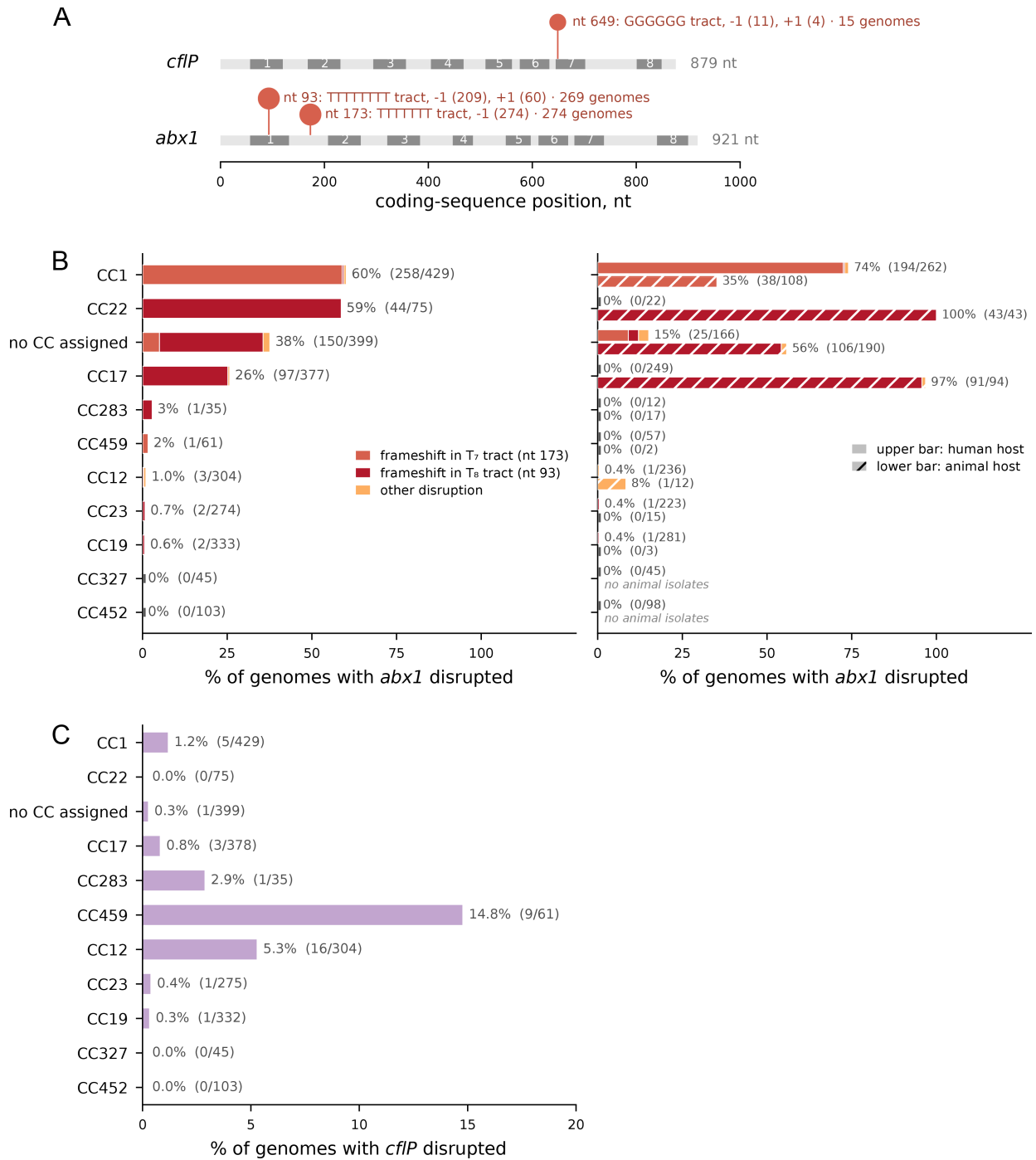

Figure S10. **Coding sequences disruptions in *abx1* and *cfIP* in GBS.**

(A) Positions of recurrent frameshift lesions in the *cfIP* and *abx1* coding sequences, for the lesions carried by at least five assemblies. The grey ribbon shows the predicted eight transmembrane helices. All frameshift lesions are listed in Table S4. (B) *abx1* disruption frequency by clonal complex, for the eleven clonal complexes with at least 30 assemblies. Every bar is partitioned by lesion: frameshift in the T<sub>7</sub> tract at coding position 173, frameshift in the T<sub>8</sub> tract at position 93, or another disruption. Left, all assemblies of each complex. Right, the same complexes split by the

assembly-annotated host: the upper bar of each pair represents human isolates (n = 1,651) and the lower hatched bar represents animal isolates (n = 484). The 300 assemblies without host annotation appear in the left sub-panel but are omitted from the right sub-panel. **(C)** *cflP* disruption frequency for the same clonal complexes. In **(B)** and **(C)**, the number of disrupted assemblies over the total number is given beside every bar, and count every non-intact class, the frameshift and other lesions. Assemblies whose locus falls on a contig edge are excluded. Clonal complexes are from MLST assignments, and “no CC assigned” represents assemblies whose sequence type cannot be mapped to any complex in the scheme.

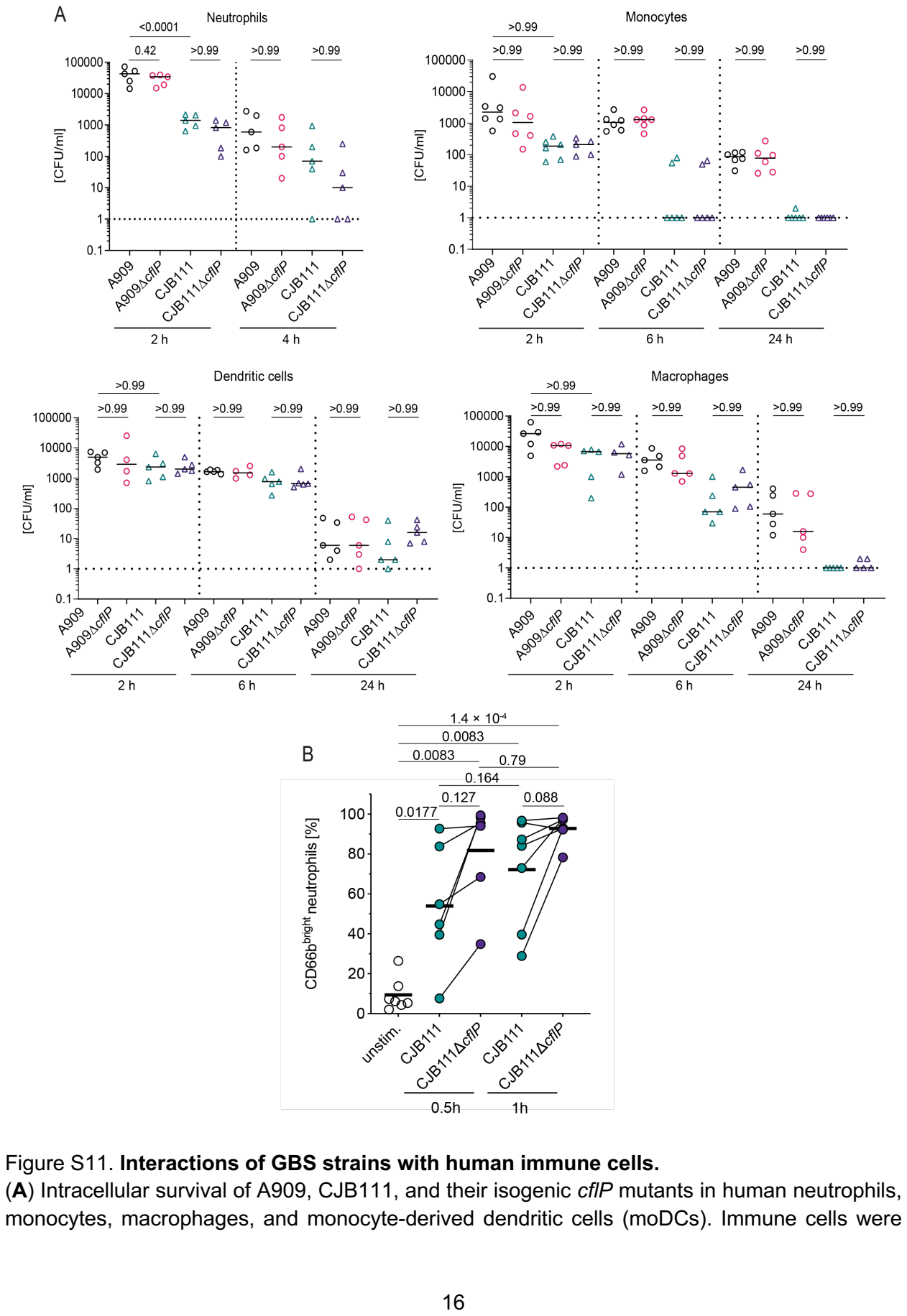

Figure S11. Interactions of GBS strains with human immune cells.

(A) Intracellular survival of A909, CJB111, and their isogenic cflP mutants in human neutrophils, monocytes, macrophages, and monocyte-derived dendritic cells (moDCs). Immune cells were infected with GBS strains at a multiplicity of infection (MOI) of 10. Extracellular bacteria were eliminated by treatment with streptomycin ($100 \mu\text{g mL}^{-1}$) and penicillin G ($100 \text{ IU mL}^{-1}$), and viable intracellular bacteria were quantified after 2 h, 6 h, and 24 h of infection. Data represent the mean ± S.D. of biologically independent replicates ($n = 5$ for neutrophils and macrophages, $n = 6$ for monocytes, $n \geq 4$ for moDC). $P$ values were determined by one-way ANOVA with Tukey's multiple-comparisons test. **(B)** Activation of neutrophils infected with GBS strains. Neutrophils were infected with GBS strains at a MOI of 10 for 0.5 and 1 h. Unstimulated cells served as negative controls. CD66b was used as a marker of activation. Each line is one biological replicate ($n \geq 6$), horizontal bar is the mean; $P$ values were calculated by paired $t$ test on logit-transformed percentages with Holm–Šidák multiple-comparison correction on pre-specified contrasts for unstimulated versus stimulated ($m = 4$) and time × genotype ($m = 4$). The gating strategy is shown in Figure S12.

A

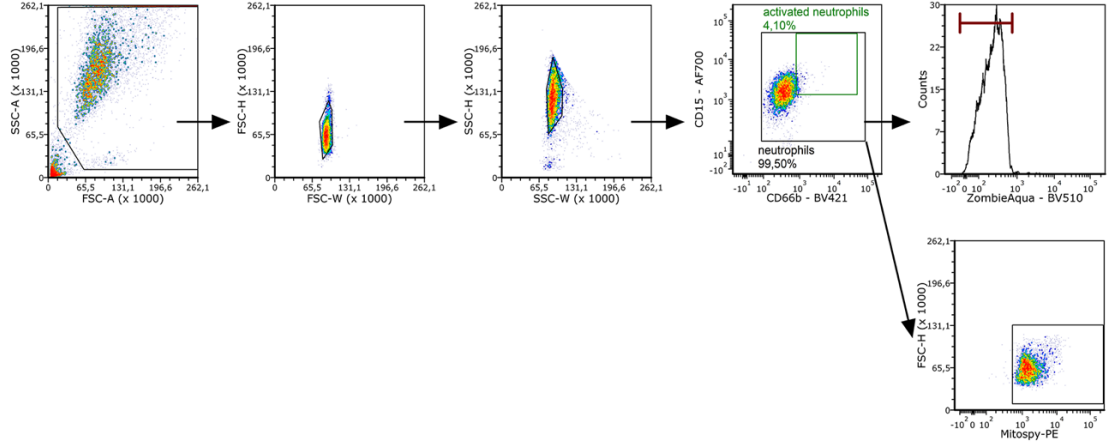

B

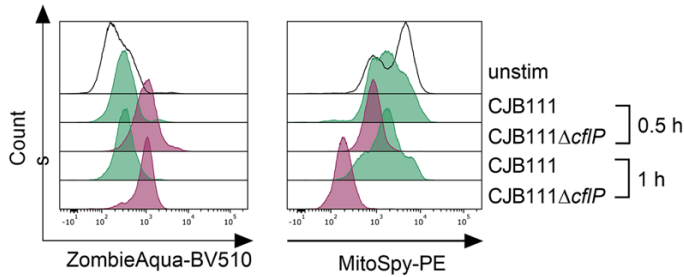

C

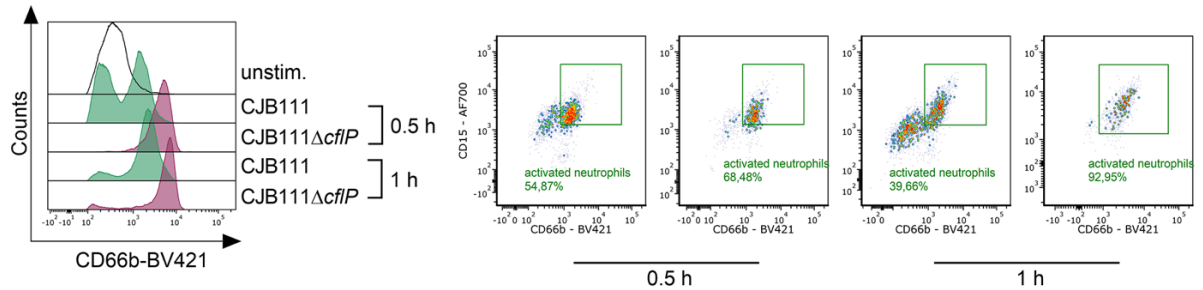

Figure S12. **Gating strategy and flow cytometric analysis of neutrophil viability, mitochondrial membrane potential, and activation.**

(A) Gating strategy for the identification of neutrophils. Doublets were excluded by consecutive gating based on FSC-H/FSC-W and SSC-H/SSC-W, and neutrophils were identified based on CD15 and CD66b expression. Activated neutrophils were defined as the CD66b<sup>bright</sup> subpopulation within the CD15<sup>+</sup>/CD66b<sup>+</sup> population. Cell viability (Zombie Aqua) and mitochondrial membrane potential (MitoSpy) were assessed within the CD15<sup>+</sup>/CD66b<sup>+</sup> population. (B) Representative histograms showing Zombie Aqua staining (viability, left) and MitoSpy staining (mitochondrial membrane potential, right) of neutrophils following infection with CJB111 or CJB111Δ*cflP* for 0.5 or 1 h. (C) Representative histograms showing CD66b expression (left) and dot plots illustrating the activated CD66b<sup>bright</sup> neutrophil population (right) following infection with CJB111 or CJB111Δ*cflP* for 0.5 or 1 h.

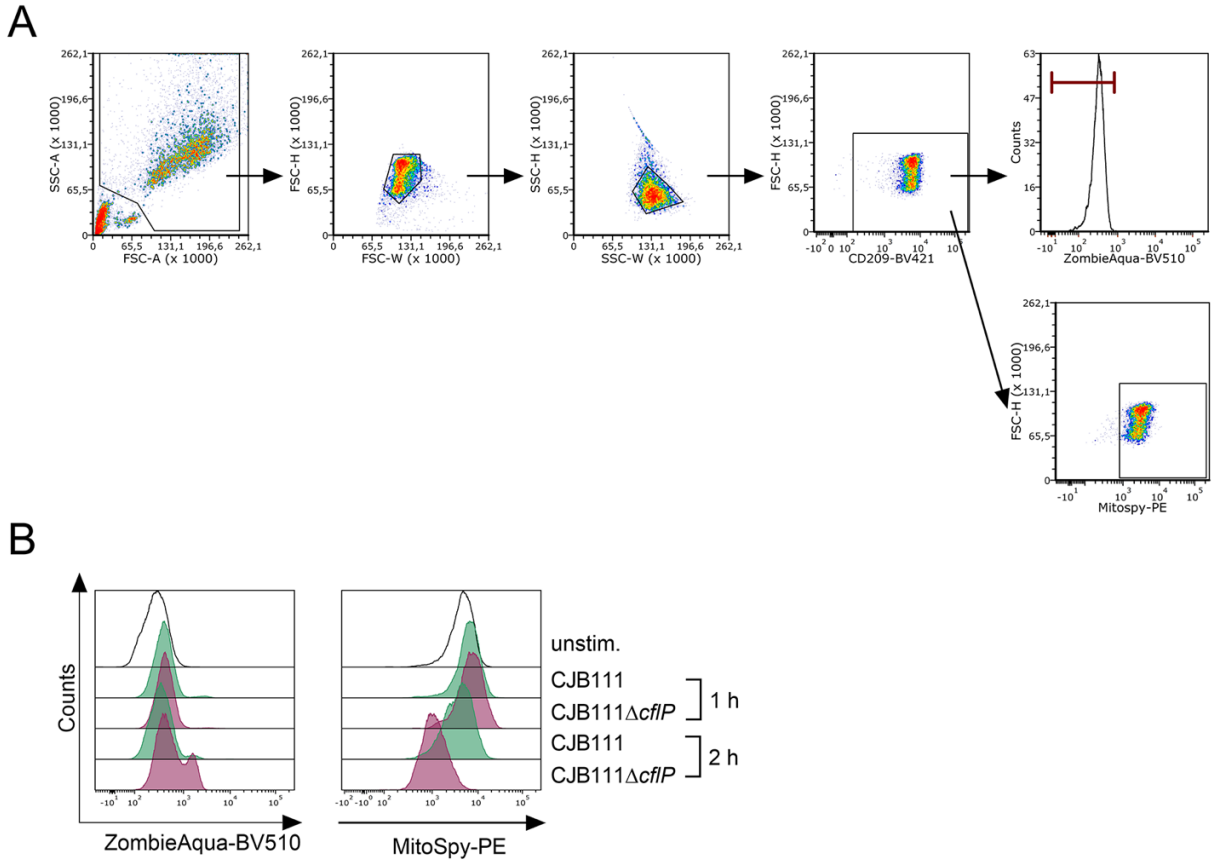

**Figure S13. Flow cytometric analysis of moDC viability and mitochondrial membrane potential.**

**(A)** Gating strategy for the identification of moDCs. Doublets were excluded by consecutive gating based on FSC-H/FSC-W and SSC-H/SSC-W, and moDCs were identified based on CD209 expression. Cell viability (Zombie Aqua) and mitochondrial membrane potential (MitoSpy) were assessed within the CD209<sup>+</sup> population. **(B)** Representative histograms showing Zombie Aqua staining (viability, left) and MitoSpy staining (mitochondrial membrane potential, right) of moDCs following infection with CJB111 or CJB111$\Delta cflP$ for 1 or 2 h.

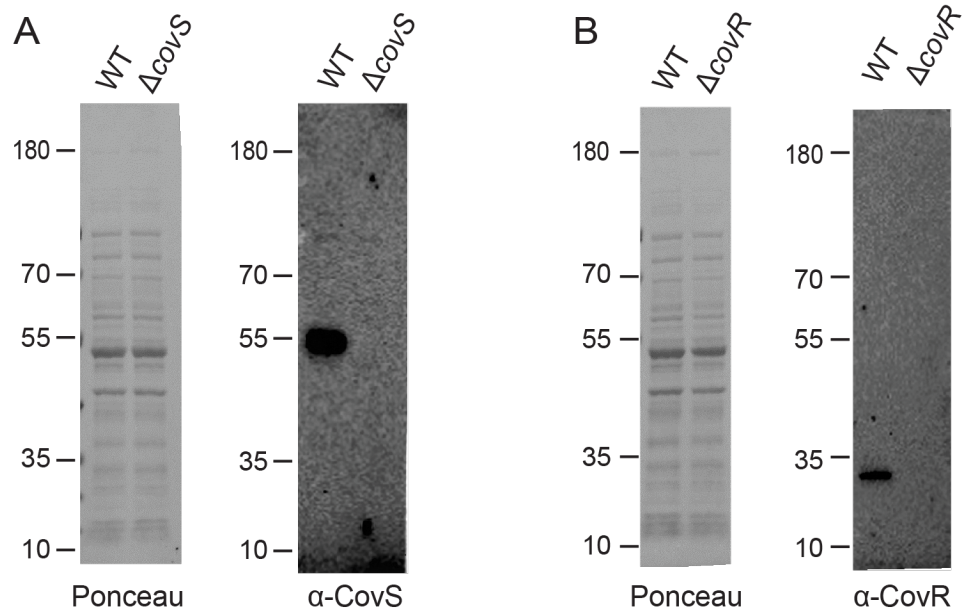

Figure S14. **Validation of anti-CovR and anti-CovS antibodies.**

(**A** and **B**) Immunoblot analysis of CovR (**A**) and CovS (**B**) in A909 WT and its isogenic $\Delta covR$ and $\Delta covS$ mutants, respectively. Proteins were separated on 4–12% SurePAGE™ Bis-Tris gels (GenScript) in MOPS buffer. The absence of the corresponding immunoreactive band in the deletion mutants confirms the specificity of the anti-CovR and anti-CovS antibodies. Experiments were performed independently three times. A representative image from one experiment is shown.

**Table S1.** Protein-level allele catalogue of CflP in GBS.

| allele | n_genomes<sup>1</sup> | freq | n_subs | n_indels | substitutions | indels |
| --- | --- | --- | --- | --- | --- | --- |
| CflP_P001 | 1528 | 0.63193 | 0 | 0 |  |  |
| CflP_P002 | 375 | 0.15509 | 1 | 0 | E3K |  |
| CflP_P003 | 138 | 0.05707 | 3 | 0 | E3K, V18I, G94R |  |
| CflP_P004 | 118 | 0.04880 | 2 | 0 | E3K, **G111E**<sup>2</sup> |  |
| CflP_P005 | 55 | 0.02275 | 6 | 0 | E3K, V18I, S149G, A150V, L193F, A219T |  |
| CflP_P006 | 27 | 0.01117 | 2 | 0 | E3K, V18I |  |
| CflP_P007 | 20 | 0.00827 | 3 | 0 | E3K, V18I, V143I |  |
| CflP_P008 | 20 | 0.00827 | 3 | 0 | E3N, V18I, G94R |  |
| CflP_P009 | 14 | 0.00579 | 1 | 0 | L201F |  |
| CflP_P010 | 12 | 0.00496 | 4 | 0 | E3K, V18I, V147L, I271T |  |
| CflP_P011 | 9 | 0.00372 | 1 | 0 | I5V |  |
| CflP_P012 | 9 | 0.00372 | 4 | 0 | E3K, V18I, V143I, V191A |  |
| CflP_P013 | 6 | 0.00248 | 1 | 0 | V58I |  |
| CflP_P014 | 5 | 0.00207 | 1 | 0 | V191I |  |
| CflP_P015 | 4 | 0.00165 | 1 | 0 | **G144E** |  |
| CflP_P016 | 3 | 0.00124 | 1 | 0 | F34S |  |
| CflP_P017 | 3 | 0.00124 | 4 | 0 | R2I, E3K, V18I, G94R |  |
| CflP_P018 | 3 | 0.00124 | 6 | 0 | E3K, V18I, V71A, V143I, R157H, M243K |  |
| CflP_P019 | 3 | 0.00124 | 3 | 0 | E3K, **G111E**, **G144E** |  |
| CflP_P020 | 3 | 0.00124 | 1 | 0 | **G106E** |  |
| CflP_P021 | 3 | 0.00124 | 1 | 0 | F264L |  |
| CflP_P022 | 2 | 0.00083 | 0 | 1 |  | del38-38<sup>3</sup> |
| CflP_P023 | 2 | 0.00083 | 3 | 0 | E3K, V18I, **I115K** |  |
| CflP_P024 | 2 | 0.00083 | 1 | 0 | G261S |  |
| CflP_P025 | 2 | 0.00083 | 2 | 0 | R2I, E3K |  |
| CflP_P026 | 2 | 0.00083 | 1 | 0 | F181L |  |
| CflP_P027 | 2 | 0.00083 | 1 | 0 | C113F |  |
| CflP_P028 | 2 | 0.00083 | 1 | 0 | V143I |  |
| CflP_P029 | 2 | 0.00083 | 5 | 0 | E3K, V18I, V71A,<br>V143I, M243K |  |
| CflP_P030 | 2 | 0.00083 | 1 | 0 | T36I |  |
| CflP_P031 | 1 | 0.00041 | 2 | 0 | E3K, P162L |  |
| CflP_P032 | 1 | 0.00041 | 1 | 0 | W282L |  |
| CflP_P033 | 1 | 0.00041 | 1 | 0 | L135M |  |
| CflP_P034 | 1 | 0.00041 | 2 | 0 | E3K, S107G |  |
| CflP_P035 | 1 | 0.00041 | 5 | 0 | E3K, V18I, **G106R**,<br>V143I, N215D |  |
| CflP_P036 | 1 | 0.00041 | 1 | 0 | A219V |  |
| CflP_P037 | 1 | 0.00041 | 3 | 0 | E3K, V18I, T131A |  |
| CflP_P038 | 1 | 0.00041 | 2 | 0 | D47E, W49R |  |
| CflP_P039 | 1 | 0.00041 | 1 | 0 | **S207L** |  |
| CflP_P040 | 1 | 0.00041 | 1 | 0 | C156S |  |
| CflP_P041 | 1 | 0.00041 | 1 | 0 | V266L |  |
| CflP_P042 | 1 | 0.00041 | 3 | 0 | E3K, V18I, A256T |  |
| CflP_P043 | 1 | 0.00041 | 3 | 0 | E3K, V18I, E241D |  |
| CflP_P044 | 1 | 0.00041 | 1 | 0 | **E267K** |  |
| CflP_P045 | 1 | 0.00041 | 1 | 0 | E3G |  |
| CflP_P046 | 1 | 0.00041 | 3 | 0 | E3K, V18I, A142T |  |
| CflP_P047 | 1 | 0.00041 | 4 | 0 | R2G, E3N, V18I,<br>G94R |  |
| CflP_P048 | 1 | 0.00041 | 4 | 0 | E3K, V18I, G94R,<br>V233I |  |
| CflP_P049 | 1 | 0.00041 | 1 | 0 | I140V |  |
| CflP_P050 | 1 | 0.00041 | 1 | 0 | G240E |  |
| CflP_P051 | 1 | 0.00041 | 2 | 0 | E3K, K213N |  |
| CflP_P052 | 1 | 0.00041 | 3 | 0 | E3K, I39V, V191I |  |
| CflP_P053 | 1 | 0.00041 | 1 | 0 | M1V |  |
| CflP_P054 | 1 | 0.00041 | 1 | 0 | E3X |  |
| CflP_P055 | 1 | 0.00041 | 1 | 0 | V71I |  |
| CflP_P056 | 1 | 0.00041 | 1 | 0 | G255D |  |
| CflP_P057 | 1 | 0.00041 | 2 | 0 | E3K, R101I |  |
| CflP_P058 | 1 | 0.00041 | 4 | 0 | E3K, V18I, T131A, G214C |  |
| CflP_P059 | 1 | 0.00041 | 1 | 0 | V58F |  |
| CflP_P060 | 1 | 0.00041 | 1 | 0 | A256T |  |
| CflP_P061 | 1 | 0.00041 | 1 | 0 | G218E |  |
| CflP_P062 | 1 | 0.00041 | 1 | 0 | R83H |  |
| CflP_P063 | 1 | 0.00041 | 4 | 0 | E3K, V18I, G94R, L283I |  |
| CflP_P064 | 1 | 0.00041 | 1 | 0 | G255S |  |
| CflP_P065 | 1 | 0.00041 | 5 | 0 | E3K, V18I, V147L, I271T, P290S |  |
| CflP_P066 | 1 | 0.00041 | 2 | 0 | E3K, L35V |  |
| CflP_P067 | 1 | 0.00041 | 6 | 0 | E3K, V18I, C113Y, V147L, V238A, I271T |  |
| CflP_P068 | 1 | 0.00041 | 1 | 0 | **G261D** |  |
| CflP_P069 | 1 | 0.00041 | 1 | 0 | A225T |  |
| CflP_P070 | 1 | 0.00041 | 2 | 0 | R2I, S149G |  |
| CflP_P071 | 1 | 0.00041 | 2 | 0 | E3K, V77L |  |
| CflP_P072 | 1 | 0.00041 | 0 | 1 |  | I40dup<sup>3</sup> |

<sup>1</sup> **n\_genomes** is the number of assemblies carrying the allele and **freq** is its fraction of the 2,418 intact copies. **substitutions** lists amino-acid changes relative to A909 reference (CflP\_P001) in A909 numbering; **indels** uses *delstart–end* for residues absent from the reference and **dup** for a residue duplicated relative to it.

<sup>2</sup> Substitution predicted to have the most deleterious effect, with a masked-marginal log-likelihood ratio $\leq -7$, are highlighted in **bold**.

<sup>3</sup> Two alleles differ from the reference in length rather than in sequence, and both involve the same (ATT)<sub>3</sub> tandem repeat encoding Ile38–Ile40. CflP\_P022 (291 aa, del38–38) removes one isoleucine and is present in two assemblies (GCF\_001592425.1, GCF\_014875205.1). CflP\_P072 (293 aa, I40dup) adds one isoleucine and is present in a single assembly (GCF\_043949215.1). Short tandem repeats are a source of indel error in short-read assemblies, and all three genomes were sequenced on Illumina platforms, so the $\pm 1$ isoleucine alleles should be treated as provisional.

**Table S2.** Substitutions predicted to have deleterious effects on protein structure and function.

| protein | substitution | position | segment | assemblies | % of intact copies | ESMC-600M LLR | landscape percentile |
| --- | --- | --- | --- | --- | --- | --- | --- |
| CflP | G106R | 106 | TM3 | 1 | 0.04 | -10.59 | 4.0 |
| CflP | S207L | 207 | TM6 | 1 | 0.04 | -9.81 | 6.6 |
| CflP | G106E | 106 | TM3 | 3 | 0.12 | -9.69 | 7.1 |
| CflP | I115K | 115 | TM3 | 2 | 0.08 | -9.18 | 9.1 |
| CflP | G261D | 261 | extracellular loop | 1 | 0.04 | -8.68 | 12.0 |
| CflP | G144E | 144 | TM4 | 7 | 0.29 | -8.00 | 15.6 |
| CflP | E267K | 267 | extracellular loop | 1 | 0.04 | -7.77 | 17.2 |
| CflP | **G111E** | 111 | TM3 | **121** | **5.00** | -7.43 | 19.4 |
| CflP | F181L | 181 | TM5 | 2 | 0.08 | -6.93 | 22.5 |
| CflP | M1V | 1 | N-terminal segment, cytoplasmic | 1 | 0.04 | -6.83 | 23.4 |
| CflP | R83H | 83 | cytoplasmic loop | 1 | 0.04 | -6.81 | 23.5 |
| CflP | G261S | 261 | extracellular loop | 2 | 0.08 | -6.68 | 24.3 |
| CflP | R157H | 157 | cytoplasmic loop | 3 | 0.12 | -6.55 | 25.3 |
| CflP | G214C | 214 | cytoplasmic loop | 1 | 0.04 | -6.53 | 25.4 |
| CflP | F264L | 264 | extracellular loop | 3 | 0.12 | -6.29 | 27.1 |
| CflP | V238A | 238 | extracellular loop | 1 | 0.04 | -6.13 | 28.2 |
| CflP | P162L | 162 | cytoplasmic loop | 1 | 0.04 | -5.10 | 35.5 |
| Abx1 | W157G | 157 | TM4 | 2 | 0.11 | -10.86 | 6.4 |
| Abx1 | T168R | 168 | cytoplasmic loop | 1 | 0.05 | -7.05 | 23.6 |
| Abx1 | K224E | 224 | cytoplasmic loop | 8 | 0.42 | -5.29 | 34.3 |

The table lists all observed substitution with a masked-marginal log-likelihood ratio $\leq -5$.

**Landscape percentile** is the position within that protein's complete ESMC-600M substitution landscape, all positions × 19 substitutions.

**Table S3.** Protein-level allele catalogue of Abx1 in GBS.

| allele | n_genomes<sup>1</sup> | freq | n_subs | substitutions |
| --- | --- | --- | --- | --- |
| Abx1_P001 | 1301 | 0.68582 | 0 |  |
| Abx1_P002 | 243 | 0.12810 | 1 | T107I |
| Abx1_P003 | 74 | 0.03901 | 1 | L135I |
| Abx1_P004 | 63 | 0.03321 | 1 | D270N |
| Abx1_P005 | 60 | 0.03163 | 1 | F60L |
| Abx1_P006 | 26 | 0.01371 | 2 | L135I, H202R |
| Abx1_P007 | 26 | 0.01371 | 1 | N227D |
| Abx1_P008 | 12 | 0.00633 | 1 | N15D |
| Abx1_P009 | 11 | 0.00580 | 1 | L289F |
| Abx1_P010 | 10 | 0.00527 | 2 | L135I, I195V |
| Abx1_P011 | 9 | 0.00474 | 1 | I195V |
| Abx1_P012 | 8 | 0.00422 | 2 | I195V, **K224E<sup>2</sup>** |
| Abx1_P013 | 8 | 0.00422 | 1 | T168I |
| Abx1_P014 | 5 | 0.00264 | 1 | A255V |
| Abx1_P015 | 4 | 0.00211 | 1 | G256K |
| Abx1_P016 | 4 | 0.00211 | 2 | L135I, L296F |
| Abx1_P017 | 3 | 0.00158 | 1 | Q253H |
| Abx1_P018 | 3 | 0.00158 | 1 | Y13C |
| Abx1_P019 | 3 | 0.00158 | 1 | L158I |
| Abx1_P020 | 3 | 0.00158 | 1 | A25T |
| Abx1_P021 | 2 | 0.00105 | 1 | F60X |
| Abx1_P022 | 2 | 0.00105 | 1 | L234V |
| Abx1_P023 | 2 | 0.00105 | 1 | **W157G** |
| Abx1_P024 | 2 | 0.00105 | 1 | L84F |
| Abx1_P025 | 1 | 0.00053 | 2 | I211V, D270N |
| Abx1_P026 | 1 | 0.00053 | 1 | S110L |
| Abx1_P027 | 1 | 0.00053 | 1 | S241I |
| Abx1_P028 | 1 | 0.00053 | 1 | L23F |
| Abx1_P029 | 1 | 0.00053 | 2 | S214L, A255V |
| Abx1_P030 | 1 | 0.00053 | 1 | A201V |
| Abx1_P031 | 1 | 0.00053 | 2 | P174H, I195V |
| Abx1_P032 | 1 | 0.00053 | 1 | C26Y |
| Abx1_P033 | 1 | 0.00053 | 2 | T107I, W298L |
| Abx1_P034 | 1 | 0.00053 | 1 | E280X |
| Abx1_P035 | 1 | 0.00053 | 2 | P17L, T107I |
| Abx1_P036 | 1 | 0.00053 | 1 | L245F |
| Abx1_P037 | 1 | 0.00053 | 1 | **T168R** |

<sup>1</sup>**n\_genomes** is the number of assemblies carrying the allele and **freq** is its fraction of the 1,897 intact copies. **substitutions** list amino-acid changes relative to A909 reference (Abx1\_P001) in A909 numbering.

<sup>2</sup>Substitution predicted to have the most deleterious effect, with a masked-marginal log-likelihood ratio $\leq -5$, are highlighted in **bold**.

**Table S4.** Frameshift lesions in *abx1* and *cflP* in GBS.

| gene | frameshifts | n_genomes | product_aa | ref_nt_pos | indel | homopolymer | n_CC | n_genomes_no_CC |
| --- | --- | --- | --- | --- | --- | --- | --- | --- |
| *abx1* | refnt173:-1 | 274 | 72 | 173 | -1 | TTTTTTT | 2 | 20 |
| *abx1* | refnt93:-1 | 209 | 36 | 93 | -1 | TTTTTTT | 6 | 66 |
| *abx1* | refnt93:+1 | 60 | 52 | 93 | 1 | TTTTTTT | 3 | 56 |
| *abx1* | refnt511:-1 | 2 | 174 | 511 | -1 | GGG | 0 | 2 |
| *abx1* | refnt172:-1 | 1 | 72 | 172 | -1 | TTTTTTT | 1 | 0 |
| *abx1* | refnt271:-1 | 1 | 111 | 271 | -1 | AAAAAA | 0 | 1 |
| *abx1* | refnt478:-1 | 1 | 166 | 478 | -1 | AAA | 0 | 1 |
| *abx1* | refnt68:-1 | 1 | 36 | 68 | -1 | TTT | 0 | 1 |
| *cflP* | refnt649:-1 | 11 | 247 | 649 | -1 | GGGGGG | 4 | 0 |
| *cflP* | refnt649:+1 | 4 | 238 | 649 | 1 | GGGGGG | 2 | 0 |
| *cflP* | refnt708:-10 | 2 | 244 | 708 | -10 | TTT | 1 | 0 |
| *cflP* | refnt13:-1 | 1 | 19 | 13 | -1 | TTTTT | 1 | 0 |
| *cflP* | refnt556:+2 | 1 | 191 | 556 | 2 |  | 1 | 0 |
| *cflP* | refnt564:-1 | 1 | 190 | 564 | -1 |  | 1 | 0 |
| *cflP* | refnt695:+1 | 1 | 238 | 695 | 1 | GGG | 1 | 0 |
| *cflP* | refnt805:-4 | 1 | 276 | 805 | -4 |  | 1 | 0 |
| *cflP* | refnt853:-1 | 1 | 309 | 853 | -1 | AAAAA | 1 | 0 |

Each row is one distinct lesion, defined by its position in the A909 coding sequence and the signed length of the indel, ordered by carrier count within each gene. **frameshifts** is the lesion identifier, refntposition:±n (for example refnt173:-1 denotes a single-base deletion at coding-sequence position 173); **ref\_nt\_pos** and **indel** give the same two quantities as separate numeric columns. **n\_genomes** is the number of assemblies carrying the lesion. **product\_aa** is the length in residues of the predicted product of the frameshifted allele. **homopolymer** is the homopolymeric tract present at that position in the reference sequence, and is left blank for the three *cflP* lesions that do not fall in one. **n\_CC** counts the named clonal complexes among carriers and **n\_genomes\_no\_CC** the carriers with no MLST assignment, which are excluded from **n\_CC**. The table accounts for all 23 frameshifted *cflP* assemblies and 549 of the 551 frameshifted *abx1* assemblies; the two remaining *abx1* genomes (GCF\_014875555.1 and GCF\_053545915.1) are contig-level assemblies whose 5' end lies at a contig boundary, so although an indel is detectable in the alignment near position 173, the start codon cannot be located and no reference coordinate can be assigned.

**Table S5.** Strains and plasmids used in this study.

| Strain or plasmid | Description <sup>a</sup> | Reference |
| --- | --- | --- |
| *Streptococcus agalactiae* |  |  |
| **A909 serotype** |  |  |
| A909 | Serotype Ia strain, wild-type (WT) | ATCC |
| A909Δ*cflP* | *cflP* deletion mutant (has a nonpolar spectinomycin resistance cassette inserted in *cflP*), Spec<sup>R</sup> | This study |
| A909Δ*cflP*::*pcfIP* | Δ*cflP* is complemented with *pcfIP* carrying WT *cflP*, Spec<sup>R</sup>, Cm<sup>R</sup> | This study |
| Δ*abx1* | *abx1* deletion mutant (has an unmarked deletion of *abx1*) | This study |
| Δ*abx1*::*pabx1* | Δ*abx1* is complemented with *pabx1* carrying WT *abx1*, Erm<sup>R</sup> | This study |
| Δ*covR* | *covR* deletion mutant (has an unmarked deletion of *covR*) | This study |
| Δ*covS* | *covS* deletion mutant (has an unmarked deletion of *covS*) | This study |
| Δ*covS*Δ*cflP* | *covS cflP* double-gene deletion mutant (has nonpolar spectinomycin resistance cassette inserted in *cflP* and a deletion of *covS*), Spec<sup>R</sup> | This study |
| Δ*covR*Δ*cflP* | *covR cflP* double-gene deletion mutant (has nonpolar spectinomycin resistance cassette inserted in *cflP* and a deletion of *covS*), Spec<sup>R</sup> | This study |
| Δ*covS*Δ*cflP*::*pcfIP* | Δ*covS* Δ*cflP* is complemented with *pcfIP* carrying WT *cflP*, Spec<sup>R</sup>, Cm<sup>R</sup> | This study |
| Δ*abx1*Δ*cflP* | *abx1 cflP* double-gene deletion mutant (has nonpolar spectinomycin resistance cassette inserted in *cflP* and a nonpolar deletion of *abx1*), Spec<sup>R</sup> | This study |
| Δ*abx1*Δ*cflP*::*pcfIP* | Δ*abx1*Δ*cflP* is complemented with *pcfIP* carrying WT *cflP*, Spec<sup>R</sup>, Cm<sup>R</sup> | This study |
| Δ*covS*Δ*abx1* | *covS abx1* double-gene deletion mutant (has nonpolar deletion of *covS* and *abx1*) | This study |
| Δ*covR*Δ*abx1* | *covR abx1* double-gene deletion mutant (has nonpolar deletion of *covS* and *abx1*) | This study |
| *cflP*-3×FLAG | *cflP* with a 3×FLAG tag in its C-terminal | This study |
| *abx1*-3×FLAG | *cflP* with a 3×FLAG tag in its C-terminal | This study |
| Δ*bac* | *bac* deletion mutant in A909 background | (3) |
| A909Δ*cflP*::*pcfIP*<sup>A909</sup>-3×FLAG | Δ*cflP* carrying *pcfIP*<sup>A909</sup>-3×FLAG, Spec<sup>R</sup>, Cm<sup>R</sup> | This study |
| A909Δ*cflP*::*pcfIP*<sup>NEM316</sup>-3×FLAG | Δ*cflP* carrying *pcfIP*<sup>NEM316</sup>-3×FLAG, Spec<sup>R</sup>, Cm<sup>R</sup> | This study |
| A909::p*Abi*-3×FLAG | A909 carrying p*Abi*-3×FLAG plasmid, Cm<sup>R</sup> | This study |
| A909Δ*cflP*::*pcfIP*<sup>H185A</sup> | A909Δ*cflP* is complemented with *pcfIP*<sup>H185A</sup>, Spec<sup>R</sup>, Cm<sup>R</sup> | This study |
| A909Δ*cflP*::*pcfIP*<sup>H223A</sup> | A909Δ*cflP* is complemented with *pcfIP*<sup>H223A</sup>, Spec<sup>R</sup>, Cm<sup>R</sup> | This study |
| **CJB111 serotype** |  |  |
| CJB111 | Serotype V strain, wild-type (WT) | (4) |
| CJB111Δ*cflP* | *cflP* deletion mutant (has a nonpolar spectinomycin resistance cassette inserted in *cflP*), Spec<sup>R</sup> | This study |
| CJB111Δ*cflP*::*pcfIP* | CJB111Δ*cflP* is complemented with *pcfIP* carrying WT *cflP*, Spec<sup>R</sup>, Cm<sup>R</sup> | This study |
| CJB111Δ*cflP*::*pcfIP*-3×FLAG | CJB111Δ*cflP* is complemented with *pcfIP*-3×FLAG carrying *cflP*-3×FLAG, Spec<sup>R</sup>, Cm<sup>R</sup> | This study |
| CJB111Δ*cflP*::*pcfIP*<sup>A909</sup>-3×FLAG | CJB111Δ*cflP* carrying *pcfIP*<sup>A909</sup>-3×FLAG, Spec<sup>R</sup>, Cm<sup>R</sup> | This study |
| CJB111Δ*cflP*::*pcfIP*<sup>NEM316</sup>-3×FLAG | CJB111Δ*cflP* carrying *pcfIP*<sup>NEM316</sup>-3×FLAG, Spec<sup>R</sup>, Cm<sup>R</sup> | This study |
| CJB111::p*Abi*-3×FLAG | CJB111 carrying p*Abi*-3×FLAG plasmid, Cm<sup>R</sup> | This study |
| CJB111Δ*cflP*::*pcfIP*<sup>H185A</sup> | CJB111Δ*cflP* is complemented with *pcfIP*<sup>H185A</sup>, Spec<sup>R</sup>, Cm<sup>R</sup> | This study |
| CJB111Δ*cflP*::*pcfIP*<sup>H223A</sup> | CJB111Δ*cflP* is complemented with *pcfIP*<sup>H223A</sup>, Spec<sup>R</sup>, Cm<sup>R</sup> | This study |
| Tn::*cflP* | Transposon insertion in *cflP* in CJB111, Kan<sup>R</sup> | This study |
| **NEM316 serotype** |  |  |
| NEM316 | Serotype III strain, wild-type (WT) | (5) |
| NEM316::*pcfIP*<sup>A909</sup>-3×FLAG | NEM316 carrying *pcfIP*<sup>A909</sup>-3×FLAG, Cm<sup>R</sup> | This study |
| NEM316::*pcfIP*<sup>NEM316</sup>-3×FLAG | NEM316 carrying *pcfIP*<sup>NEM316</sup>-3×FLAG, Cm<sup>R</sup> | This study |
| **Escherichia coli** |  |  |
| DH5α | *E. coli* cells used for cloning | Invitrogen |
| LOBSTR(DE3)::pRARE2 | *E. coli* strain with rare-codon supplementation from the pRARE2 plasmid, used for protein expression; Cm<sup>R</sup> | Kerafast |
| **Plasmids** |  |  |
| pDC123 | *E. coli-streptococcus* shuttle vector, JS-3 replicon, chloramphenicol resistance cassette; Cm<sup>R</sup> | (6) |
| pDC123-CflP (*pcfIP*) | A pDC123 derived plasmid expressing *cflP*, Cm<sup>R</sup> | This study |
| pDC123-CflP<sup>H185A</sup> (*pcfIP*<sup>H185A</sup>) | A pDC123 derived plasmid expressing *cflP*<sup>H185A</sup>, Cm<sup>R</sup> | This study |
| pDC123-CflP<sup>H223A</sup> (*pcfIP*<sup>H223A</sup>) | A pDC123 derived plasmid expressing *cflP*<sup>H223A</sup>, Cm<sup>R</sup> | This study |
| *pcfIP*-3×FLAG | A pDC123 derived plasmid expressing *cflP*-3×FLAG, Cm<sup>R</sup> | This study |
| *pcfIP*<sup>PG105E</sup>-3×FLAG | A pDC123 derived plasmid expressing *cflP*<sup>PG105E</sup>-3×FLAG, Cm<sup>R</sup> | This study |
| p*Abi*-3×FLAG | A pDC123 derived plasmid expressing *Abi*-3×FLAG (*sak_0054*), Cm<sup>R</sup> | This study |
| pJLL286 | a low-copy-number vector with a nitrate-inducible promoter, Erm<sup>R</sup> | <sup>80</sup> |
| *pabx1* | A pJLL286 derived plasmid expressing *abx1*, Erm<sup>R</sup> | This study |
| pCOLA-Duet1 | Expression vector for *E. coli* | Millipore |
| pCOLA-covS | A pCOLA-Duet1 derived plasmid expressing *covS*, Kan<sup>R</sup> | This study |
| pCOLA-cflP-3×FLAG | A pCOLA-Duet1 derived plasmid expressing *cflP*-3×FLAG, Kan<sup>R</sup> | This study |
| pCOLA-covS-*cflP*-3×FLAG | A pCOLA-Duet1 derived plasmid expressing *covS* and *cflP*-3×FLAG, Kan<sup>R</sup> | This study |
| pMBsacB | A temperature-sensitive shuttle vector for in-frame deletions, Erm<sup>R</sup> | (7) |
| pMBsacB-Δ*cflP* | A pMBsacB derived plasmid to construct Δ*cflP*::Sp<sup>R</sup>, Erm<sup>R</sup> | This study |
| pLT06 | Temperature-sensitive shuttle vector used for gene deletion and genomic modification in *Enterococcus* species | (8) |
| pLT06-Δ*covR* | A pLT06 derived plasmid to construct Δ*covR*, Cm<sup>R</sup> | This study |
| pLT06-Δ*covS* | A pLT06 derived plasmid to construct Δ*covS*, Cm<sup>R</sup> | This study |
| pLT06-Δ*abx1* | A pLT06 derived plasmid to construct Δ*abx1*, Cm<sup>R</sup> | This study |
| pRSF-NT | A modified pRSF-Duet1 (Novagen) vector for making N-terminus His-tagged proteins with a TEV protease cleavage site; Kan<sup>R</sup> | (9) |
| pRSF-covR | A pRSF-NT derived plasmid expressing *covR* for its overexpression and purification, Kan<sup>R</sup> | This study |
| pRSF-covS | A pRSF-NT derived plasmid expressing *covS* for its overexpression and purification, Kan<sup>R</sup> | This study |

<sup>a</sup> Antibiotic resistance markers: Erm<sup>R</sup>, erythromycin; Kan<sup>R</sup>, kanamycin; Spec<sup>R</sup>, spectinomycin; Cm<sup>R</sup>, chloramphenicol,

**Table S6.** Primers used in this study.

| Name | Description | Sequence (5' to 3') |
| --- | --- | --- |
| CaaX-EcoRI-F | Primer used to amplify *cflP* and add a 5' EcoRI site for insertion into pDC123 vector | GCTTCGAATTCGTATAAAGTTAGATAAAA<br>GTGAG |
| CaaX-BamHI-R | Primer used to amplify *cflP* and add a 3' BamHI site for insertion into pDC123 vector | CGTCTGGATCCGATTAAATCTATTAGTC<br>GTAAGG |
| CaaX-3xFLAG-BamH | Primer used to amplify *cflP* with a C-terminal 3xFLAG sequence and add a 3' BamHI site for insertion into pDC123 vector. Used with CaaX-EcoRI-F to construct pDC123-*cflP*-3xFLAG | CCTGGATCCTTACTTGTCATCGTCATCC<br>TTGTAATCGATATCATGATCTTTATAATC<br>ACCGTCATGGTCTTTGTAGTCGTCGTAA<br>GGAAAATTCGATTTTTG |
| CaaX_H185A_F | Primer used with CaaX_H185A_R to construct CflP<sup>H185A</sup> | CTTTTTCGGTCTTCTGGCTAGCGCTAAT<br>AATC |
| CaaX_H185A_R | Primer used with CaaX_H185A_F to construct CflP<sup>H185A</sup> | GATTATTAGCGCTAGCCAGAAGACCGAA<br>AAAG |
| CaaX_H223A_F | Primer used with CaaX_H223A_R to construct CflP<sup>H223A</sup> | GCGTGTGGTATTGCTGGCGCCTGGAAC |
| CaaX_H223A_R | Primer used with CaaX_H223A_F to construct CflP<sup>H223A</sup> | GTTCCAGGCGCCAGCAATACCACACGC |
| cflP-G106E-F | Primer used with cflP-G106E-R to construct CflP<sup>NEM316</sup> | TGGTTAGTCGAGAGCGTTCTCTTTGGAG |
| cflP-G106E-R | Primer used with cflP-G106E-F to construct CflP<sup>NEM316</sup> | AGAACGCTCTCGACTAACCATCCTCTAG |
| Abx1-BamH | Primer used to amplify *abx1* and add a 5' BamHI site for insertion into pJLL286 vector | CTACAAAATAAATTATGGATCCACACAAA<br>AAGGATTTAGG |
| Abx1-Xho | Primer used to amplify *abx1* and add a 3' XhoI site for insertion into pJLL286 vector | CTATAGGGAGACCGGCCTCGAGCTCAA<br>TCTGTTTCTTCAGTC |
| covS-Nde | Primer used to amplify *covS* and add a 5' NdeI site for insertion into pCOLA-Duet1 vector | AAGTATAAGAAGGAGATATACATATGAA<br>AAATAAAAAAGATCAGTTTATAG |
| covS-Xho | Primer used to amplify *covS* and add a 3' XhoI cut site for insertion into pCOLA-Duet1 vector | CAGCGGTTTCTTTACCAGACTCGAGCTT<br>ATATTTCTTTAGTTTCTTC |
| cflP-BspH | Primer used to amplify *cflP* and add a 5' BspHI site for insertion into pCOLA-Duet1 vector | GTTTAACTTTAATAAGGAGATATATCATG<br>AGAGAGTCTATTTTTGAG |
| cflP-3xFLAG-Hind | Primer used to amplify *cflP*-3xFLAG and add a 3' HindIII site for insertion into pCOLA-Duet1 vector | TTAAGCATTATGCGGCCGCAAGCTTACT<br>TGTCATCGTCATCC |
| CaaX-XhoI-f | Primer used with Spec-CaaX-r1 to amplify arm 1 (731 bp upstream *cflP*) and add a 5' XhoI site for insertion into pMBSacB vector | GTGCGCTCGAGTCAATGCGAATGTTGG<br>TC |
| Spec-CaaX-r1 | Primer used with CaaX-XhoI-f to amplify arm 1 (731 bp upstream *cflP*) with a short homology sequence to CaaX-Spec-f1 | CTCACTATTTTGGTCGACTCTCTCATAAA<br>TATCTC |
| CaaX-Spec-f1 | Primer used with CaaX-Spec-r2 to amplify spectinomycin cassette flanked by short homology arms to Spec-CaaX-r1 | TTTATGAGAGAGTCGACCAAAATAGTGA<br>GGAG |
| CaaX-Spec-r2 | Primer used with CaaX-Spec-f1 to amplify spectinomycin cassette flanked by short homology arms to Spec-CaaX-f2 | TCATGCTTCCTTCCGATCCTTATAATTTT<br>TTTAATCTG |
| Spec-CaaX-f2 | Primer used with CaaX-BamH-r to amplify arm 2 (765 bp downstream *cflP*) with a short homology sequence to CaaX-Spec-r2 | AAAATTATAAGGATCGGAAGGAAGCATG<br>ATTACCAG |
| CaaX-BamH-r | Primer used with Spec-CaaX-f2 to amplify arm 2 (765 bp downstream *cflP*) and add a 3' BamHI site for insertion into pMBSacB vector | CGTCTGGATCCGTCTGATGTCATTTTGG<br>ATAC |
| CaaX-check-f | Primer used with CaaX-check-r to check $\Delta cflP$ mutation | CGATTGACAGCAAGTAATC |
| CaaX-check-r | Primer used with CaaX-check-f to check $\Delta cflP$ mutation | GATAAAGCCAATAACATGCTAC |
| covR-EcoR-F | Primer used with covR-R1 to amplify arm 1 (907 bp downstream *covR*) and add a 5' EcoRI site for insertion into pLT06 vector | ATCCCCGGGTACCGAGCTCGAATTCTTT<br>ACAACAGCGACAGG |
| covR-R1 | Primer used with covR-EcoR-F to amplify arm 1 (907 bp upstream *covR*) with short homology sequence to covR-F1 | GTTGGTGTAGAGAATCATATATTCAAAC<br>TGTTCCG |
| covR-F1 | Primer used with covR-BamH-R to amplify arm 2 (907 bp upstream *covR*) with short homology sequence to covR-R1 | GAATATATGATTCTCTACACCAACCTTCT<br>TATTC |
| covR-BamH-R | Primer used with covR-F1 to amplify arm 2 (907 bp upstream *covR*) and add a 3' BamHI site for insertion into pLT06 vector | CCTGCAGGTCGACTCTAGAGGATCCTG<br>CAGATCGTATTGAGCG |
| covR-chF | Primer used with covR-chF to check $\Delta covR$ mutation | TTTTTCTAATGACCTACCGAC |
| covR-chR | Primer used with covR-chR to check $\Delta covR$ mutation | AAGGTATGTTTGATAGTGATTAG |
| covS-EcoR-F | Primer used with covS-R1 to amplify arm 2 (913 bp downstream *covS*) and add a 5' EcoRI site for insertion into pLT06 vector | ATCCCCGGGTACCGAGCTCGAATTCTCA<br>ACTTAACAAGGTAAGTAC |
| covS-R1 | Primer used with covS-EcoR-F to amplify arm 2 (913 bp downstream *covS*) | AGGGGTAAAACTAAGAAAAATCCTTATG<br>CTTAATG |
| covS-F1 | Primer used with covS-BamH-R to amplify arm 1 (892 bp upstream covS) | AGGATTTTTCTTAGTTTTACCCCTATAAA<br>CTGATC |
| covS-BamH-R | Primer used with covS-F1 to amplify arm 1 (892 bp upstream covS) and add a 3' BamHI site for insertion into pLT06 vector | CCTGCAGGTCGACTCTAGAGGATCCGT<br>GACAGAGGACACTACTC |
| covS-chF | Primer used with covS-chF to check ΔcovS mutation | TCAGCTTGTCGGCTATC |
| covS-chR | Primer used with covS-chR to check ΔcovS mutation | GTGTTATGACACGAGAAGAG |
| Abx1-EcoR-F | Primer used with Abx1-R1 to amplify arm 1 (957 bp downstream *abx1*) and add a 5' EcoRI site for insertion into pLT06 vector | GGAGAATTCATATTCTGTCCTTTGTCG |
| Abx1-R1 | Primer used with Abx1-EcoR-F to amplify arm 1 (957 bp upstream *abx1*) with short homology sequence to arm 2 | AGGATTTAGGATTATTTGAGTGTTGCAG<br>CTCAGTC |
| Abx1-F1 | Primer used with Abx1-BamH-R to amplify arm 2 (969 bp upstream *abx1*) with a short homology sequence to arm 1 | GCAACACTCAAATAATCCTAAATCCTTTT<br>TGTGTG |
| Abx1-BamH-R | Primer used with Abx1-F1 to amplify arm 2 (969 bp upstream *abx1*) and add a 3' BamHI site for insertion into pLT06 vector | GGAGGATCCATTGGCATTACCTGTTGC |
| Abx1-check-F | Primer used with Abx1-check-F to check Δ*abx1* mutation | ACCAGTACCAGTAGCACC |
| Abx1-check-R | Primer used with Abx1-check-R to check Δ*abx1* mutation | AAGCTTTTAACTTGTGCTTC |
| Abx1-EcoR-F | Primer used with Abx1-FLAG-R1 to amplify *abx1* and add a 5' EcoRI site for insertion into pLT06 vector | GGAGAATTCATATTCTGTCCTTTGTCG |
| Abx1-FLAG-R1 | Primer used with Abx1-EcoR-F to amplify *abx1* with an N-terminal 3×FLAG sequence | GGTGATTATAAAGATCATGATATCGATTA<br>CAAGGATGACGATGACAAGTGAGTGTT<br>GCAGCTCAGTC |
| Abx1-FLAG-F1 | Primer used with Abx1-3xF-BamH-R to amplify *abx1* with an N-terminal 3×FLAG sequence | CGATATCATGATCTTTATAATCACCGTCA<br>TGGTCTTTGTAGTCATCTGTTTCTTCAGT<br>CTTAATTAG |
| Abx1-3xF-BamH-R | Primer used with Abx1-FLAG-F1 to amplify *abx1* and add a 3' BamHI site for insertion into pLT06 vector | CCTGGATCCTATCACACAAAAAGGATTT<br>AGG |
| Abx1-3xF-chR | Primer used with Abx1-check-F to check *abx1*-3×FLAG fusion | TTAAGACTGAAGAAACAGATGAC |
| CaaX-FL-F | Primer used with Sp-CaaX-FL-R1 to amplify *cflP* and add a 5' BamHI for insertion into pLT06 vector | ATCCCCGGGTACCGAGCTCGAATTCGT<br>ATAAAGTTAGATAAAAGTGAG |
| Sp-CaaX-FL-R1 | Primer used with CaaX-FL-F to amplify *cfiP*-3×FLAG with short | ACTATTTTGGTCGTTACTTGTCATCGTCA<br>TCC |
|  | homology sequence to CaaX-FL-Sp-F1 |  |
| CaaX-FL-Sp-F1 | Primer used with CaaX-FL-Sp-R2 to amplify spectinomycin cassette with short homology sequence to CaaX-FL-Sp-R1 | CGATGACAAGTAACGACCAAAATAGTGA<br>GGAG |
| CaaX-FL-Sp-R2 | Primer used with CaaX-FL-Sp-F1 to amplify spectinomycin cassette with short homology sequence to CaaX-FL-Sp-F2 | TTTAGATTAAATCTAGATCCTTATAATTTT<br>TTAATCTG |
| Sp-CaaX-FL-F2 | Primer used with CaaX-FL-R to amplify 951 bp downstream *cflP*-3×FLAG with short homology sequence to Sp-CaaX-FL-R2 | TATAAGGATCTAGATTTAATCTAAATAAG<br>CTTCTCTTAC |
| CaaX-FL-R | Primer used with Sp-CaaX-FL-F2 to amplify 951 bp downstream *cflP*-3×FLAG and a 5' BamHI site for insertion into pLT06 vector | CCTGCAGGTCGACTCTAGAGGATCCTTT<br>TTCTACAACAACTTCCTG |
| CovR-Nco | Primer used to amplify *covR* and add a 5' NcoI site for insertion into pRSF-NT vector | AACCTTTACTTCCAGGGCGCCATGGGTA<br>AAAAGATCTTAATAATCG |
| CovR-Hind | Primer used to amplify *covR* and add a 3' HindIII site for insertion into pRSF-NT vector | TTAAGCATTATGCGGCCGCAAGCTTATT<br>TTTCACGAATCACATAGC |
| covS-Nco | Primer used to amplify *covS* and add a 5' NcoI site for insertion into pRSF-NT vector | AACCTTTACTTCCAGGGCGCCATGGCTC<br>GCGTCCAAGGAAGC |
| covS-Hind | Primer used to amplify *covS* and add a 3' HindIII site for insertion into pRSF-NT vector | TTAAGCATTATGCGGCCGCAAGCTTATT<br>CAAATTTTTTTATAGGAATGCG |
| Q-cflP-F | Primer used with Q-cflP-R to evaluate relative expression of *cflP* | ATGGTTAGTCGGGAGCGTTC |
| Q-cflP-R | Primer used with Q-cflP-F to evaluate relative expression of *cflP* | TGTACCTGCCATCCAACAGC |
| Q-covS-F | Primer used with Q-covS-R to evaluate relative expression of *covS* | AGAGTTCGTTTGTCGCAAGC |
| Q-covS-R | Primer used with Q-covS-F to evaluate relative expression of *covS* | GACTCGCCCGTTCTAACAATTC |
| Q-recA-F | Primer used with Q-recA-R to evaluate relative expression of *recA* | GTGGGATTGCTGCCTTTATTG |
| Q-recA-R | Primer used with Q-recA-F to evaluate relative expression of *recA* | CTGAGTCAGGTTGAGACAAGAG |
| Abi-EcoR | Primer used with FLAG-Abi-R to amplify sak_0054 and add a 5' EcoRI site for insertion into pDC123 vector | ATATCGATGCATATGAATTCTAATATTAG<br>GATAAACTAATATGG |
| FLAG-Abi-R | Primer used with Abi-EcoR to amplify sak_0054 with short homology sequence to Abi-FLAG-F | TCTTTGTAGTCATATCTAACAGATACTCT<br>CAC |
| Abi-FLAG-F | Primer used with Abi-Xho-R to amplify 3×FLAG tag from pDC123-*cfIP*-3×FLAG with short homology sequence to Abi-FLAG-R | ATCTGTTAGATATGACTACAAAGACCAT<br>GACG |
| Abi-Xho-R | Primer used with Abi-FLAG-F to amplify 3×FLAG tag from pDC123-*cfIP*-3×FLAG and add a 3' XhoI site for insertion into pDC123 vector | GGGGCAAAGACAAGCTCGAG |

**Table S7.** CDM final composition.

| Component | Final concentration |
| --- | --- |
| K<sub>2</sub>HPO<sub>4</sub> | 2.2 mM |
| KH<sub>2</sub>PO<sub>4</sub> | 14.69 mM |
| NaH<sub>2</sub>PO<sub>4</sub> H<sub>2</sub>O | 53.2 mM |
| Na<sub>2</sub>HPO<sub>4</sub> 7H<sub>2</sub>O | 195.4 mM |
| MgSO<sub>4</sub> | 28.4 mM (HighMg) or 7 mM (LowMg) |
| MnSO<sub>4</sub> | 0.236 mM |
| CH<sub>3</sub>COONa | 94.48 mM |
| NAHCO<sub>3</sub> | 59.52 mM |
| CaCl<sub>2</sub> | 91.8 µM |
| FeSO<sub>4</sub> | 35.8 µM |
| Fe(NO<sub>3</sub>)<sub>3</sub> 9H<sub>2</sub>O | 4.8 µM |
| Cysteine-HCl | 7.4028 mM |
| p-aminobenzoic acid | 2.8 µM |
| Biotin | 1.6 µM |
| Folic acid | 3.6 µM |
| Nicotinamide* | 0.25 µM |
| Pantothenate calcium salt | 8.4 µM |
| Pyridoxal | 11.8 µM |
| Pyridoxamine dihydrochloride | 8.2 µM |
| Riboflavin | 10.6 µM |
| Thiamine hydrochloride | 5.8 µM |
| Adenine | 296 µM |
| Guanine hydrochloride | 212 µM |
| Uracil | 356 µM |
| DL-alanine | 2.244 mM |
| L-arginine | 1.148 mM |
| L-aspartic acid | 1.5 mM |
| L-asparagine | 1.512 mM |
| L-cystine | 0.824 mM |
| L-glutamic acid | 1.356 mM |
| L-glutamine | 2.736 mM |
| Glycine | 2.664 mM |
| L-histidine | 1.288 mM |
| L-isoleucine | 1.524 mM |
| L-leucine | 1.524 mM |
| L-lysine | 1.368 mM |
| L-methionine | 1.34 mM |
| L-phenylalanine | 1.208 mM |
| L-proline | 1.736 mM |
| Hydroxi-L-proline | 1.736 mM |
| L-serine | 1.9 mM |
| L-threonine | 1.676 mM |
| L-tryptophan | 0.976 mM |
| L-tyrosine | 1.1 mM |
| L-valine | 1.704 mM |
| Glucose | 111.01 mM |

\*Nicotinamide was substituted with 3.84 μM of NAD for transposon screening

### Supplementary References

1. Zhang Y, Skolnick J. 2005. TM-align: a protein structure alignment algorithm based on the TM-score. *Nucleic Acids Res* 33:2302-9.
2. Lembo A, Gurney MA, Burnside K, Banerjee A, de los Reyes M, Connelly JE, Lin WJ, Jewell KA, Vo A, Renken CW, Doran KS, Rajagopal L. 2010. Regulation of CovR expression in Group B Streptococcus impacts blood-brain barrier penetration. *Mol Microbiol* 77:431-43.
3. Areschoug T, Stalhammar-Carlemalm M, Karlsson I, Lindahl G. 2002. Streptococcal beta protein has separate binding sites for human factor H and IgA-Fc. *J Biol Chem* 277:12642-8.
4. Spencer BL, Chatterjee A, Duerkop BA, Baker CJ, Doran KS. 2021. Complete Genome Sequence of Neonatal Clinical Group B Streptococcal Isolate CJB111. *Microbiol Resour Announc* 10.
5. Glaser P, Rusniok C, Buchrieser C, Chevalier F, Frangeul L, Msadek T, Zouine M, Couve E, Lalioui L, Poyart C, Trieu-Cuot P, Kunst F. 2002. Genome sequence of *Streptococcus agalactiae*, a pathogen causing invasive neonatal disease. *Mol Microbiol* 45:1499-513.
6. Chaffin DO, Rubens CE. 1998. Blue/white screening of recombinant plasmids in Gram-positive bacteria by interruption of alkaline phosphatase gene (*phoZ*) expression. *Gene* 219:91-9.
7. Hooven TA, Bonakdar M, Chamby AB, Ratner AJ. 2019. A Counterselectable Sucrose Sensitivity Marker Permits Efficient and Flexible Mutagenesis in *Streptococcus agalactiae*. *Appl Environ Microbiol* 85.
8. Thurlow LR, Thomas VC, Hancock LE. 2009. Capsular polysaccharide production in *Enterococcus faecalis* and contribution of CpsF to capsule serospecificity. *J Bacteriol* 191:6203-10.
9. Korotkov KV, Delarosa JR, Hol WGJ. 2013. A dodecameric ring-like structure of the N0 domain of the type II secretin from enterotoxigenic *Escherichia coli*. *J Struct Biol* 183:354-362.
